# A Competitive Framework for Modeling EEG Microstate Durations

**DOI:** 10.64898/2026.05.20.726605

**Authors:** Carlos M. Gómez, Brenda Y. Angulo

## Abstract

**Background:** This study examines a competition-based model (C-model) designed to capture the temporal dynamics of successive brain microstates derived from electroencephalography (EEG) recordings during eyes-open conditions. The analyzed data were obtained from a public repository comprising microstate sequences from 60 sessions of a single subject [1]. When applied to microstate dynamics, the C-model posits a stochastic competition among neural circuits underlying the expression of individual microstates.

**Methods:** The model is formulated at a conceptual level (computational level in Marr’s framework) and employs a geometric distribution to account for the long right tail of microstate duration distributions, interpreted as the probability of “failure” of the currently active microstate to persist. To account for the short-lived left tail, the model incorporates a transient increase in the stability of the currently active network, or equivalently, a temporary decrease in the activation probability of competing microstates (refractory period).

**Results:** The model provides a good fit to the microstate duration distributions across all 60 sessions. One third of sessions showed microstate identity sequential dependency with respect to the previous microstates.

**Discussion:** These results suggest that the C-model captures key aspects of microstate temporal structure. Moreover, because microstate probabilities can be modulated by psychophysiological conditions—including the influence of previously active networks—the model may serve as a building block for more comprehensive neurobiological frameworks of neural and behavioral dynamics. In such frameworks, microstate sequences could emerge from structured competition and flow among neural networks supporting microstate expression.

## Introduction

The present study aims to model the duration of EEG microstates observed during spontaneous resting-state activity. The proposed approach builds on previous work in which diverse processes have been modeled as emerging from competition among neural networks subserving mutually incompatible behaviors. In its simplest form, this framework accounts for the duration of a given behavioral or neural state using a geometric distribution. However, deviations from this distribution are typically observed for short event durations, where the currently expressed state exhibits a transient advantage over competing alternatives.

Within this framework, the competition-based model (C-model) can be understood as a winner-take-all architecture [2] that incorporates a refractory period (see Methods). During this period, the currently active state retains a temporary dominance, increasing its probability of persistence immediately after a transition. This approach has been successfully applied across multiple domains. For example, lever-pressing behavior in rats under variable-interval reinforcement schedules shows inter-press intervals consistent with a geometric distribution, alongside a refractory period linked to subsequent responses [3]. Similarly, the durations of bistable perception in ambiguous figures—such as the Necker cube or binocular rivalry—have been described within this framework [4], and competition between saccadic and fixation-related networks has also been proposed [5]. More recently, the durations of immobility and activity periods in *Drosophila melanogaster* during free exploration have also been successfully modeled using the C-model [6].

The core assumption of the C-model is that a perceptual, behavioral, or physiological state is expressed when the activity of its underlying neural network exceeds that of all competing networks. The model further assumes the existence of a refractory period, during which the probability that an alternative network overtakes the currently dominant one is governed by a sigmoid function. Once the asymptotic value of this function is reached, competition becomes effectively unbiased. The model is fundamentally probabilistic as the dynamics of the underlying networks. It can accommodate modulation by time-dependent factors such as attention, motivation, behavioral sequences or rhythmic brain states, which may influence the probability of state expression.

EEG microstates are defined as quasi-stable periods of scalp electrical topography [7,8]. Microstate parameters have been shown to differentiate cognitive states as well as clinical populations [9]. From a more theoretical perspective, microstates have been described as the “atoms of thought,” suggesting that their sequential organization may form a syntax-like structure underlying brain function.

In this context, we test whether the temporal durations of EEG microstates can be accounted for by the C-model. Additionally, given prior evidence that microstate sequences are not entirely random but exhibit preferred transitions, we assess whether transition probabilities between consecutive states deviate from randomness. If such sequential structure is confirmed, it would support the notion of a constrained flow among microstates, requiring the incorporation of structured transition dynamics into the C-model framework.

## Methods

The data analyzed in the present study were obtained from a publicly available dataset [1], accessible at https://doi.org/10.6084/m9.figshare.24877770.v3. The analysis scripts, provided in the Supplementary Material, were designed to operate on data organized in the *ALLEEG* structure. This dataset comprises categorized microstate sequences from a single subject, recorded across 60 resting-state sessions with eyes open over a three-month period.

Preprocessing of the EEG data included line-noise removal using the *cleanLineNoise*.*m* function, re-referencing to the average reference, and band-pass filtering in the 2–30 Hz range. Independent component analysis (ICA) was subsequently applied for further denoising. Microstate segmentation was performed using the EEGLAB Toolbox [10]. Each EEG time point was assigned to one of four prototypical microstate classes, yielding a symbolic microstate sequence representation of the signal. Microstates with durations shorter than 30 ms were excluded, as they were considered insufficiently stable, following the criteria proposed by the original data contributors.

The dataset field *ALLEEG(session)*.*microstate*.*fit*.*labels* from the aforementioned repository contains the classification of each 2 ms time point into one of the prototypical microstates. Custom preprocessing scripts enabled the computation of microstate durations, defined as the time intervals between successive microstate transitions. Additional methodological details can be found in the original publication [1].

The main working hypothesis of this study is that the duration of independent microstates can be adequately described by a geometric distribution, in which the probability (1-p) is the probability that a microstate (A,B,C or D) have a higher activity than any other microstate in the current sampling period. For each microstate and session (1-p) is estimated by the C-model, see below. This hypothesis is grounded in prior evidence showing that geometric distributions effectively characterize transitions between incompatible behavioral states and bistable perceptual phenomena, as outlined in the Introduction. For the analysis of the microstate data, the geometric model is used to evaluate the duration of the four characterized microstates assumed to arise from competition between mutually exclusive neural stochastic processes.

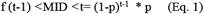

This equation implies that the probability for a microstate (e.g. microstate A) to have a duration (MID) taking a value between *(t-1)* and *t (*f (t-1) *<*MID *<*t;) *t* measured from the preceding microstate transition (*t*=0), depends on the probability *(1-p)* that the neural network inducing A exhibits higher activation than the networks promoting any other alternative (e.g. B,C,D) microstate (p: probability than any other alternative microstate gets a higher activity) (Fig. 1A). A similar relationship holds for the duration of any other microstate. This formulation reflects a memoryless process in which the probability of transition remains constant over time. However, empirical microstate duration distributions exhibit a monomodal right-skewed, long-tailed profile, and the geometric distribution fails to adequately capture the short-duration regime. The C-model approaches this asymmetry in the distributions by considering that once a network wins competition with the other networks, it has a much higher probability (1-p’(t)) to continue being prevalent, showing a dynamics that makes its probability to be function of the time elapsed from the previous transition (Gómez et al., 2025). This is obtained by modelling p’(t) as a sigmoid from the origin (Fig. 1B).

**Figure 1.**
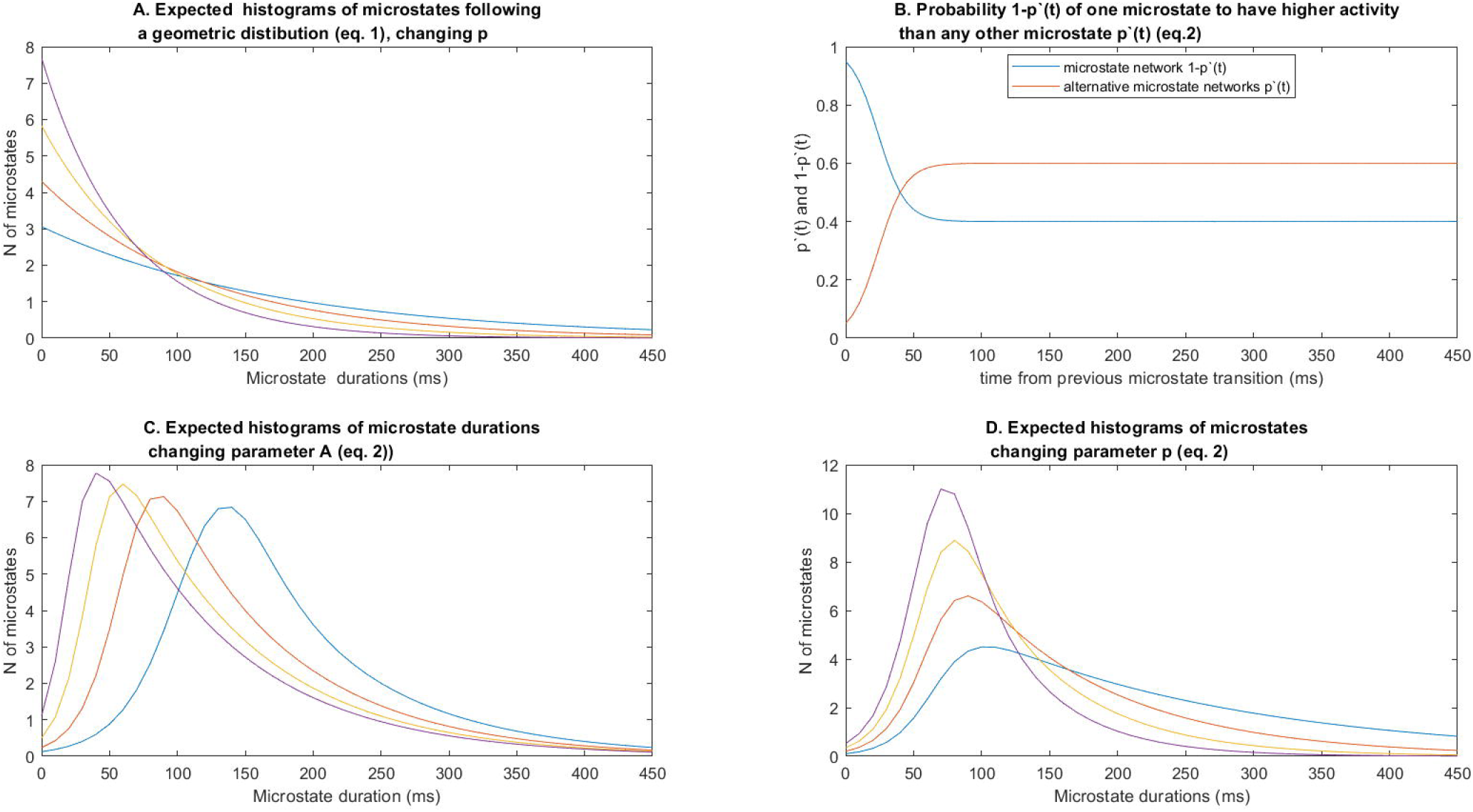
Competition model. (A) Frequency histogram of a geometric distribution as defined by Equation (1). (B) Temporal evolution of 1-p’(t) representing the probability that the neural network underlying a given microstate (e.g., microstate A) is expressed. Here p’(t) denotes the summed probability of expression of the alternative microstates (Equation 2). (C) Frequency histogram of microstate duration times computed from Equation (3), obtained by modulating the parameter A of the geometric distribution. (D) Same as in (C), but varying the asymptotic values of p’ (t) while keeping parameter A constant.

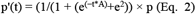

A: parameter to modulate the curvature of the sigmoid.

e^2^ is introduced into the sigmoid equation to have the origin at time zero (time of previous microstate transition).

p: Asymptotic value for the alternative microstate (e.g. B,C,D) to present higher activity than the currently expressed microstate (e.g. A).

The time evolution of p’ (t) and (1-p’(t)) is shown in Fig. 2B. Finally the probability f would be:

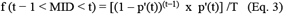

**Figure 2.**
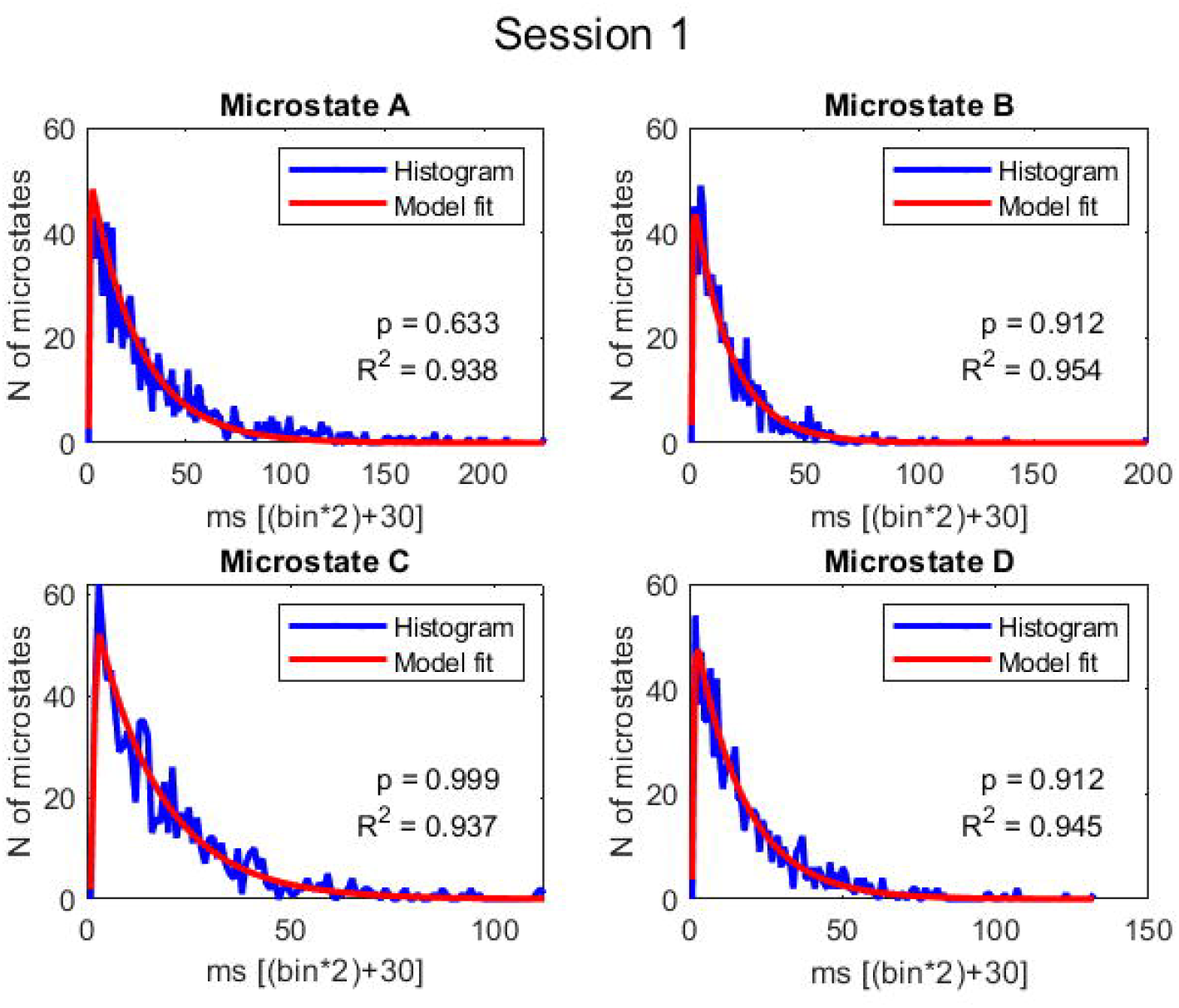
C-Model adjusting. Frequency histograms and C-model adjusted. The Kolmogorov_Smirnov p-value, and the R^2^ between the c-model and the histogram are also represented. The microstates durations are expressed as a function on the number of bins, and adding 30ms as indicated in methods.

The term T is introduced to normalize Equation (3), so that the area under the curve would approximate to 1. T is not computed analytically but numerically, by summing the area under the curve created by the numerator of Equation (3).

By changing systematically the curvature parameters of the sigmoid (A), and p values, and correlating the results between the empirical histogram and the results of equation 3, the optimal A and p parameters are estimated. The effects of the parameters A and p for changing the shape of microstates frequency distribution computed from equation 3, are displayed in Figs. 1C and 1D. The function *comp_model*.*m* used for estimation of A and p is enclosed in the supplementary material.

Once the parameters A and p have been estimated, the level of significance of the fit between the empirical distribution of microstates and the C-model were estimated by means of the Kolmogorov–Smirnoff goodness-of-fit test, and by displaying the R^2^ of the estimated model vs. the empirical histograms [11].

Apart from the refractory period indexed by parameter A, there are at least four possible departures from unbiased competition among the neural networks underlying the different microstates: (i) non-random sequential dependencies between microstates; (ii) sequential dependencies in the durations of successive microstates (i.e., periods of relatively short or long durations); (iii) influences of internal states, such as attention, motivation, or brain rhythms at different scales; and (iv) reinforcement contingencies. We focus here on the first two possibilities. To assess potential non-random sequential dependencies between microstates, a chi-squared test was applied to each microstate. This test evaluated whether transitions to alternative microstates occurred randomly or, instead, exhibited a preferred directionality. No further analyses were conducted to determine the specific direction of these transitions, as this aspect of microstate syntax has been addressed in greater detail elsewhere [12,13].

To examine sequential duration dependencies between successive microstates, the autocorrelation function (ACF) was computed. To minimize the potential influence of differences in session or microstate durations, z-scores were calculated for each microstate within each session and then reordered to match the original temporal sequence. The statistical significance of the observed effects was assessed by comparison with surrogate datasets generated from the same recordings. These surrogates were created using a random temporal shuffling procedure. For each dataset, the original interleaved time series were randomly permuted 1,000 times. This approach preserves the marginal distribution of values while eliminating all temporal dependencies, thereby providing a null hypothesis of no serial correlation. Significance thresholds for temporal dependencies in microstates were corrected using the False Discovery Rate method [14].

## Results

Figure 2 shows the frequency histograms of the durations of the four canonical microstates for the subject in one session, together with the overlaid C-model fit. A significant goodness of fit, along with high R^2^ values obtained from regressing the frequency histograms on the fitted C-model, was consistently observed across all the remaining 59 sessions for the four canonical microstates (see Supplementary Figure 1). Figure 3 illustrates the superposition of the fitted C-model across all 60 sessions, revealing a high degree of similarity in the distribution shapes generated by the model.

**Figure 3.**
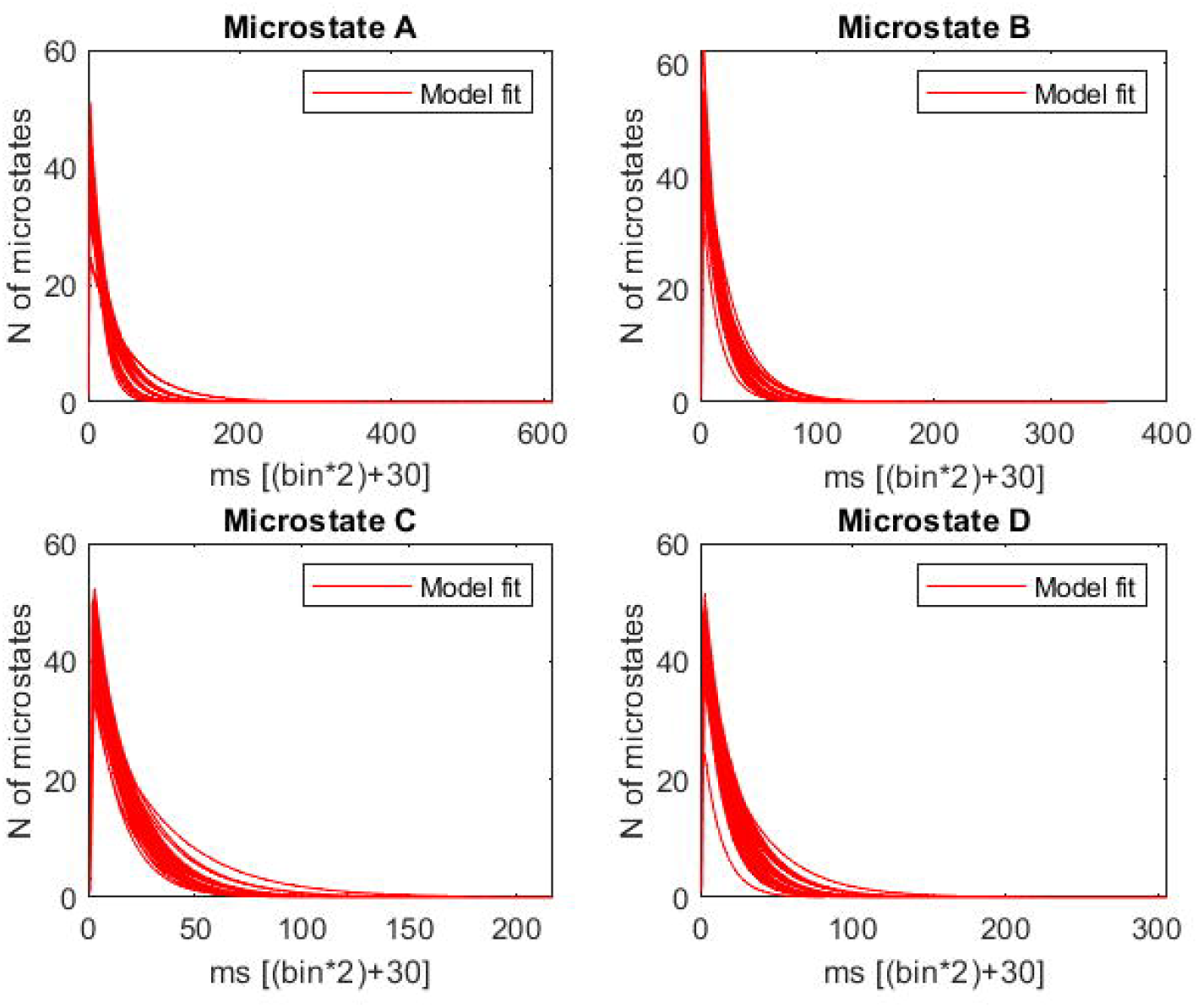
C-model shape across sessions. Overlay of the histograms representing the frequency distributions of the four microstates generated by the C-model for the 60 recorded sessions. Note the high similarity in shape across different sessions.

To test whether transitions between microstates occurred randomly, a chi-squared test was performed. The expected number of transitions—assuming randomness—was computed based on the relative frequency of each microstate divided by the total number of transitions, and compared with the observed transition counts.

Supplementary table 1 summarizes, for each microstate, the total number of sessions showing non-random transitions, along with the corresponding p-values from the chi-squared tests. As shown in Supplementary table 1, both random and non-random transition patterns varied across sessions.

To examine potential temporal dependencies in microstate durations across sessions, the autocorrelation function (ACF) was computed on the z-scores of the continuous series of microstate durations. z-score normalization was applied to minimize the influence of differences in mean duration across microstates. The ACF results for all 60 sessions are presented in Supplementary Figure 2. A total of 35 out of 60 sessions showed significant temporal dependencies. However, as illustrated in Suppl. Figure 2, no consistent temporal pattern emerged, and significant effects appeared at different lag bins (which in fact do not correspond to strict time lags, as the analyzed series represents successive microstate durations). Therefore no consistent time durations dependence was obtained.

## Discussion

The present report shows that the C-model provides a good fit to data from 60 sessions of a single subject recorded under resting-state conditions, which are assumed to reflect spontaneous neural dynamics, or at least activity unrelated to specific task demands. According to the assumptions of the model, a competitive interaction is established among the networks underlying the different microstates. In addition, a refractory period is incorporated and modeled by a sigmoid function, whereby the probability that a non-dominat network becomes expressed increases as a function of time since the previous microstate transition. As noted in the Introduction, this model has previously been successful in describing mutually incompatible behaviors in flies, rats, and humans [3,4,5,6].

Several models have been proposed to explain microstate duration distributions, each grounded in different theoretical frameworks [15,16]. However, none of these models explicitly incorporates the type of refractory period modeled here, which likely reflects a form of lateral inhibition among competing neural networks that gradually decays over time. The geometric distribution is typically considered the discrete analogue of the exponential distribution, where (1 − p) represents the probability of failure before the event occurs with probability p. In the present framework, however, the interpretation differs: (1 − p) reflects the persistence of the currently active microstate network, while p represents the probability that an alternative microstate network becomes active. The sigmoid modulation of p provides a mechanism ensuring that the hypothesized physiological function of a microstate can be sustained for a sufficient duration.

A wide range of factors—including cognitive and behavioral processes, age, and clinical conditions—may influence the probability of microstate expression. Sequential dependency has been one of the most extensively studied aspects, as it may reveal underlying “syntactic” structures of brain activity [12,13]. In the present report, we examined only pairwise (two-microstate) dependencies and found that approximately one-third of the cases (across 60 sessions and four microstates) could be classified as non-random. This suggests that transition probabilities are, at least in part, influenced by preceding microstates during resting state. Interestingly, the same subject exhibited varying levels of dependency across sessions (Supplementary Table 1), indicating that cognitive state, experimental conditions, physiological factors, and other variables may modulate sequential structure. In contrast, the possibility that specific recording periods systematically influence microstate duration, in a systematic manner, independently of microstate class was not supported by the autocorrelation analysis. The presence of sequential dependency suggests that the C-model should ultimately be integrated into a more comprehensive framework that incorporates contextual and organism-level influences on temporal structure, in some sort of underlying neural networks flow of biased transition probabilities [12,13].

The main limitation of the present study lies in the fact that the model was tested on a single subject, albeit across 60 sessions. A more complete evaluation would require replication in a larger sample.

## Data and code availability

The data analyzed in the present study were obtained from a publicly available dataset [1], accessible at https://doi.org/10.6084/m9.figshare.24877770.v3

The code used for generate the model and analyze the data can be obtained from the Supplementary material

## Author information Authors Affiliation

## Contributions

C.M.G . conceived the study, wrote the simulation code and main manuscript text.

B.Y Angulo prepared all figures, Supplementary figures and supplementary table, and participated in data analysis and editing the manuscript.

