## Supplementary figures, table and code for "A Competitive Framework for Modeling EEG Microstate Durations"

#### Supplementary material:

**Supplementary Figure 1. C-Model adjusting.** Frequency histograms and c-model adjusted. The Kolmogorov_Smirnov p-value, and the $R^2$ between the c-model and the histogram are also represented. Sessions 2-60 are displayed. The microstates durations are expressed as a function on the number of bins, and adding 30ms as indicated in methods.

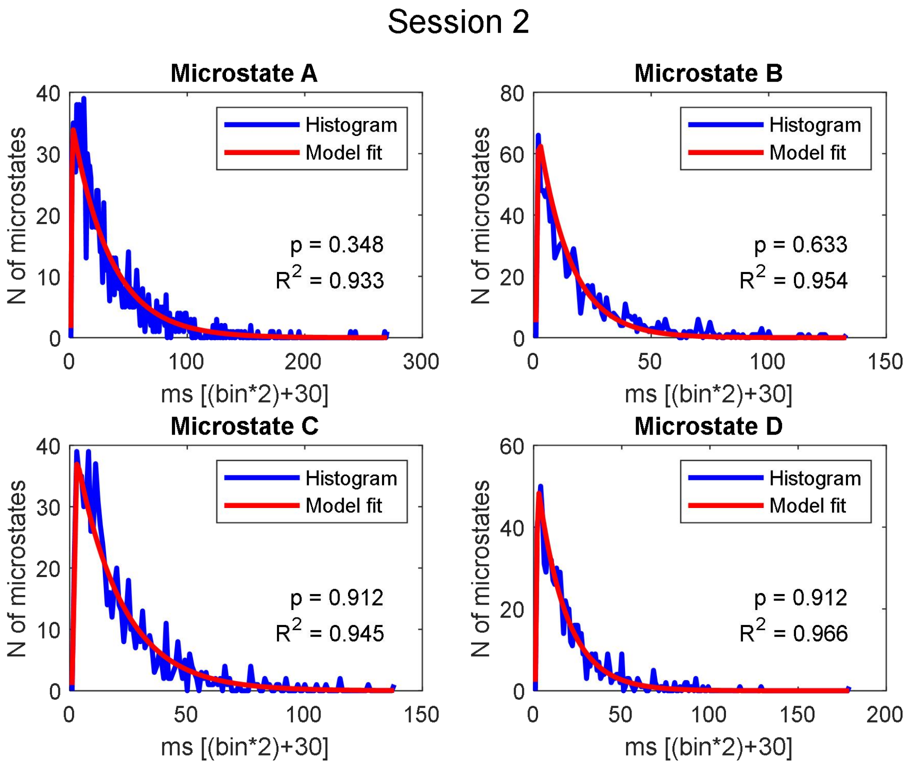

#### Session 3

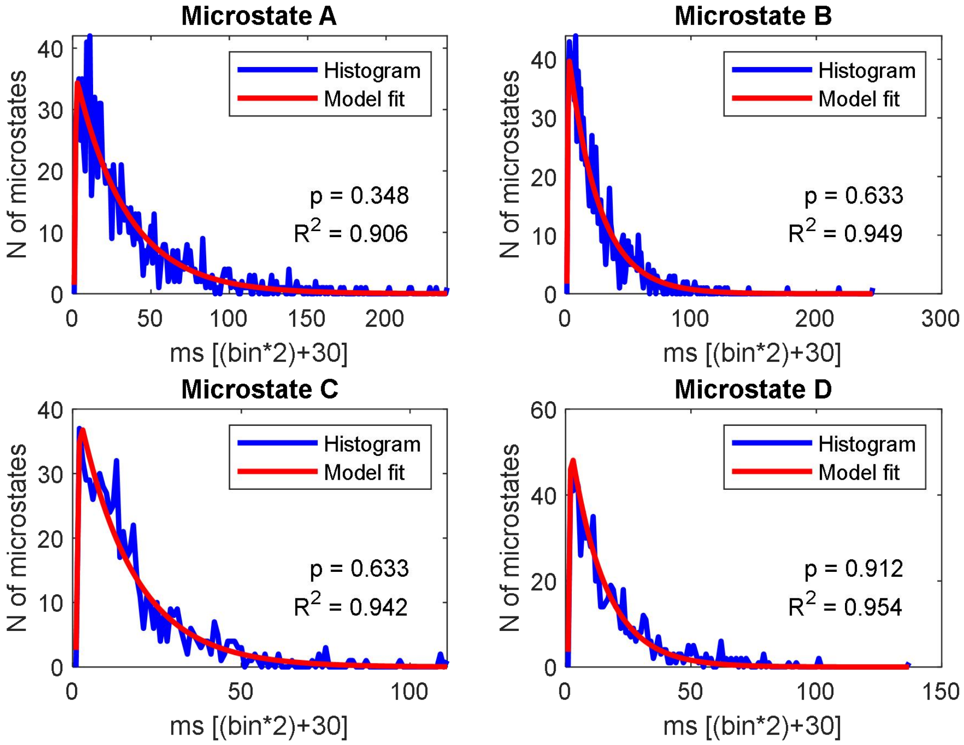


#### Session 4

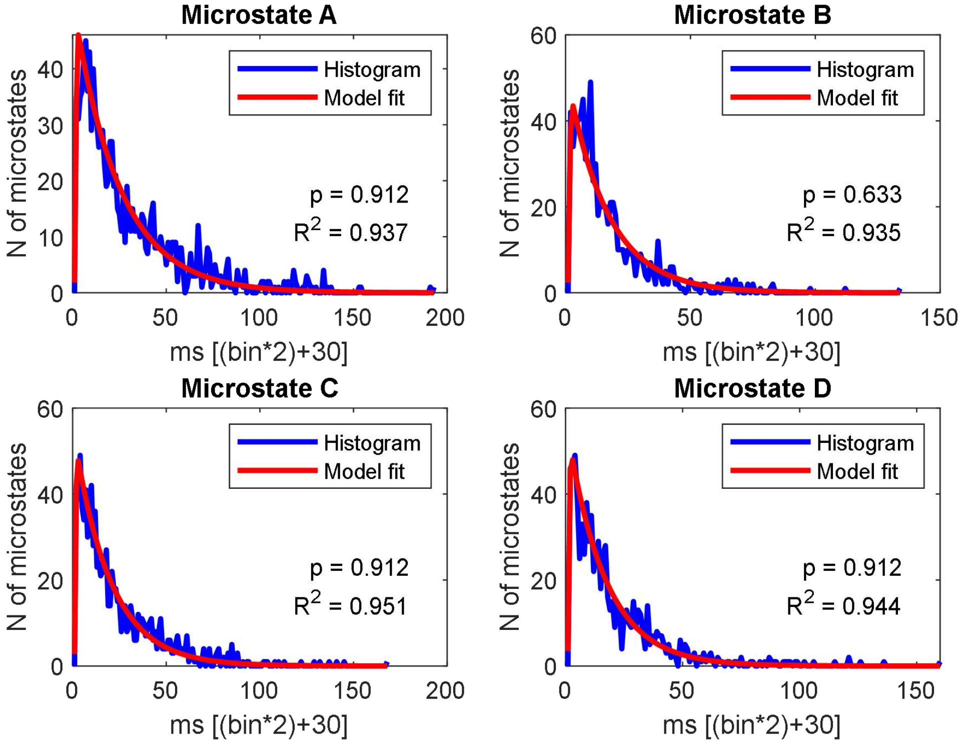

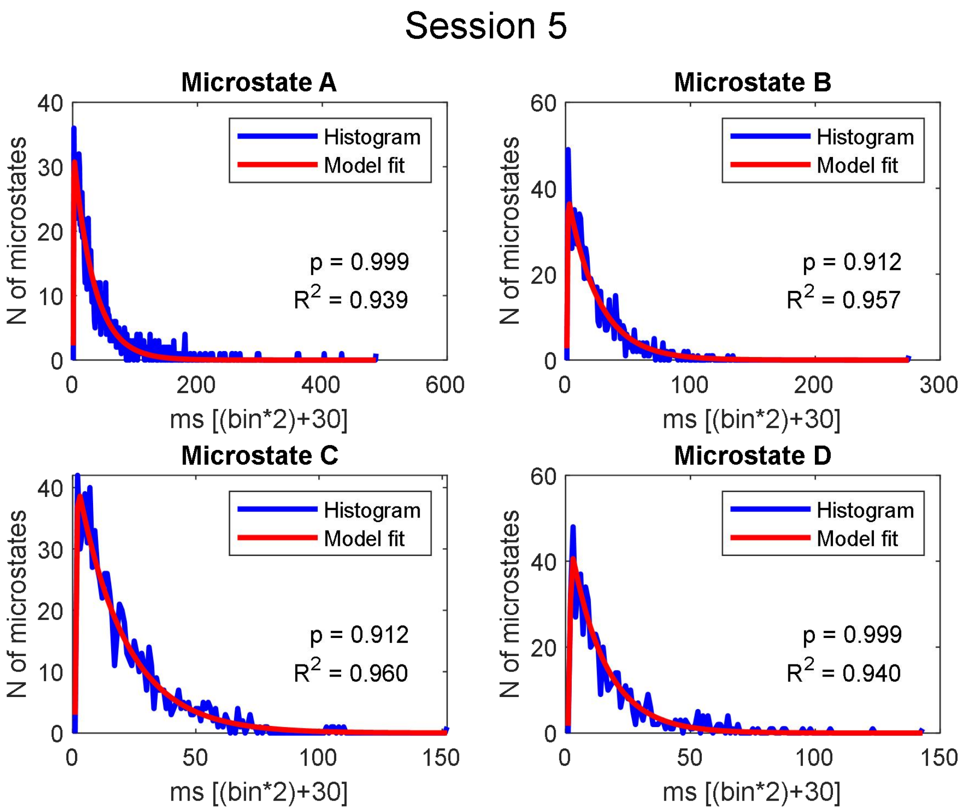
Session 5
Microstate A
Histogram
Model fit
p = 0.999
R2 = 0.939
N of microstates
ms [(bin*2)+30]
Microstate B
Histogram
Model fit
p = 0.912
R2 = 0.957
N of microstates
ms [(bin*2)+30]
Microstate C
Histogram
Model fit
p = 0.912
R2 = 0.960
N of microstates
ms [(bin*2)+30]
Microstate D
Histogram
Model fit
p = 0.999
R2 = 0.940
N of microstates
ms [(bin*2)+30]

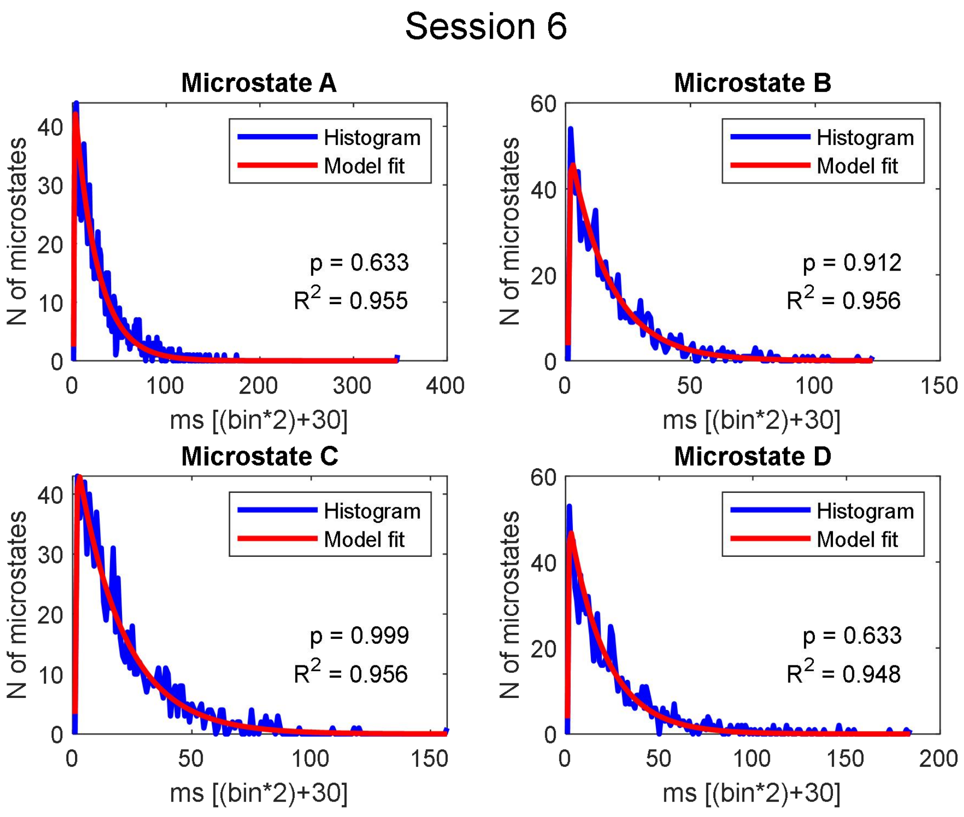
Session 6
Microstate A
Histogram
Model fit
p = 0.633
R2 = 0.955
N of microstates
ms [(bin*2)+30]
Microstate B
Histogram
Model fit
p = 0.912
R2 = 0.956
N of microstates
ms [(bin*2)+30]
Microstate C
Histogram
Model fit
p = 0.999
R2 = 0.956
N of microstates
ms [(bin*2)+30]
Microstate D
Histogram
Model fit
p = 0.633
R2 = 0.948
N of microstates
ms [(bin*2)+30]

Session 7

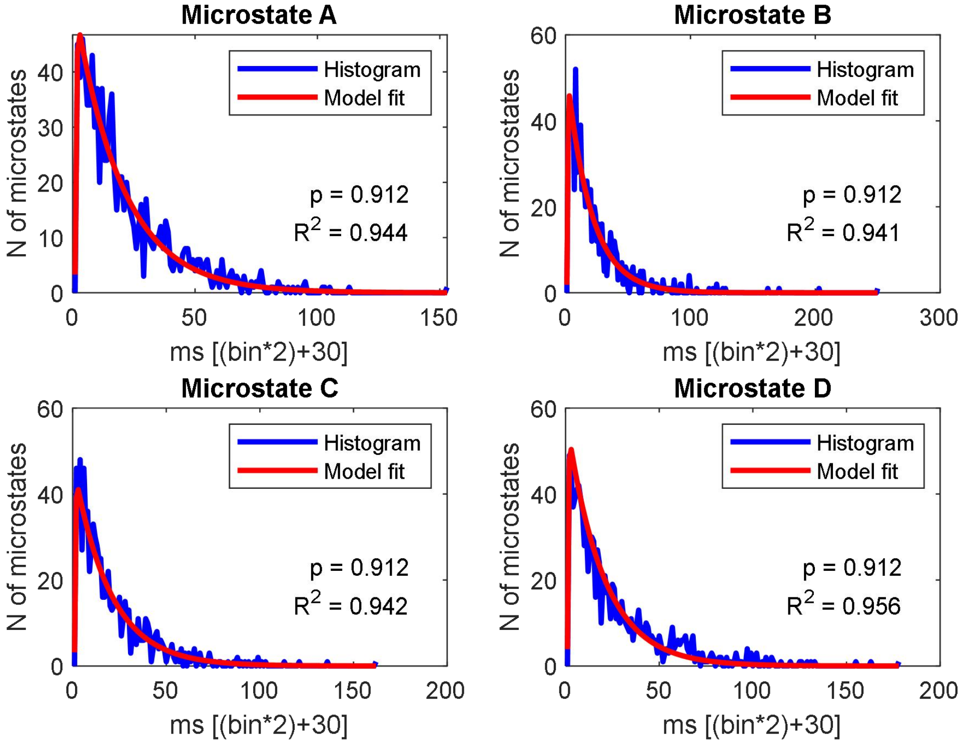


Session 8

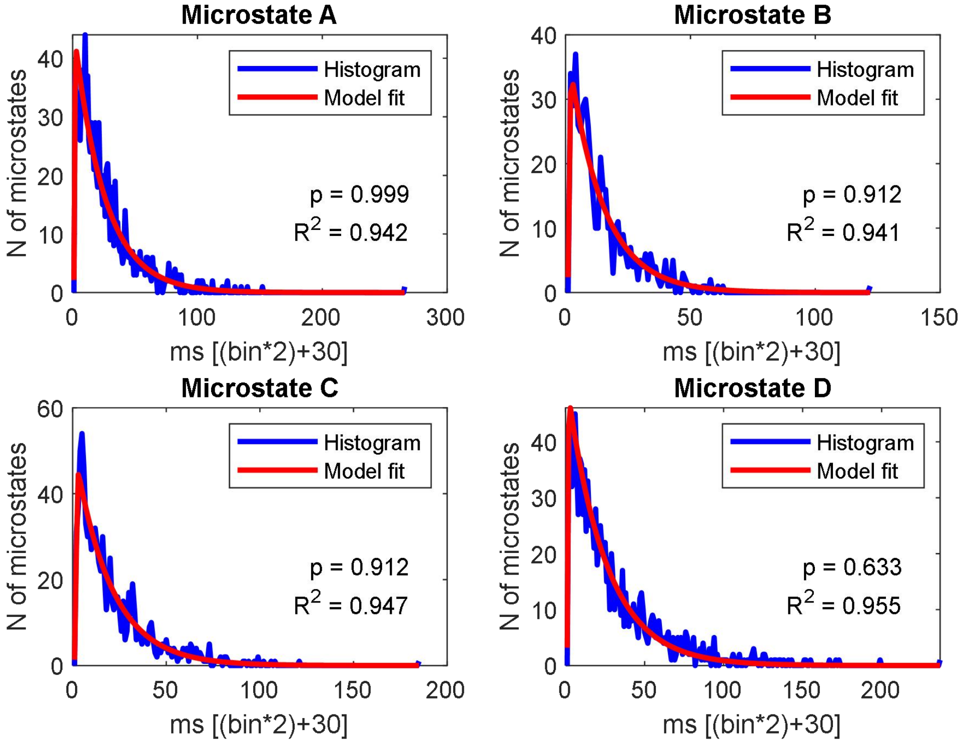

#### Session 9

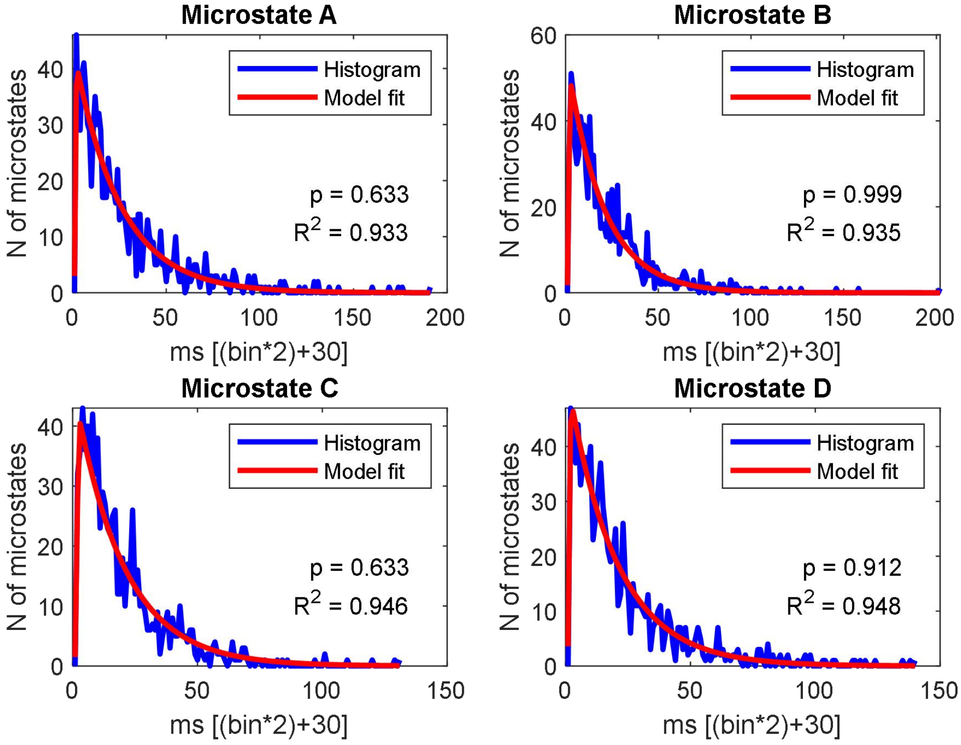


#### Session 10

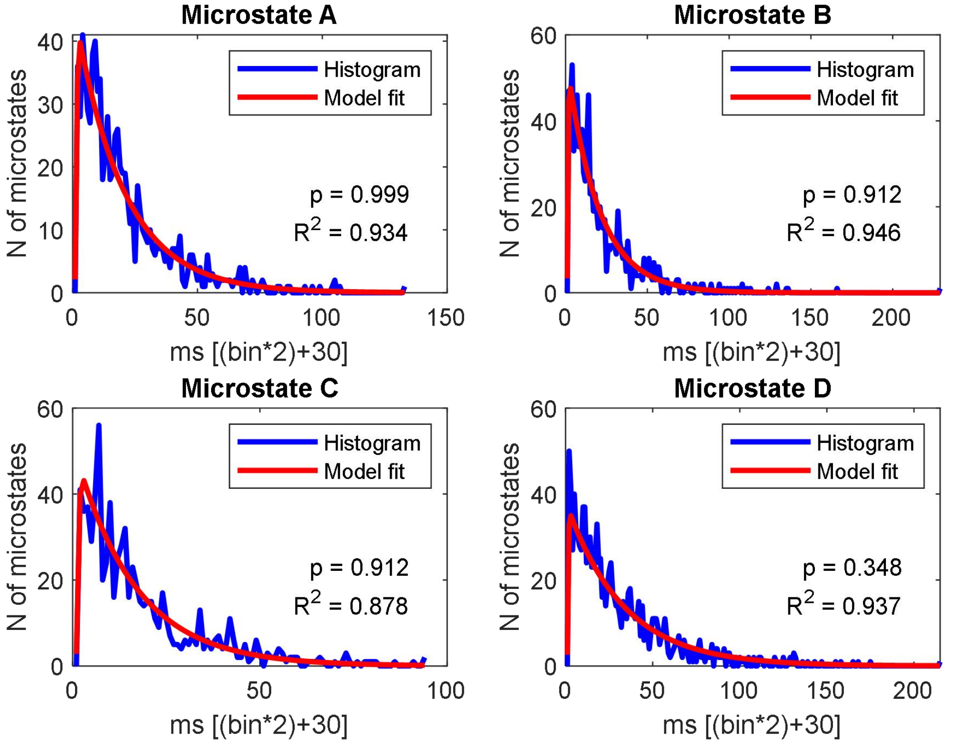

#### Session 11

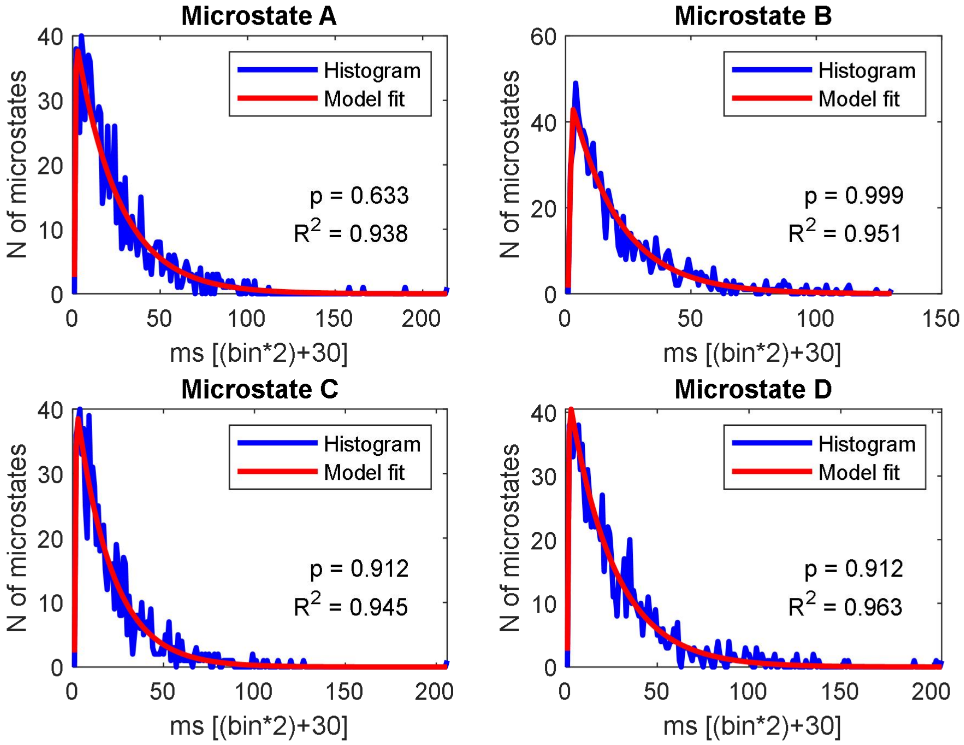


#### Session 12

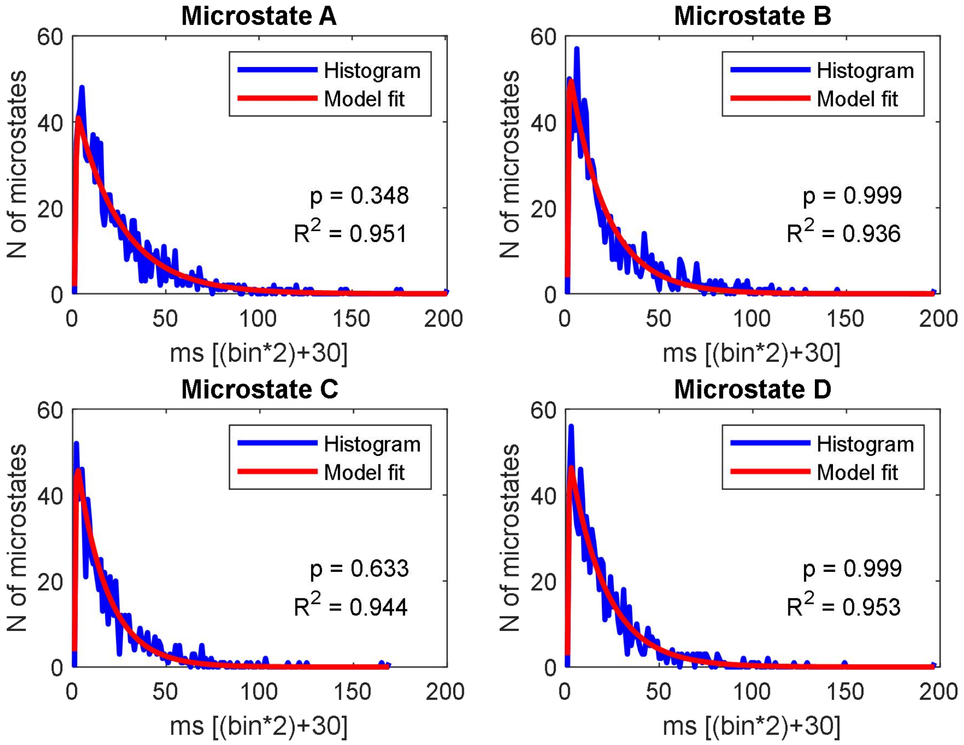

#### Session 13

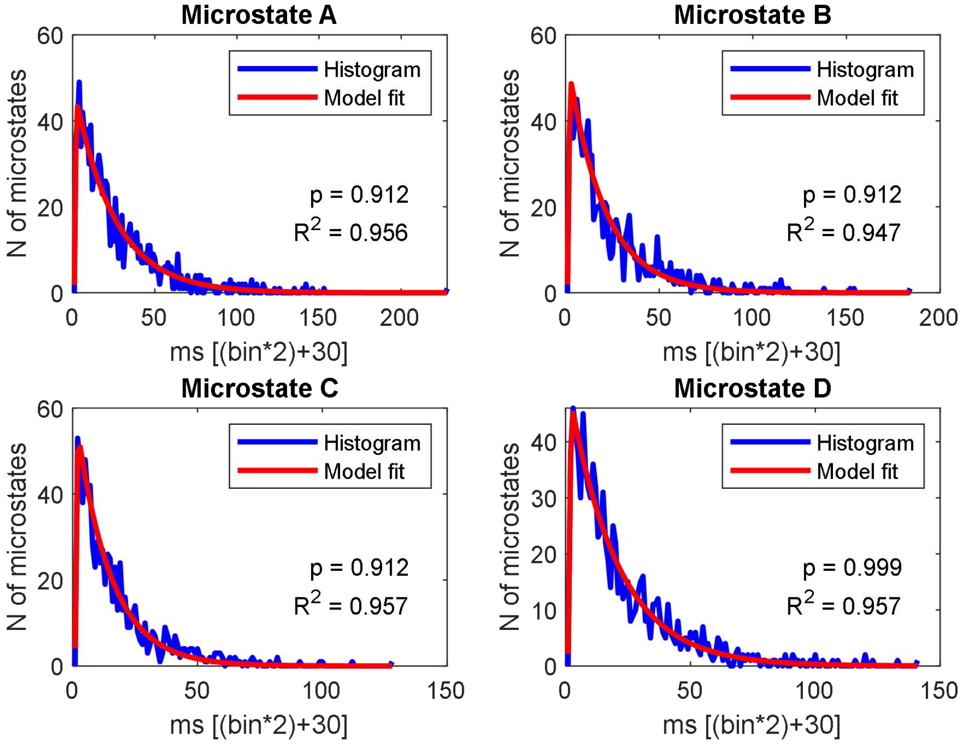


#### Session 14

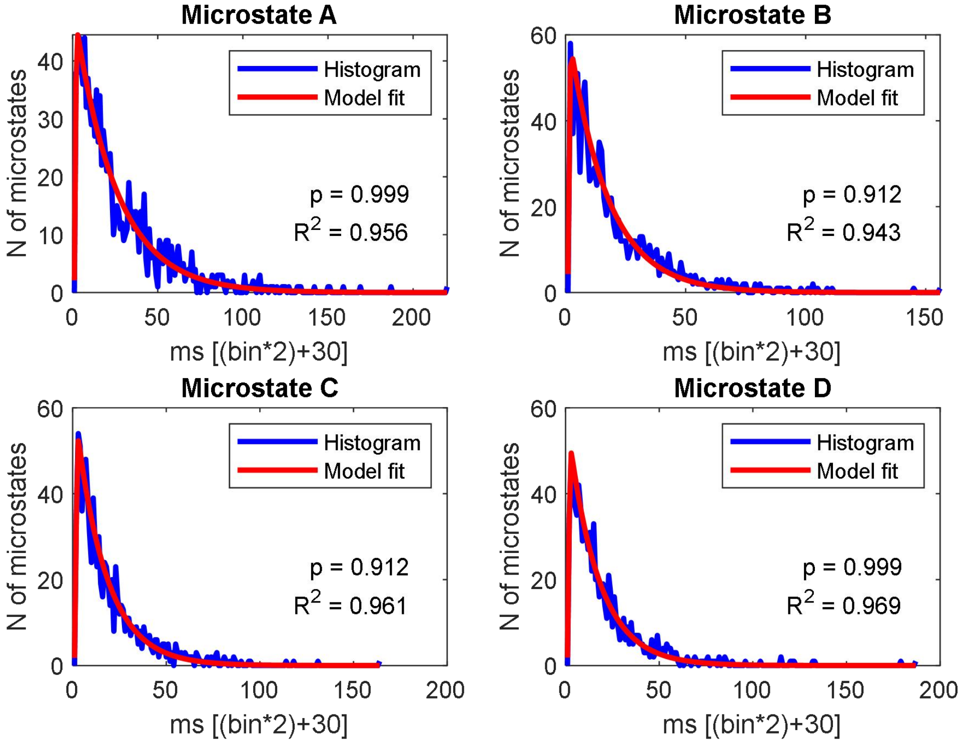

Session 15

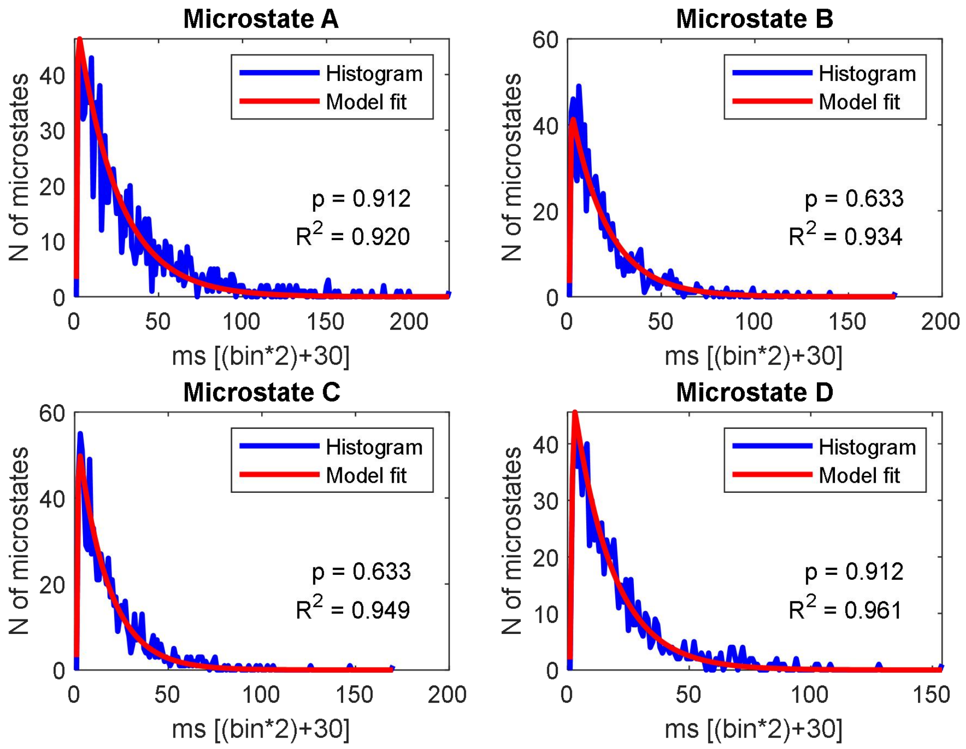


Session 16

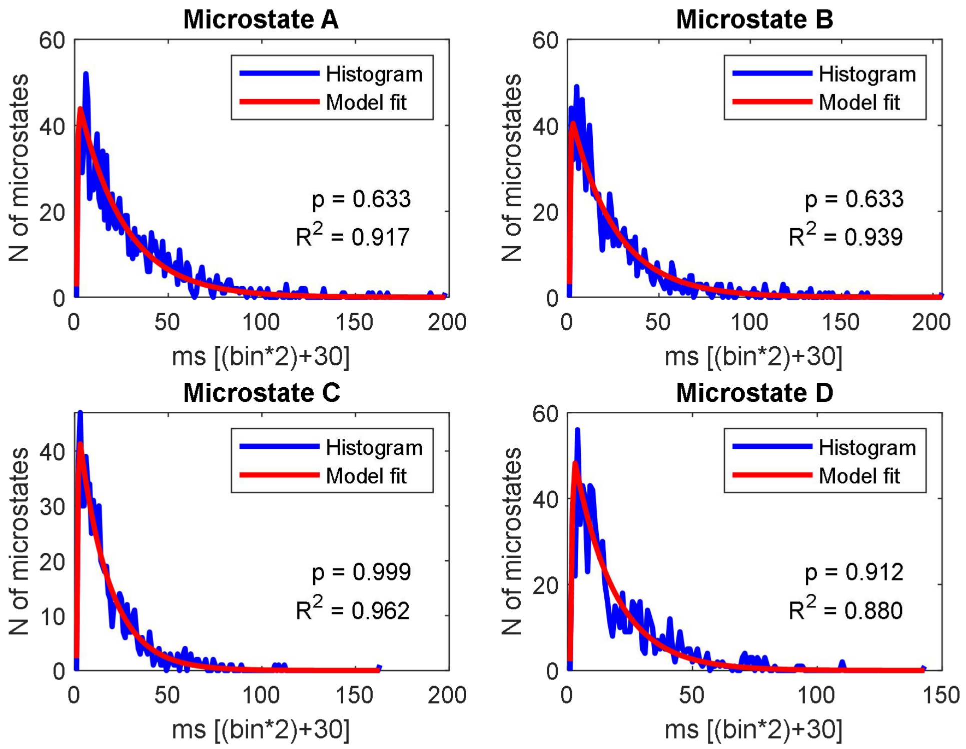

#### Session 17

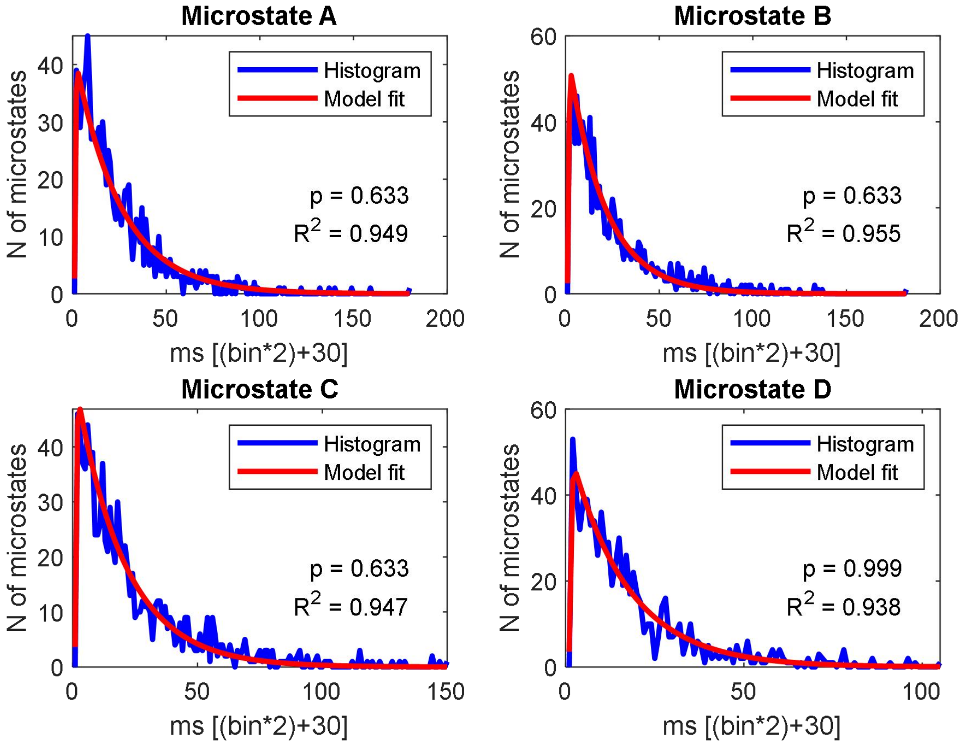


#### Session 18

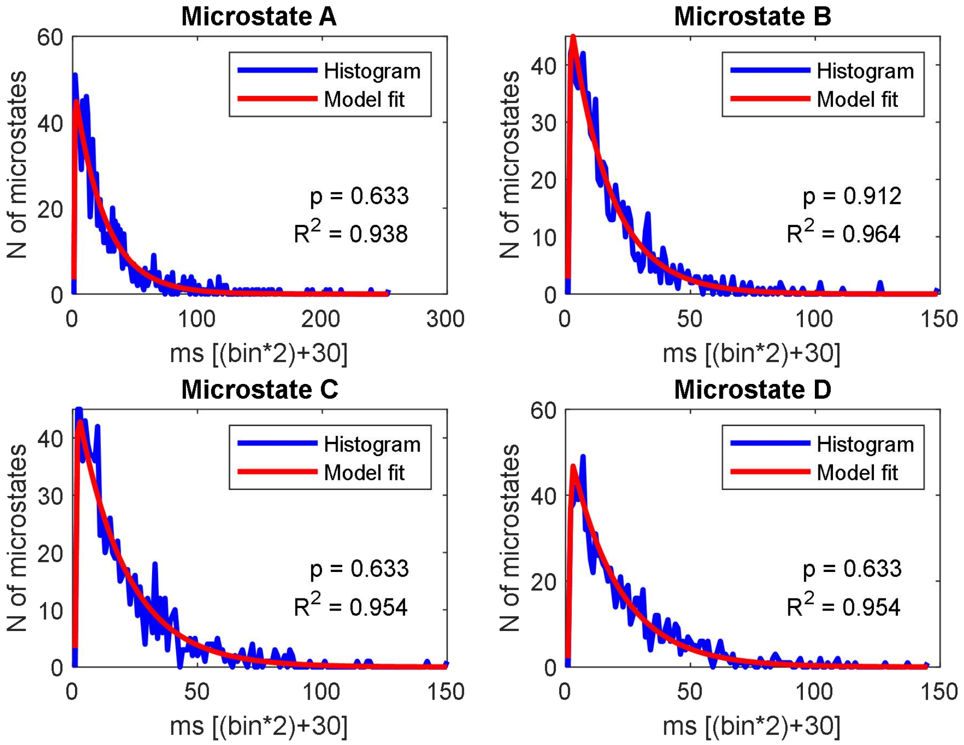

#### Session 19

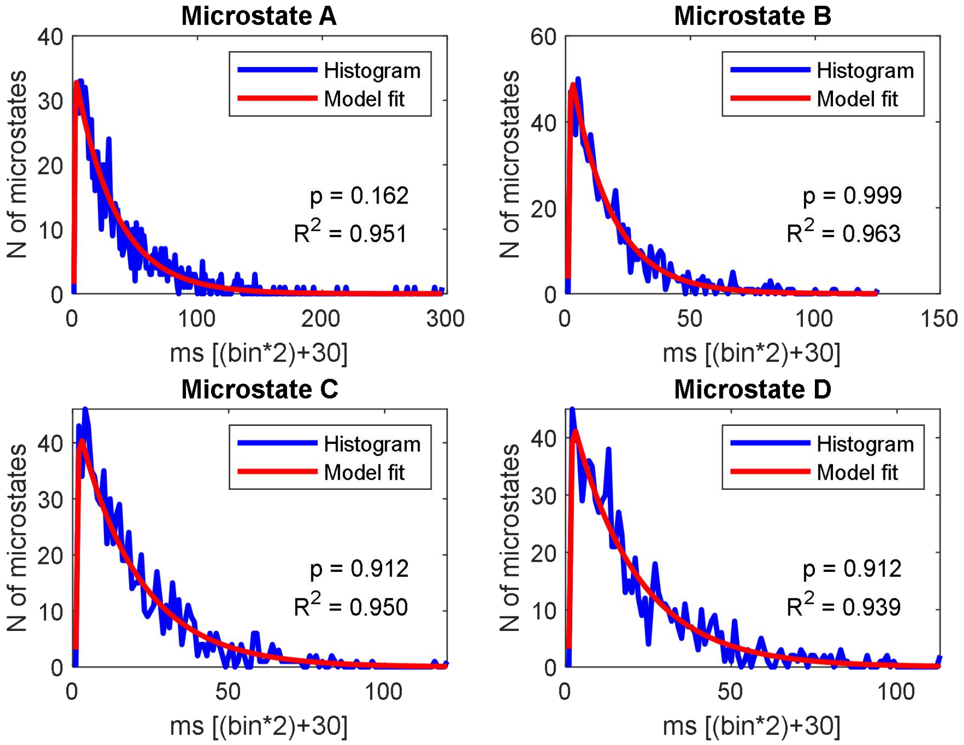


#### Session 20

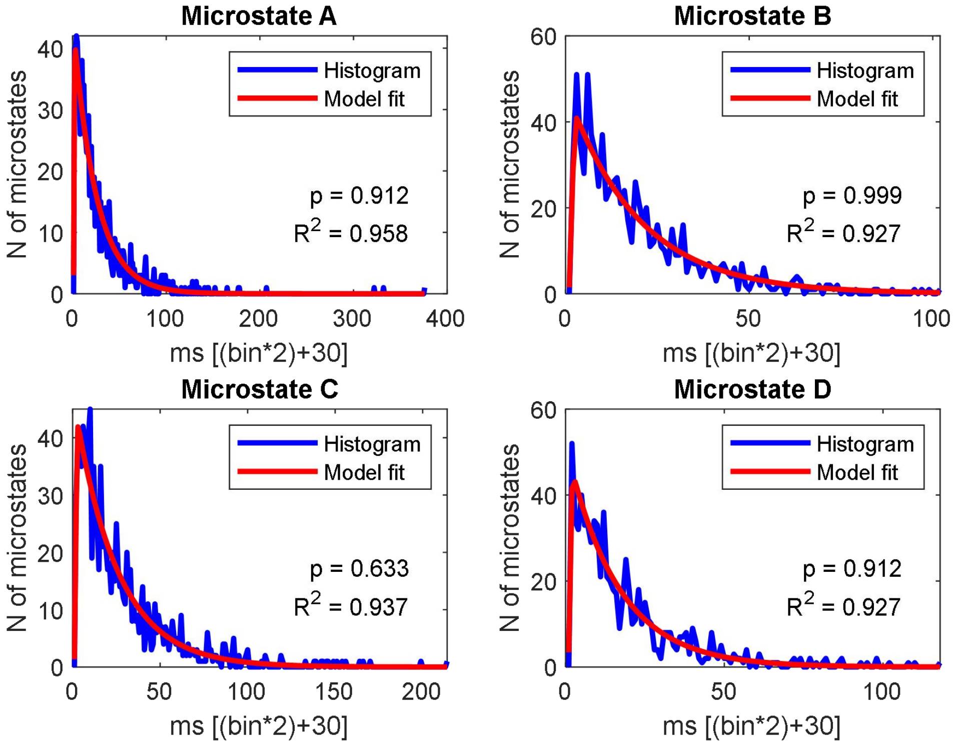

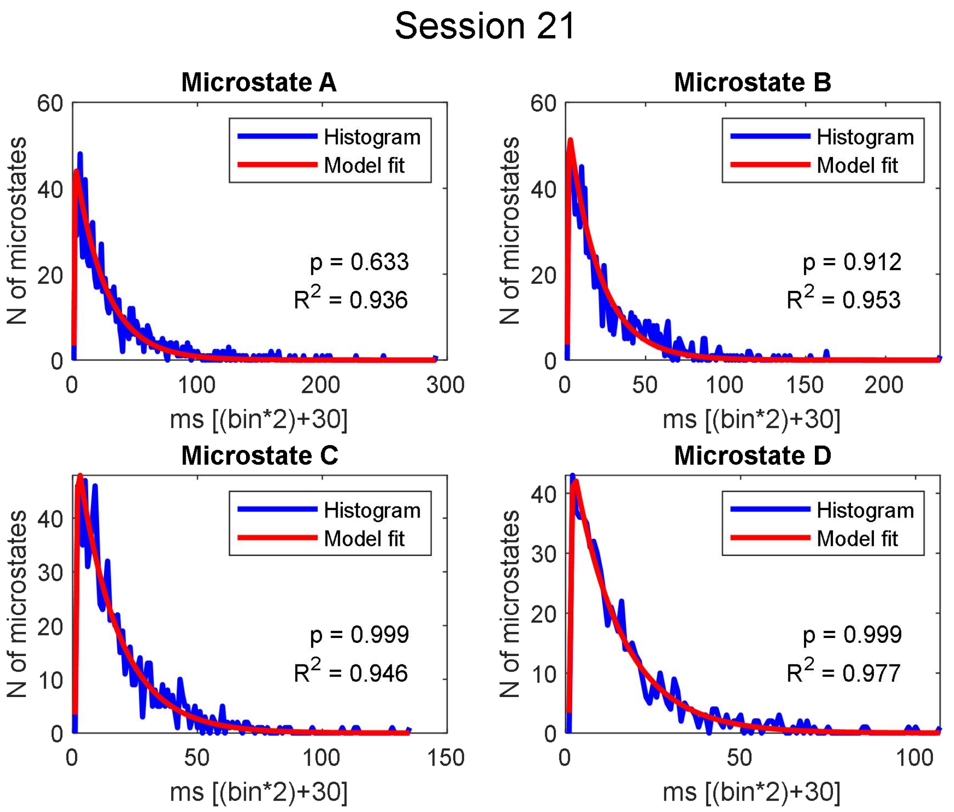
Session 21
Microstate A
Histogram
Model fit
p = 0.633
$R^2$ = 0.936
N of microstates
ms [(bin*2)+30]
Microstate B
Histogram
Model fit
p = 0.912
$R^2$ = 0.953
N of microstates
ms [(bin*2)+30]
Microstate C
Histogram
Model fit
p = 0.999
$R^2$ = 0.946
N of microstates
ms [(bin*2)+30]
Microstate D
Histogram
Model fit
p = 0.999
$R^2$ = 0.977
N of microstates
ms [(bin*2)+30]

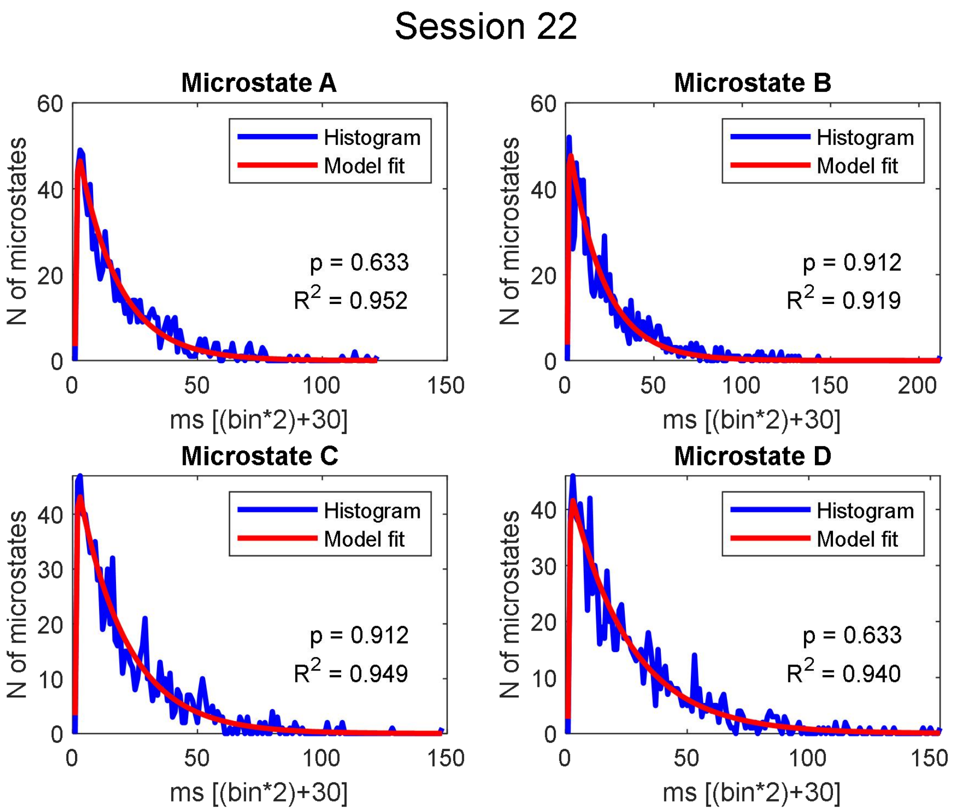
Session 22
Microstate A
Histogram
Model fit
p = 0.633
$R^2$ = 0.952
N of microstates
ms [(bin*2)+30]
Microstate B
Histogram
Model fit
p = 0.912
$R^2$ = 0.919
N of microstates
ms [(bin*2)+30]
Microstate C
Histogram
Model fit
p = 0.912
$R^2$ = 0.949
N of microstates
ms [(bin*2)+30]
Microstate D
Histogram
Model fit
p = 0.633
$R^2$ = 0.940
N of microstates
ms [(bin*2)+30]

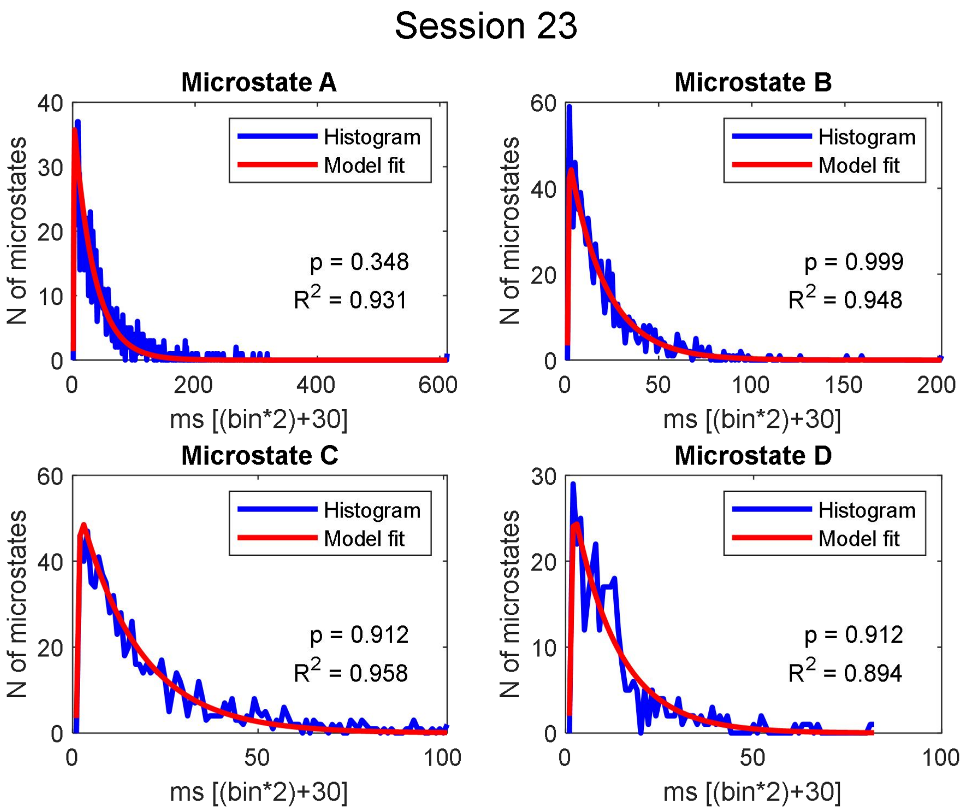
Session 23
Microstate A
Histogram
Model fit
p = 0.348
R2 = 0.931
N of microstates
ms [(bin*2)+30]
Microstate B
Histogram
Model fit
p = 0.999
R2 = 0.948
N of microstates
ms [(bin*2)+30]
Microstate C
Histogram
Model fit
p = 0.912
R2 = 0.958
N of microstates
ms [(bin*2)+30]
Microstate D
Histogram
Model fit
p = 0.912
R2 = 0.894
N of microstates
ms [(bin*2)+30]

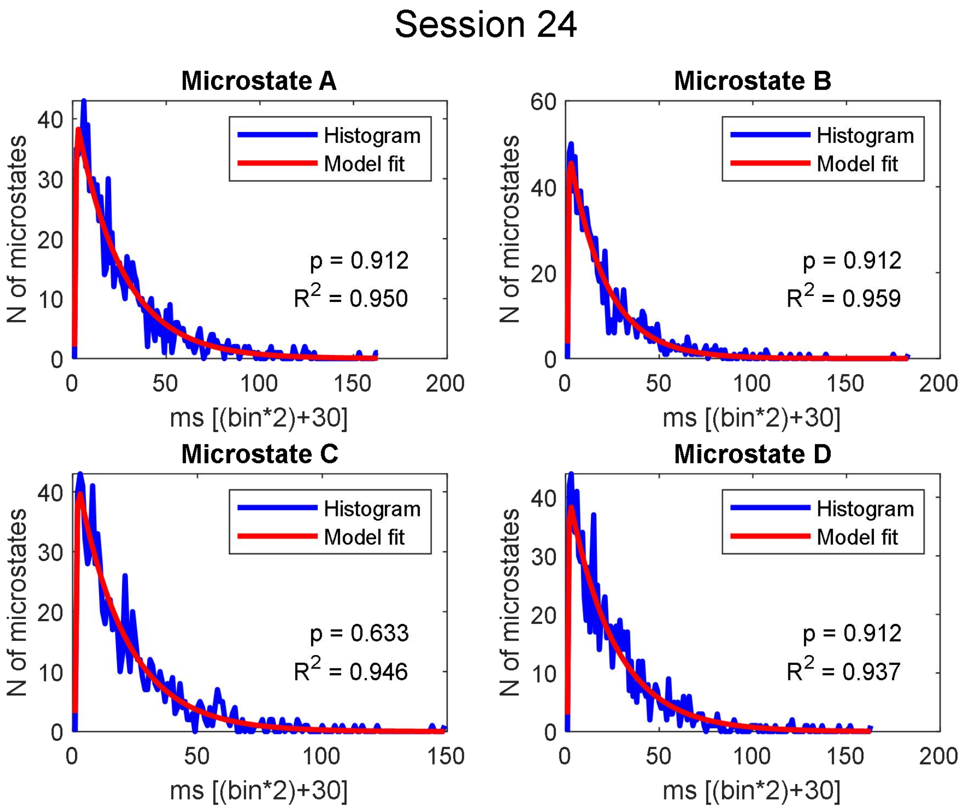
Session 24
Microstate A
Histogram
Model fit
p = 0.912
R2 = 0.950
N of microstates
ms [(bin*2)+30]
Microstate B
Histogram
Model fit
p = 0.912
R2 = 0.959
N of microstates
ms [(bin*2)+30]
Microstate C
Histogram
Model fit
p = 0.633
R2 = 0.946
N of microstates
ms [(bin*2)+30]
Microstate D
Histogram
Model fit
p = 0.912
R2 = 0.937
N of microstates
ms [(bin*2)+30]

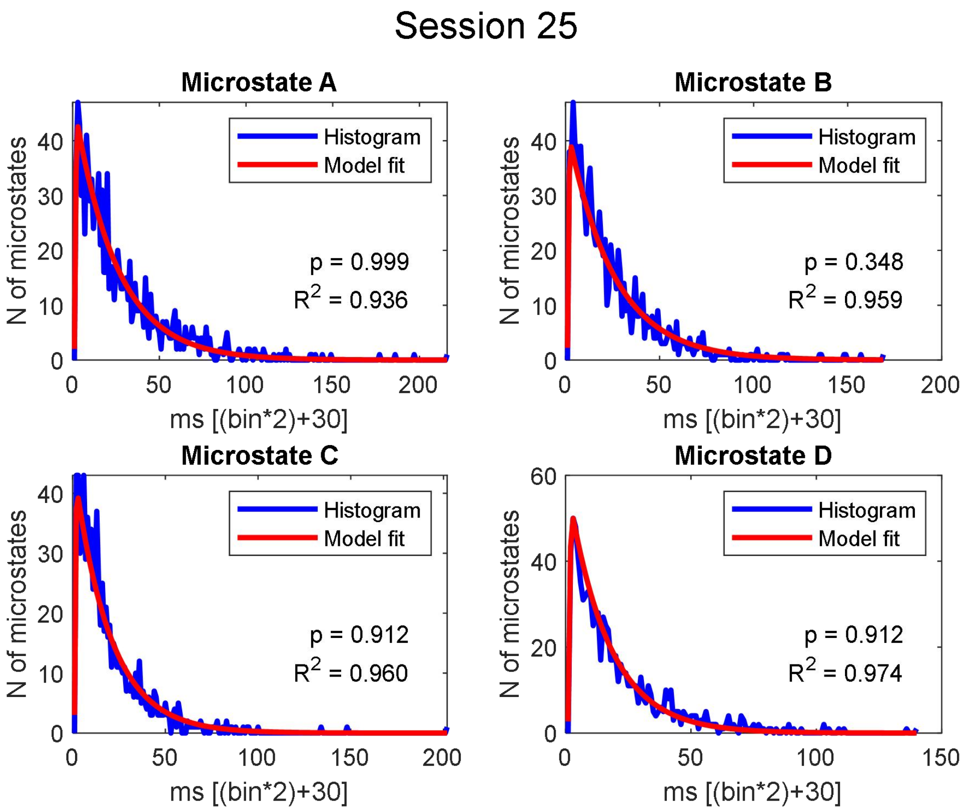

Session 25
Microstate A
Histogram
Model fit
p = 0.999
R2 = 0.936
N of microstates
ms [(bin*2)+30]
Microstate B
Histogram
Model fit
p = 0.348
R2 = 0.959
N of microstates
ms [(bin*2)+30]
Microstate C
Histogram
Model fit
p = 0.912
R2 = 0.960
N of microstates
ms [(bin*2)+30]
Microstate D
Histogram
Model fit
p = 0.912
R2 = 0.974
N of microstates
ms [(bin*2)+30]


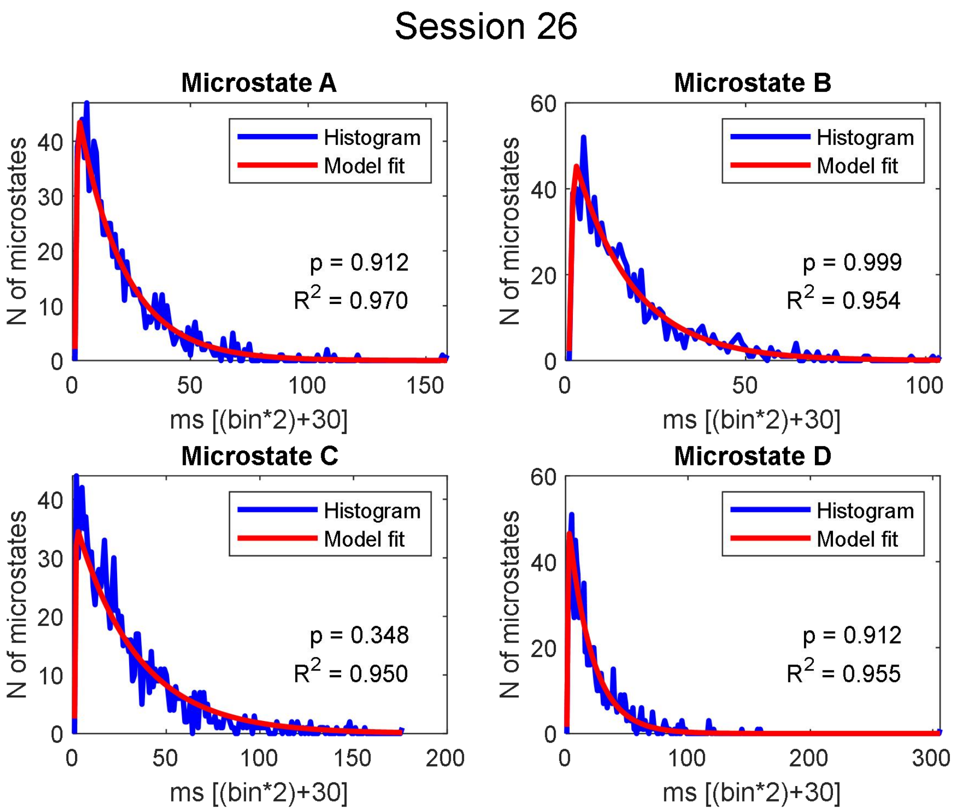

Session 26
Microstate A
Histogram
Model fit
p = 0.912
R2 = 0.970
N of microstates
ms [(bin*2)+30]
Microstate B
Histogram
Model fit
p = 0.999
R2 = 0.954
N of microstates
ms [(bin*2)+30]
Microstate C
Histogram
Model fit
p = 0.348
R2 = 0.950
N of microstates
ms [(bin*2)+30]
Microstate D
Histogram
Model fit
p = 0.912
R2 = 0.955
N of microstates
ms [(bin*2)+30]

#### Session 27

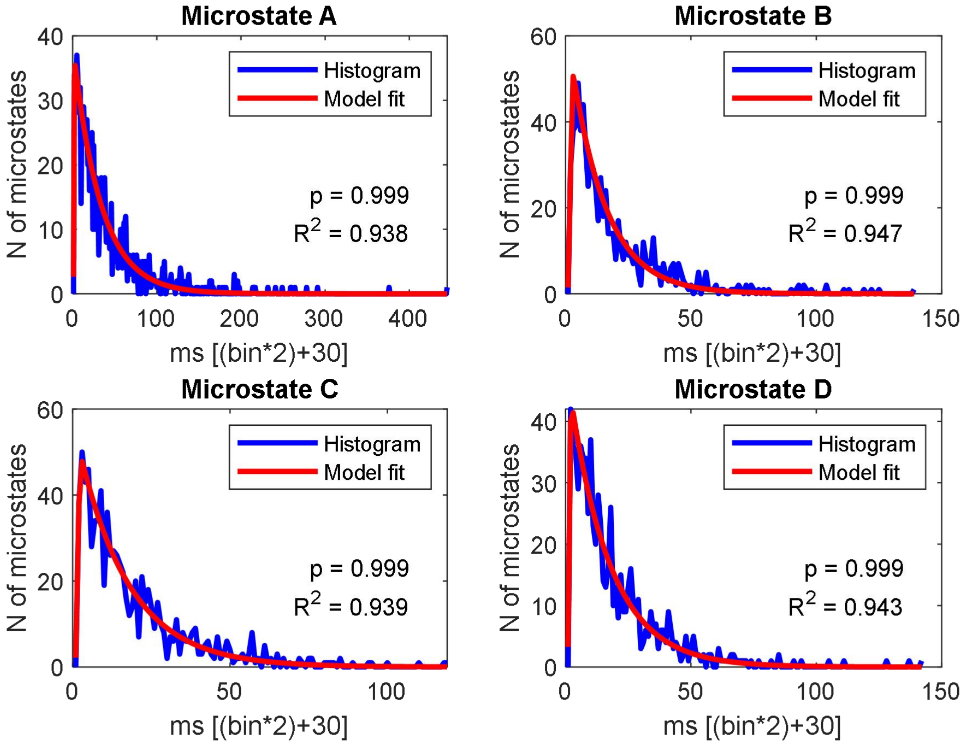


#### Session 28

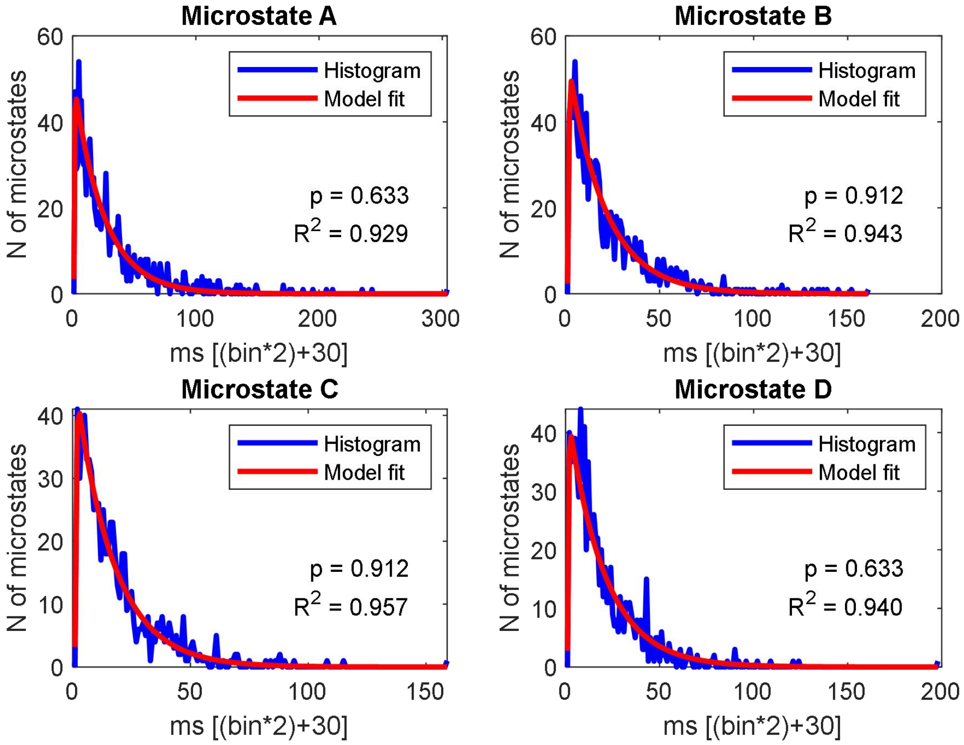

#### Session 29

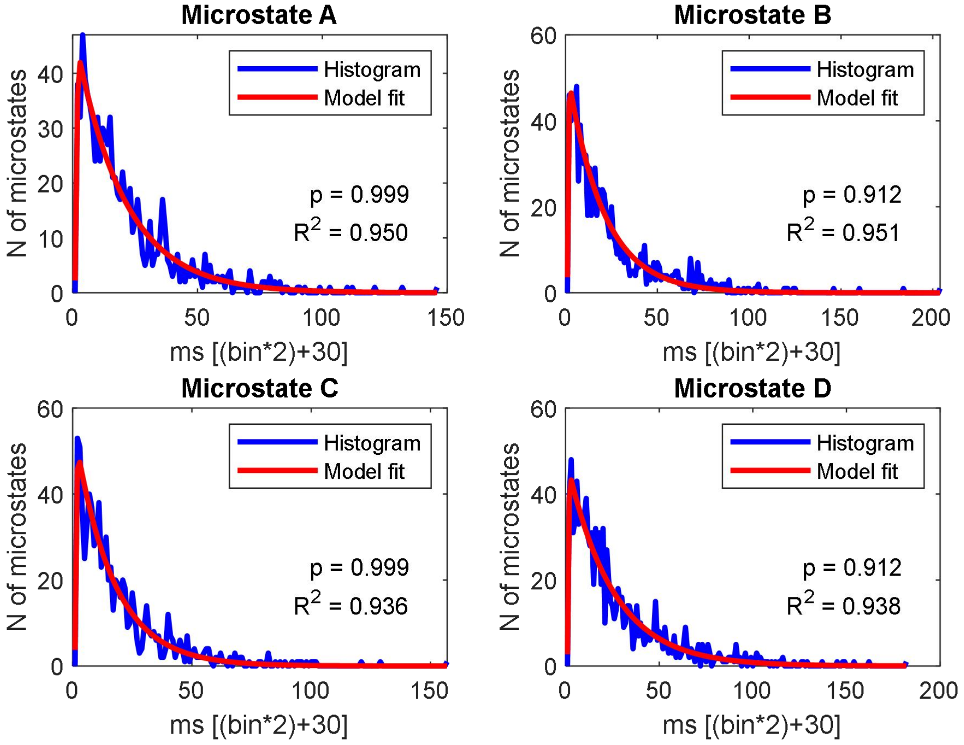


#### Session 30

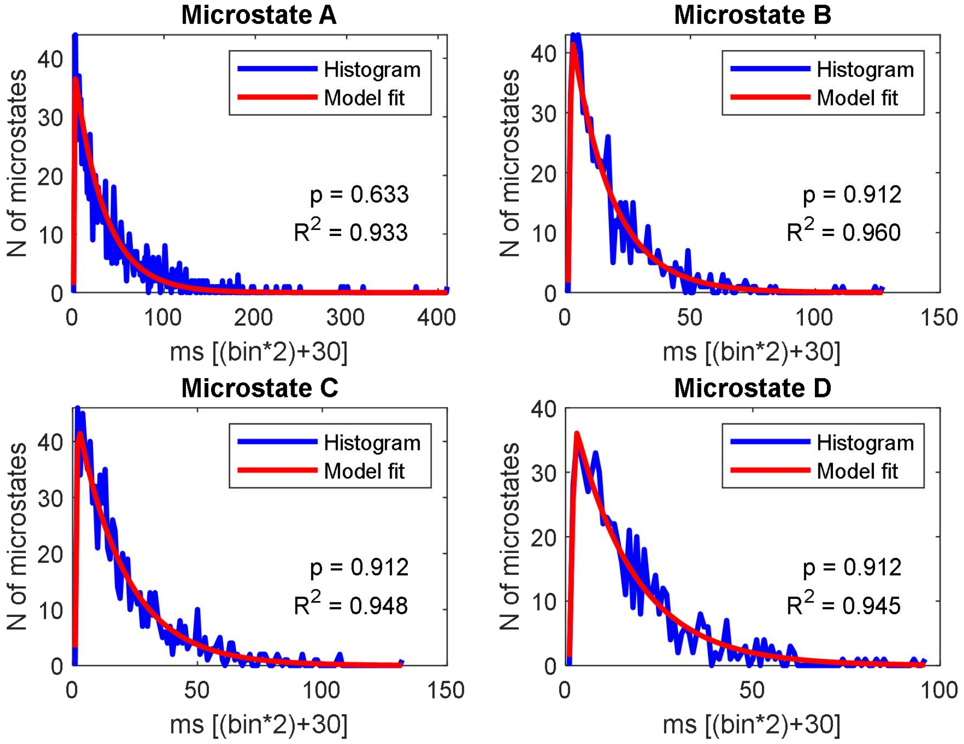

Session 31

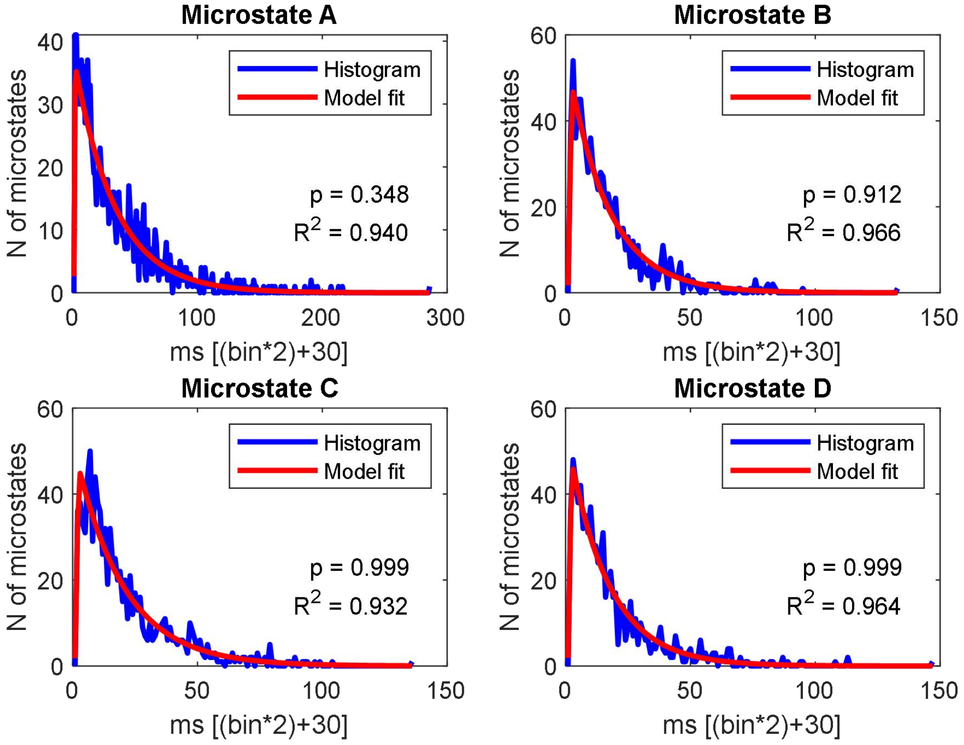


Session 32

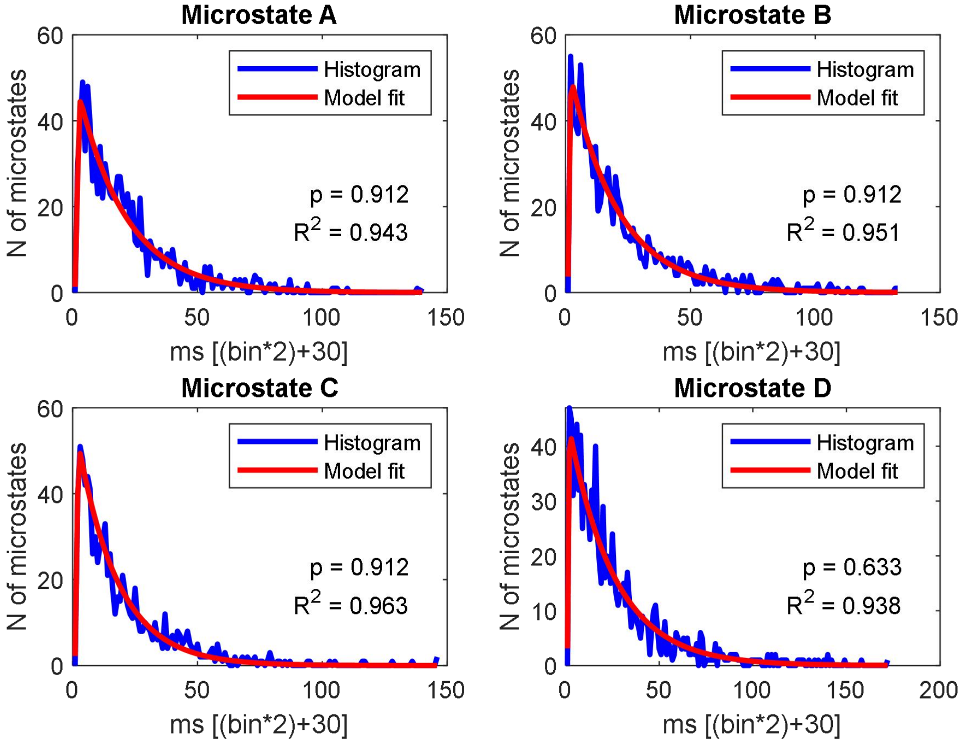

#### Session 33

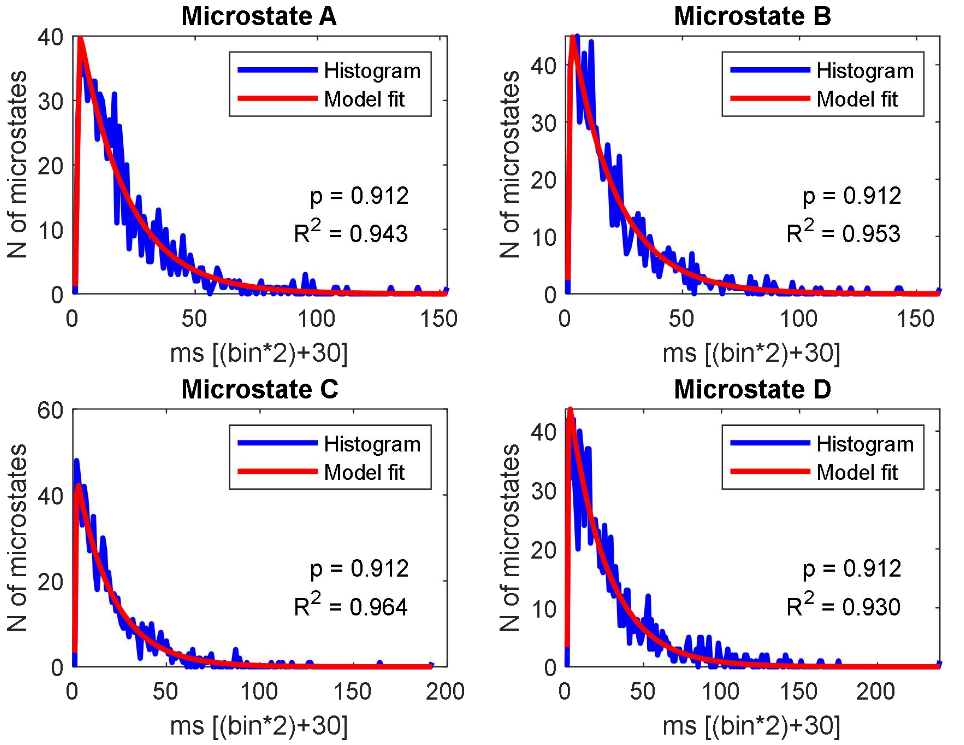


#### Session 34

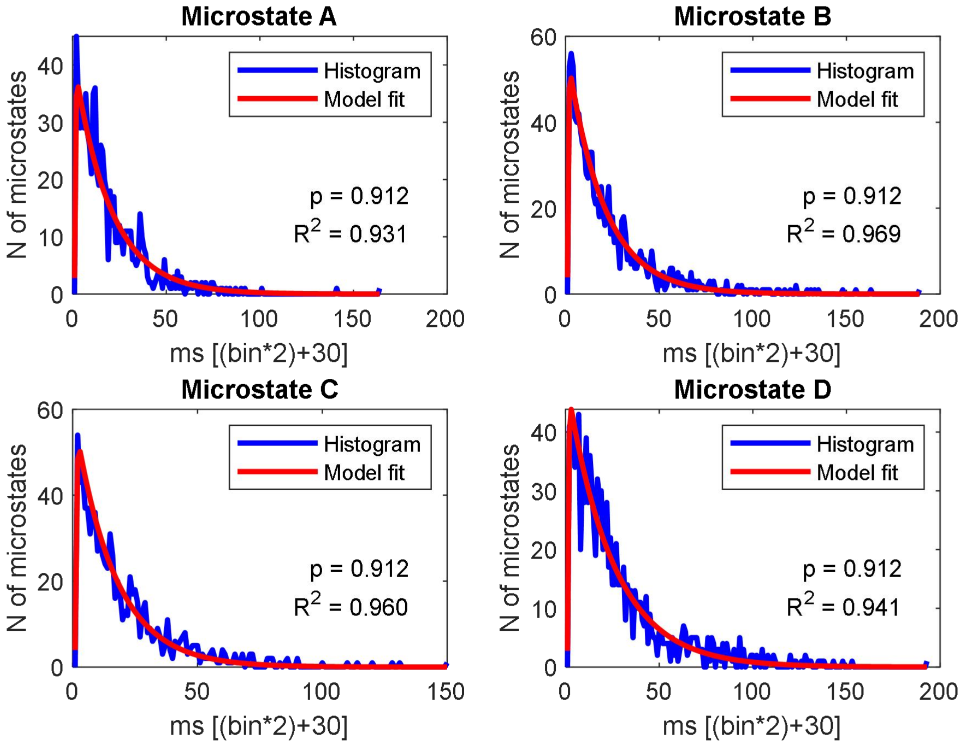

#### Session 35

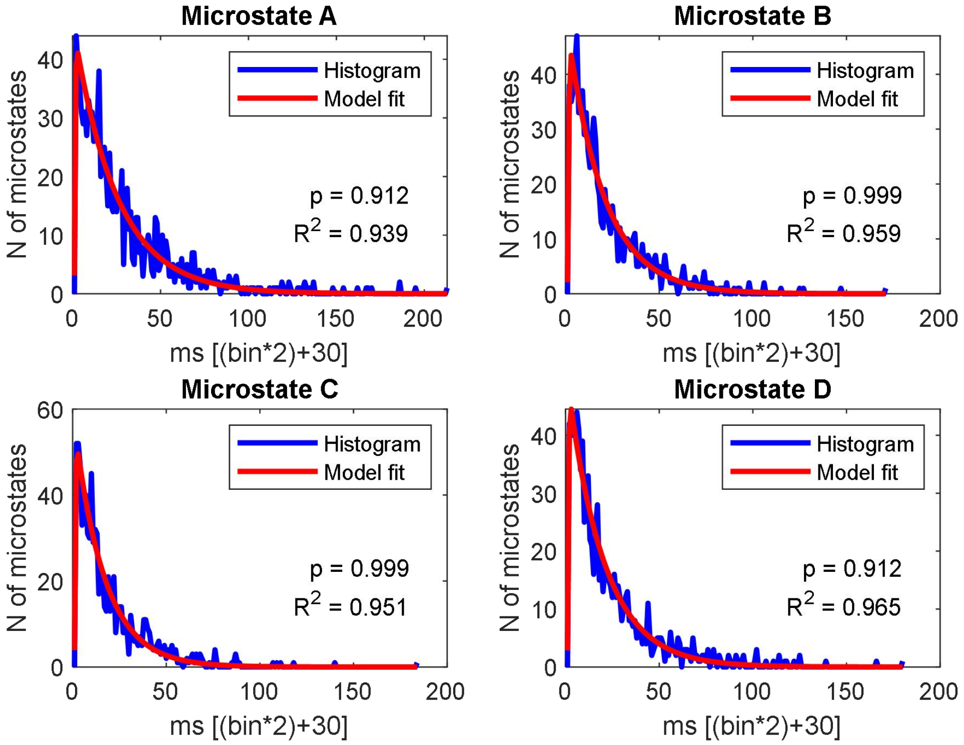


#### Session 36

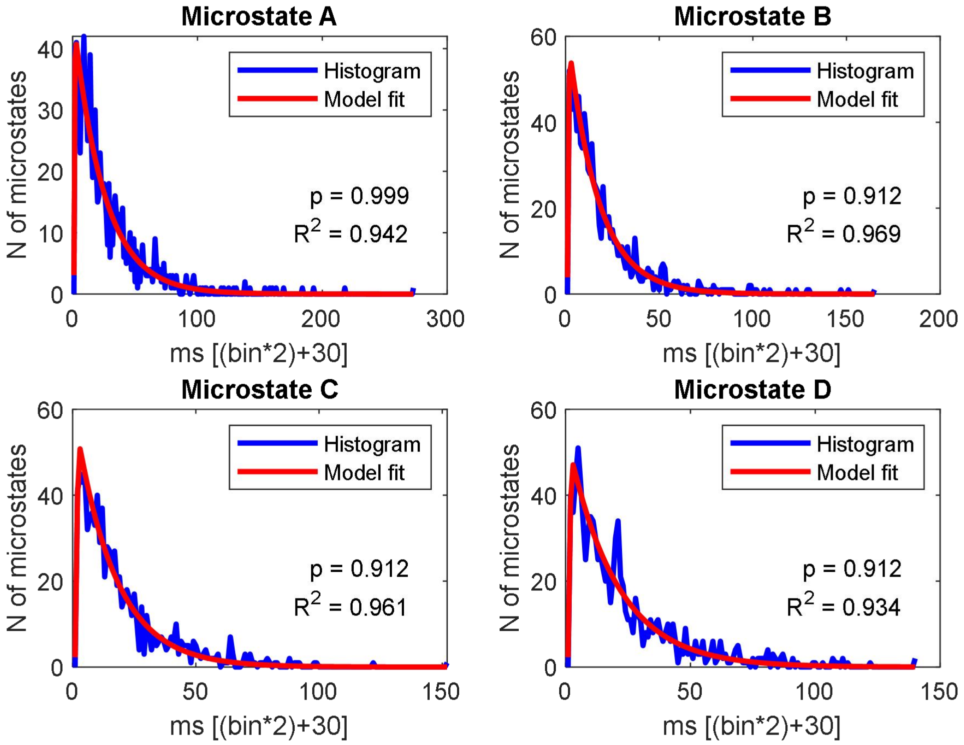

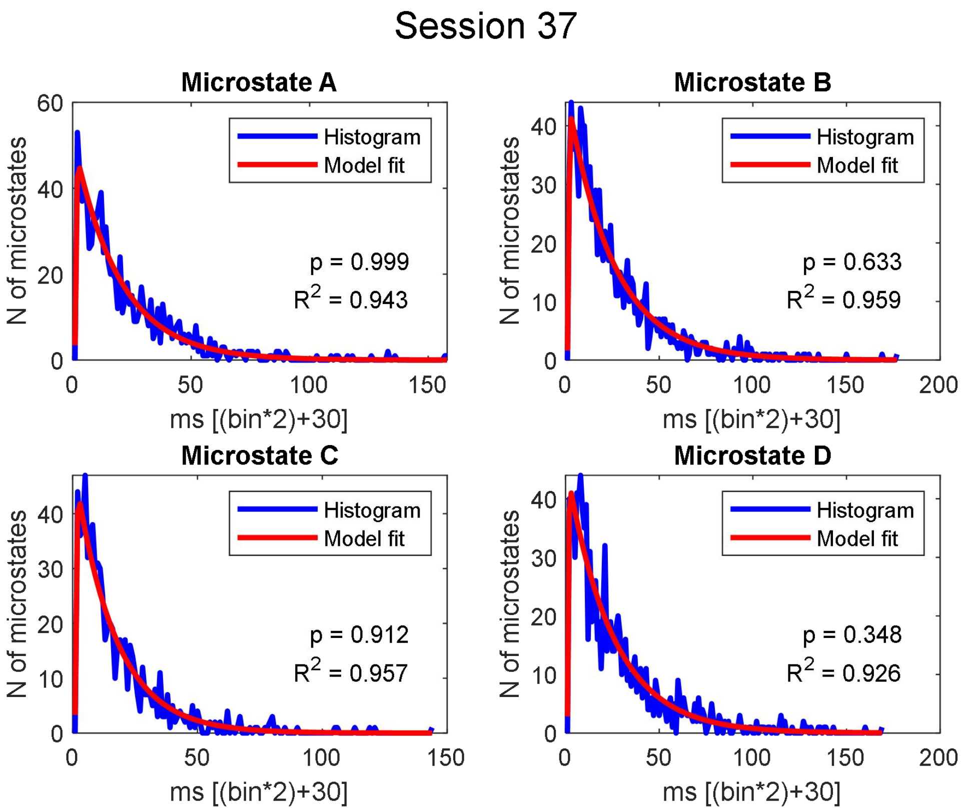
Session 37
Microstate A
Histogram
Model fit
p = 0.999
R2 = 0.943
N of microstates
ms [(bin*2)+30]
Microstate B
Histogram
Model fit
p = 0.633
R2 = 0.959
N of microstates
ms [(bin*2)+30]
Microstate C
Histogram
Model fit
p = 0.912
R2 = 0.957
N of microstates
ms [(bin*2)+30]
Microstate D
Histogram
Model fit
p = 0.348
R2 = 0.926
N of microstates
ms [(bin*2)+30]

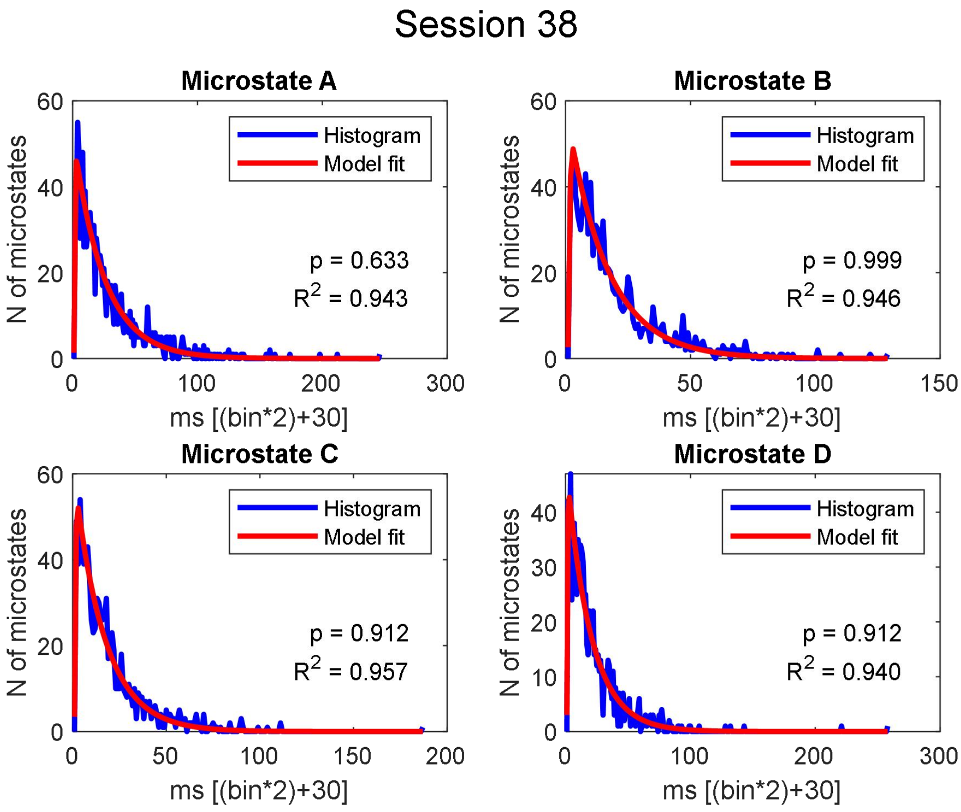
Session 38
Microstate A
Histogram
Model fit
p = 0.633
R2 = 0.943
N of microstates
ms [(bin*2)+30]
Microstate B
Histogram
Model fit
p = 0.999
R2 = 0.946
N of microstates
ms [(bin*2)+30]
Microstate C
Histogram
Model fit
p = 0.912
R2 = 0.957
N of microstates
ms [(bin*2)+30]
Microstate D
Histogram
Model fit
p = 0.912
R2 = 0.940
N of microstates
ms [(bin*2)+30]

#### Session 39

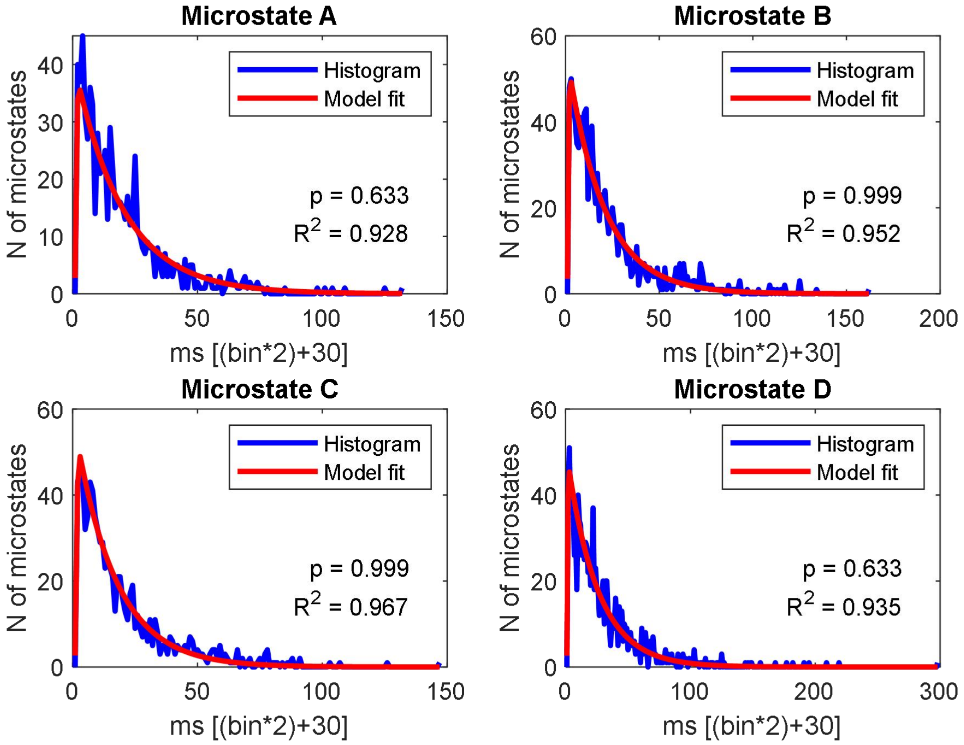


#### Session 40

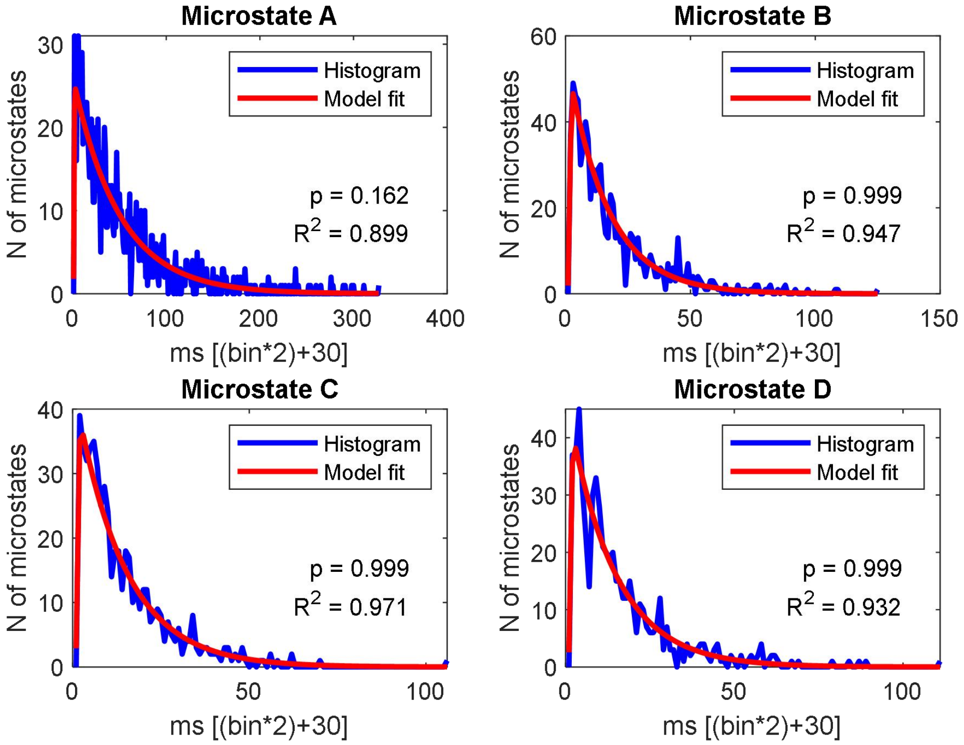

### Session 41

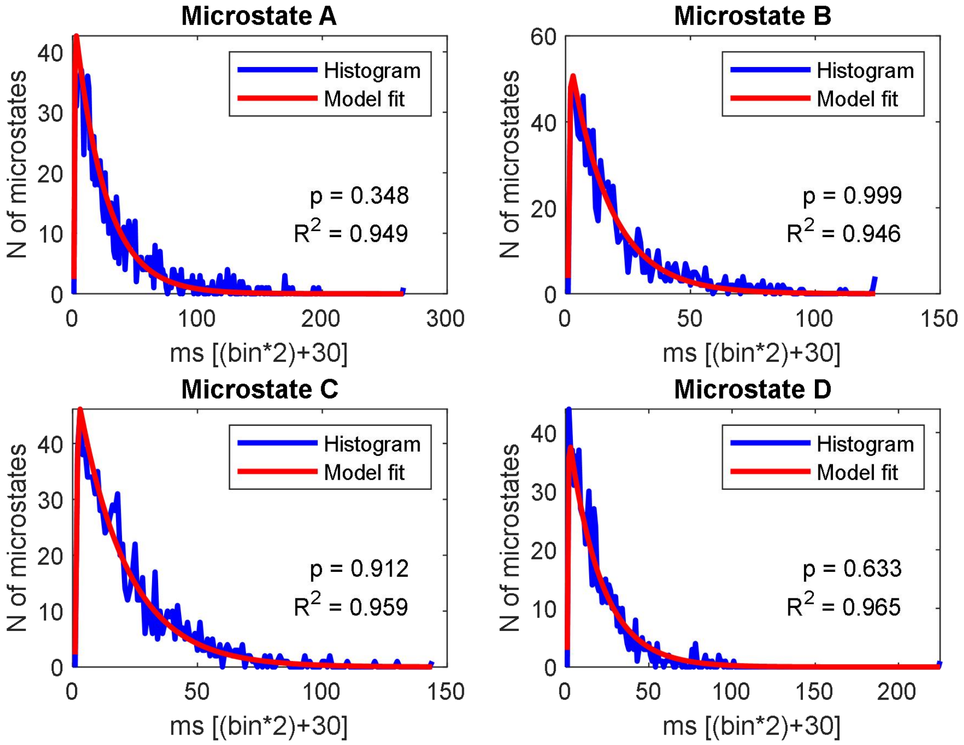


### Session 42

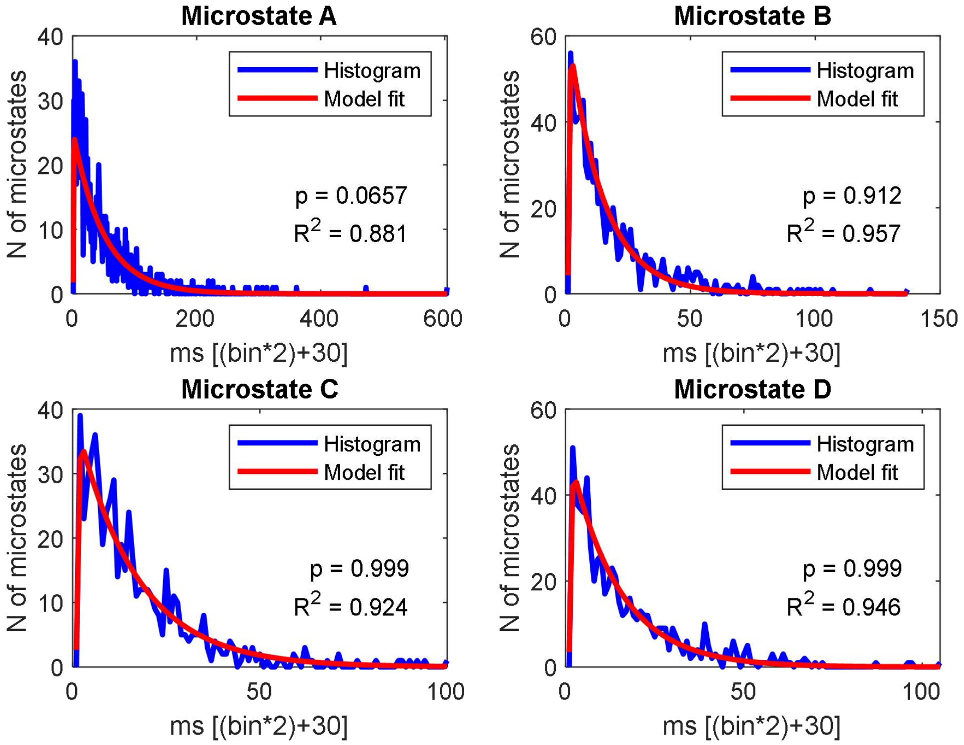

#### Session 43

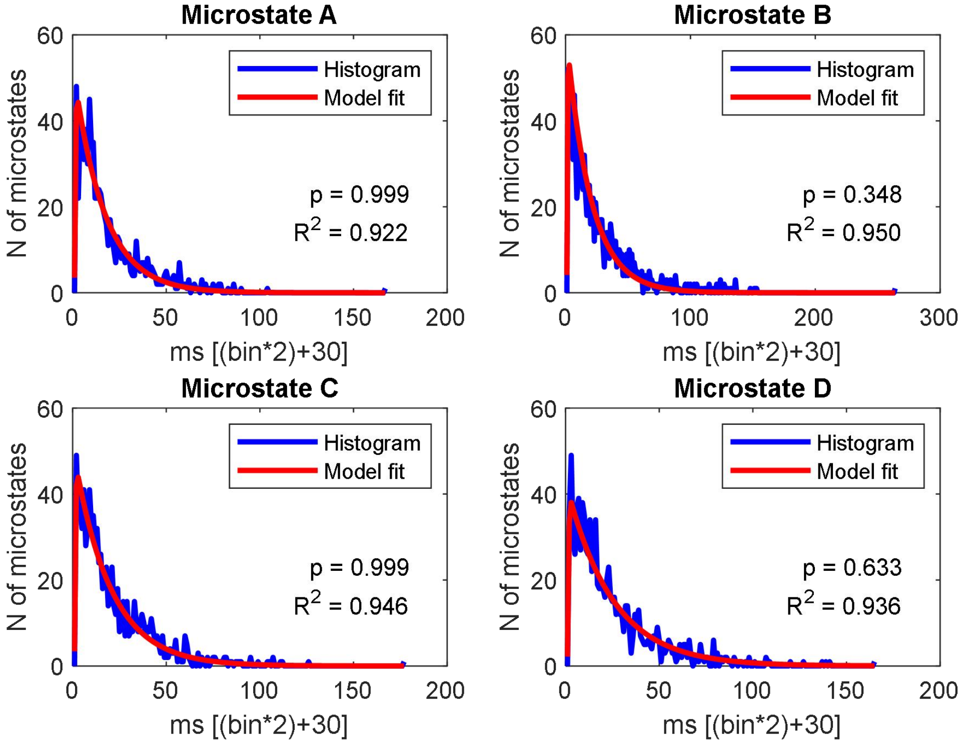


#### Session 44

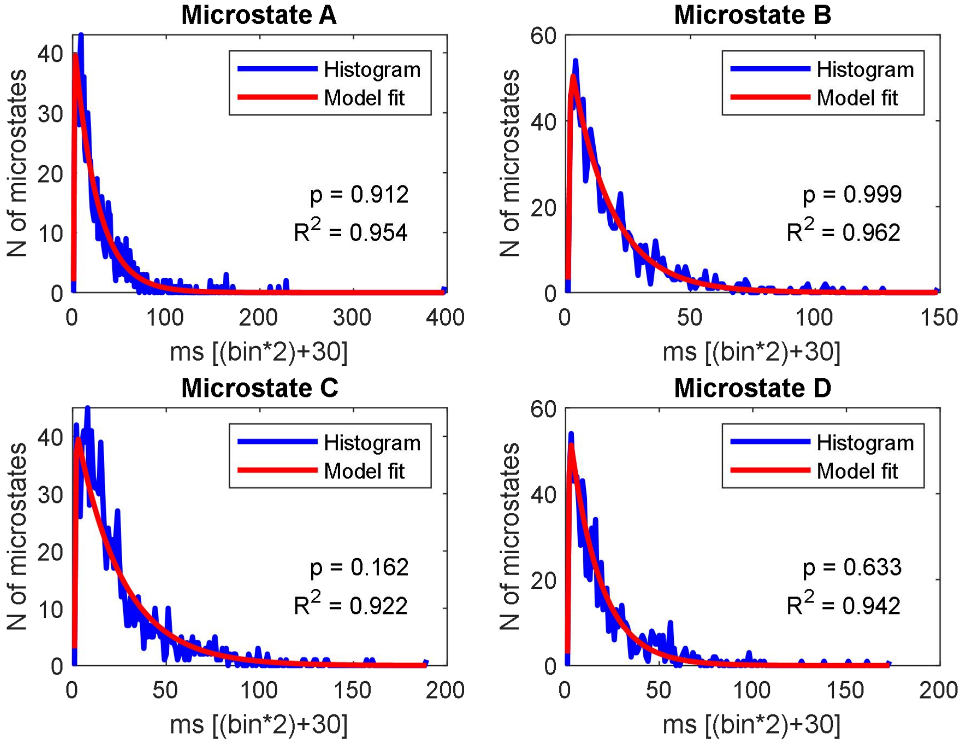

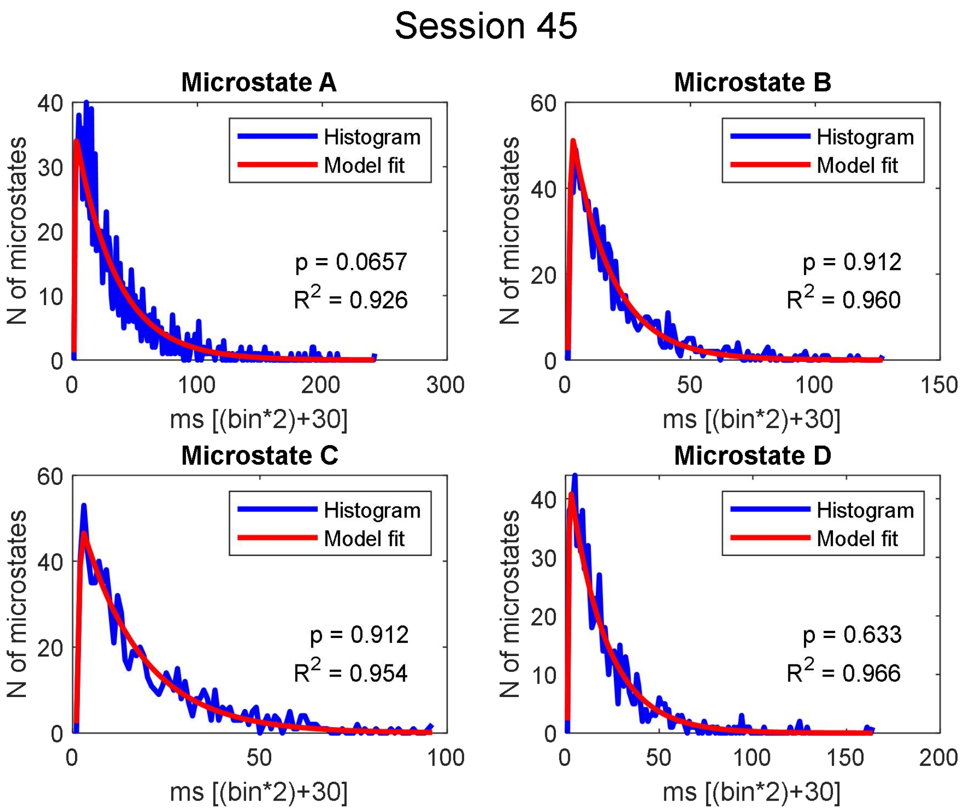
Session 45
Microstate A
Histogram
Model fit
p = 0.0657
R2 = 0.926
N of microstates
ms [(bin*2)+30]
Microstate B
Histogram
Model fit
p = 0.912
R2 = 0.960
N of microstates
ms [(bin*2)+30]
Microstate C
Histogram
Model fit
p = 0.912
R2 = 0.954
N of microstates
ms [(bin*2)+30]
Microstate D
Histogram
Model fit
p = 0.633
R2 = 0.966
N of microstates
ms [(bin*2)+30]

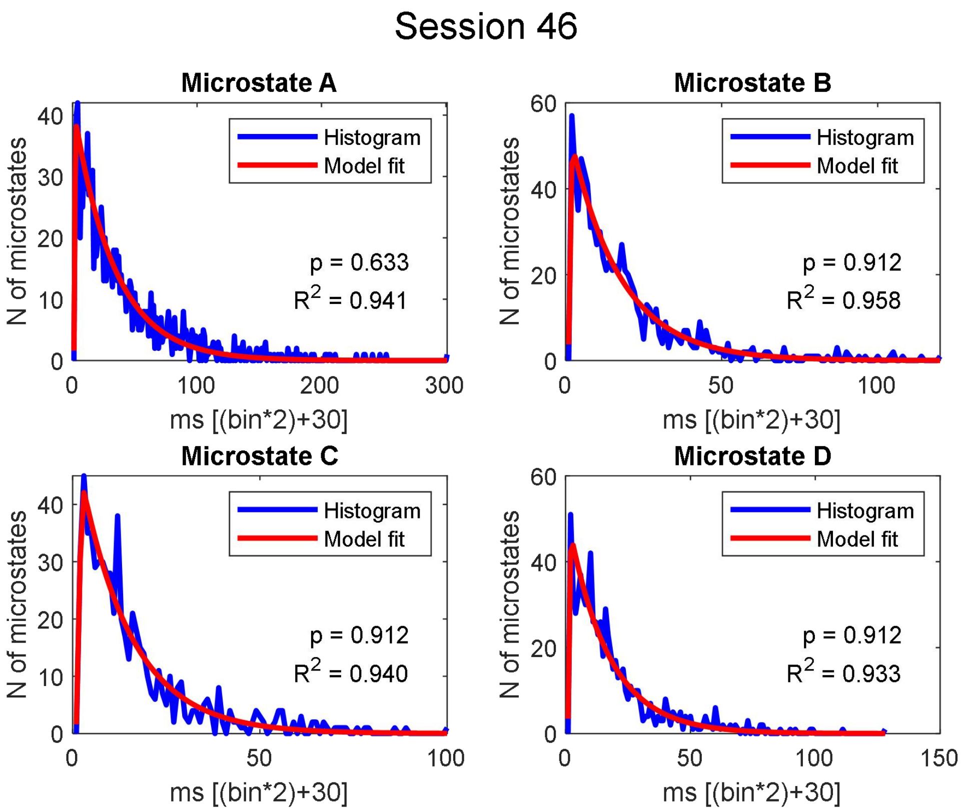
Session 46
Microstate A
Histogram
Model fit
p = 0.633
R2 = 0.941
N of microstates
ms [(bin*2)+30]
Microstate B
Histogram
Model fit
p = 0.912
R2 = 0.958
N of microstates
ms [(bin*2)+30]
Microstate C
Histogram
Model fit
p = 0.912
R2 = 0.940
N of microstates
ms [(bin*2)+30]
Microstate D
Histogram
Model fit
p = 0.912
R2 = 0.933
N of microstates
ms [(bin*2)+30]

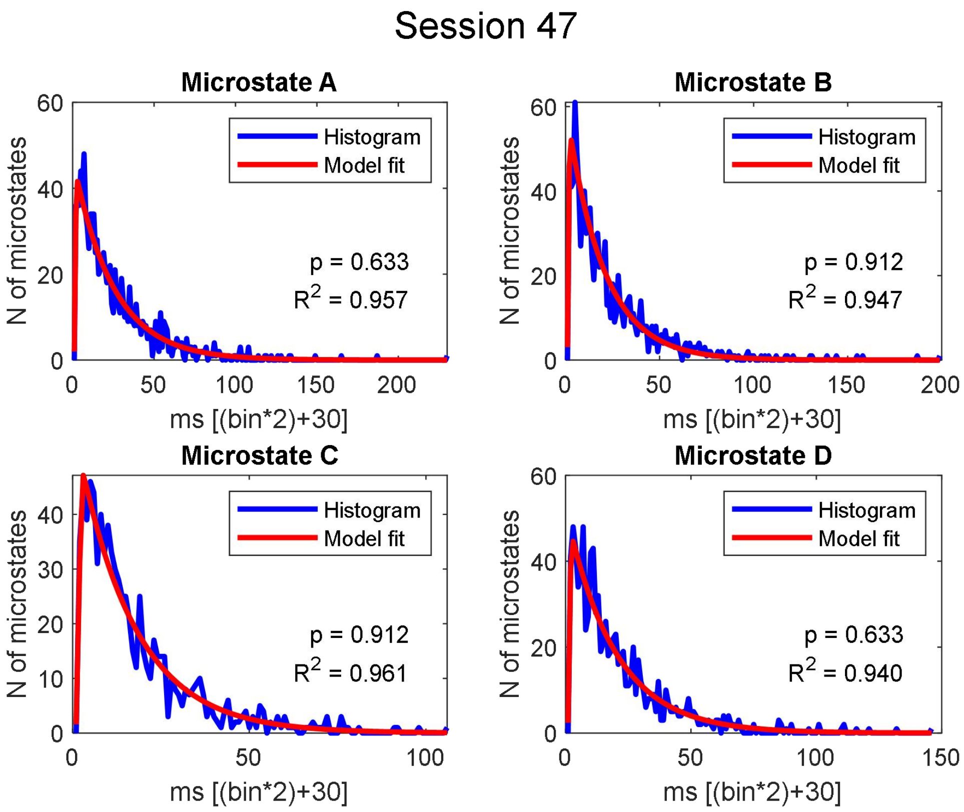
Session 47
Microstate A
Histogram
Model fit
p = 0.633
$R^2$ = 0.957
N of microstates
ms [(bin*2)+30]
Microstate B
Histogram
Model fit
p = 0.912
$R^2$ = 0.947
N of microstates
ms [(bin*2)+30]
Microstate C
Histogram
Model fit
p = 0.912
$R^2$ = 0.961
N of microstates
ms [(bin*2)+30]
Microstate D
Histogram
Model fit
p = 0.633
$R^2$ = 0.940
N of microstates
ms [(bin*2)+30]

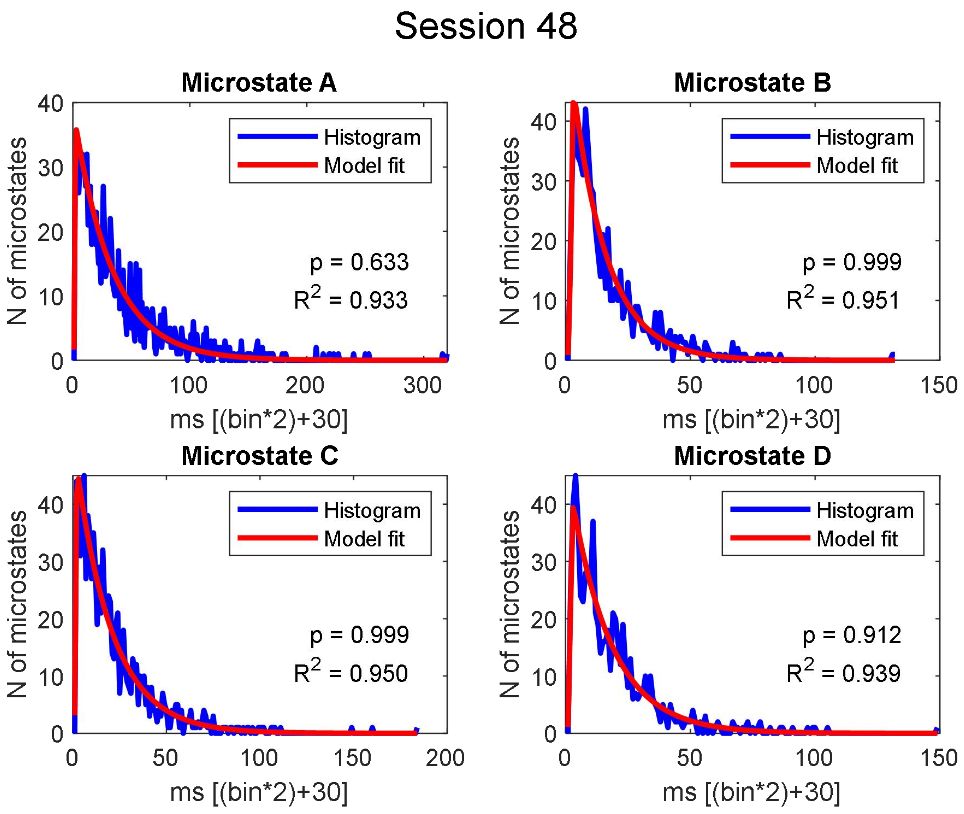
Session 48
Microstate A
Histogram
Model fit
p = 0.633
$R^2$ = 0.933
N of microstates
ms [(bin*2)+30]
Microstate B
Histogram
Model fit
p = 0.999
$R^2$ = 0.951
N of microstates
ms [(bin*2)+30]
Microstate C
Histogram
Model fit
p = 0.999
$R^2$ = 0.950
N of microstates
ms [(bin*2)+30]
Microstate D
Histogram
Model fit
p = 0.912
$R^2$ = 0.939
N of microstates
ms [(bin*2)+30]

#### Session 49

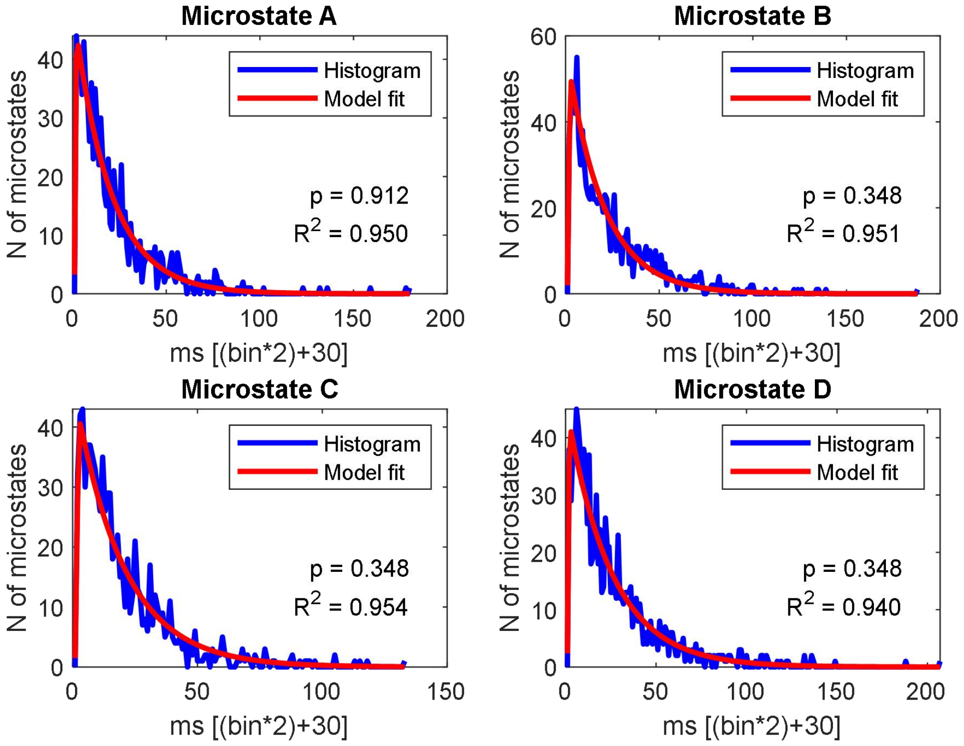


#### Session 50

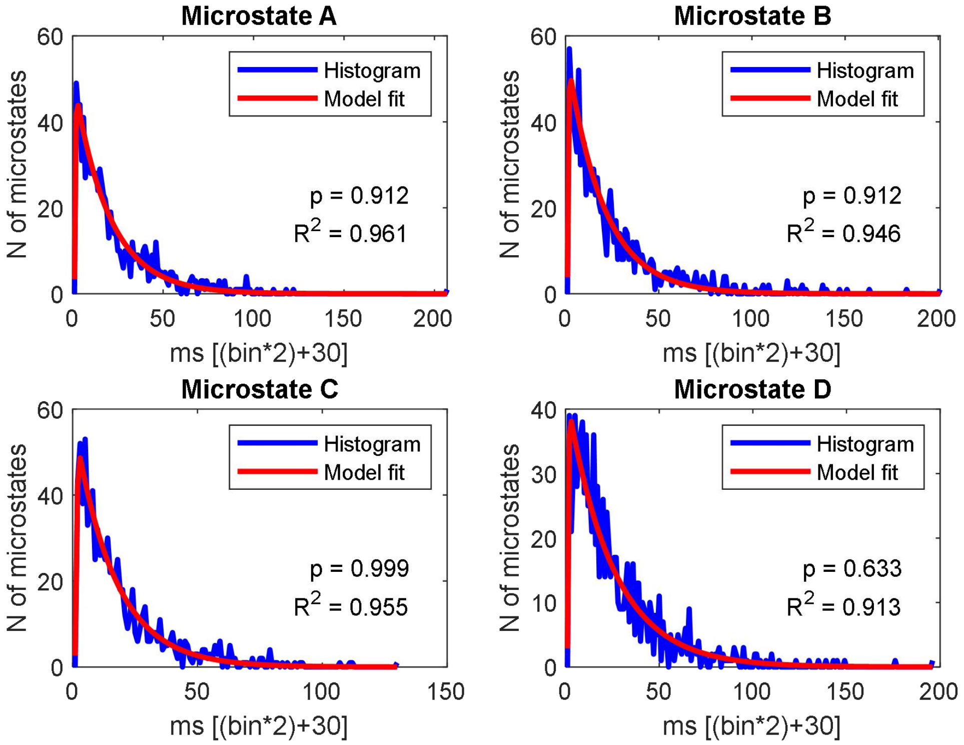

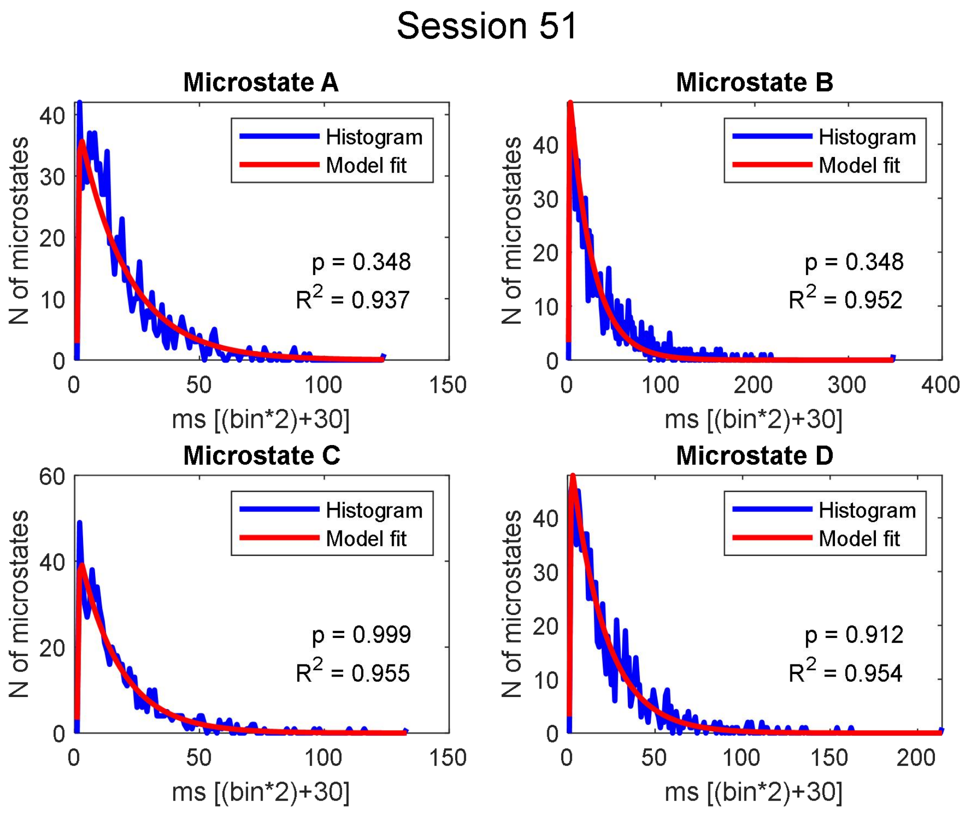

Session 51
Microstate A
Histogram
Model fit
p = 0.348
R2 = 0.937
N of microstates
ms [(bin*2)+30]
Microstate B
Histogram
Model fit
p = 0.348
R2 = 0.952
N of microstates
ms [(bin*2)+30]
Microstate C
Histogram
Model fit
p = 0.999
R2 = 0.955
N of microstates
ms [(bin*2)+30]
Microstate D
Histogram
Model fit
p = 0.912
R2 = 0.954
N of microstates
ms [(bin*2)+30]


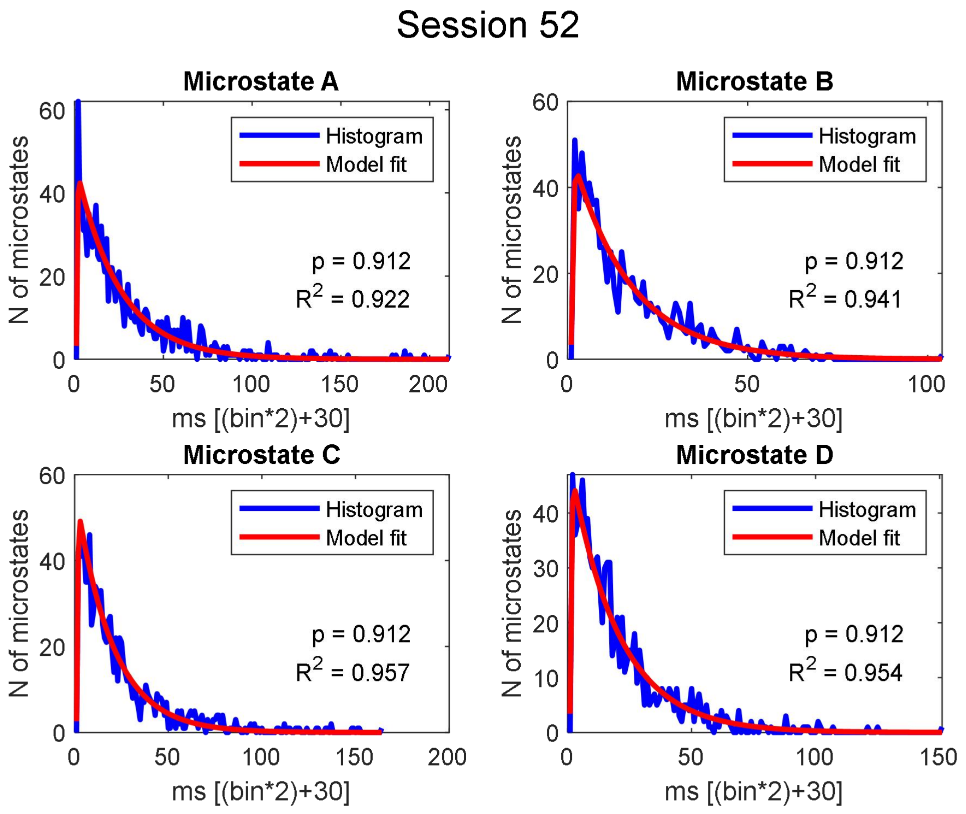

Session 52
Microstate A
Histogram
Model fit
p = 0.912
R2 = 0.922
N of microstates
ms [(bin*2)+30]
Microstate B
Histogram
Model fit
p = 0.912
R2 = 0.941
N of microstates
ms [(bin*2)+30]
Microstate C
Histogram
Model fit
p = 0.912
R2 = 0.957
N of microstates
ms [(bin*2)+30]
Microstate D
Histogram
Model fit
p = 0.912
R2 = 0.954
N of microstates
ms [(bin*2)+30]

#### Session 53

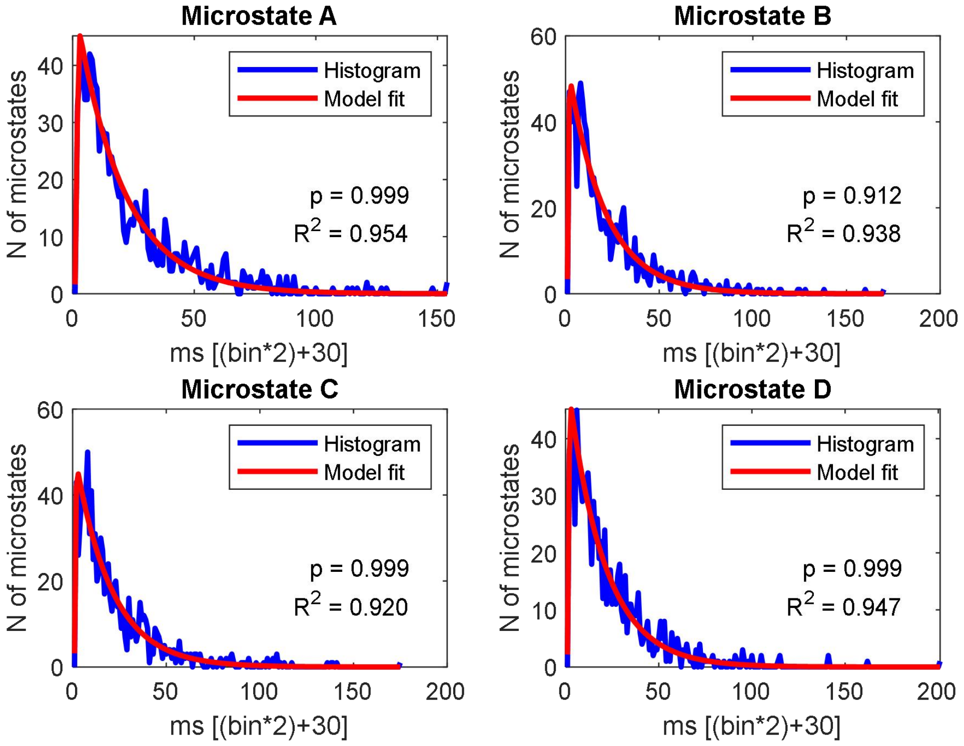


#### Session 54

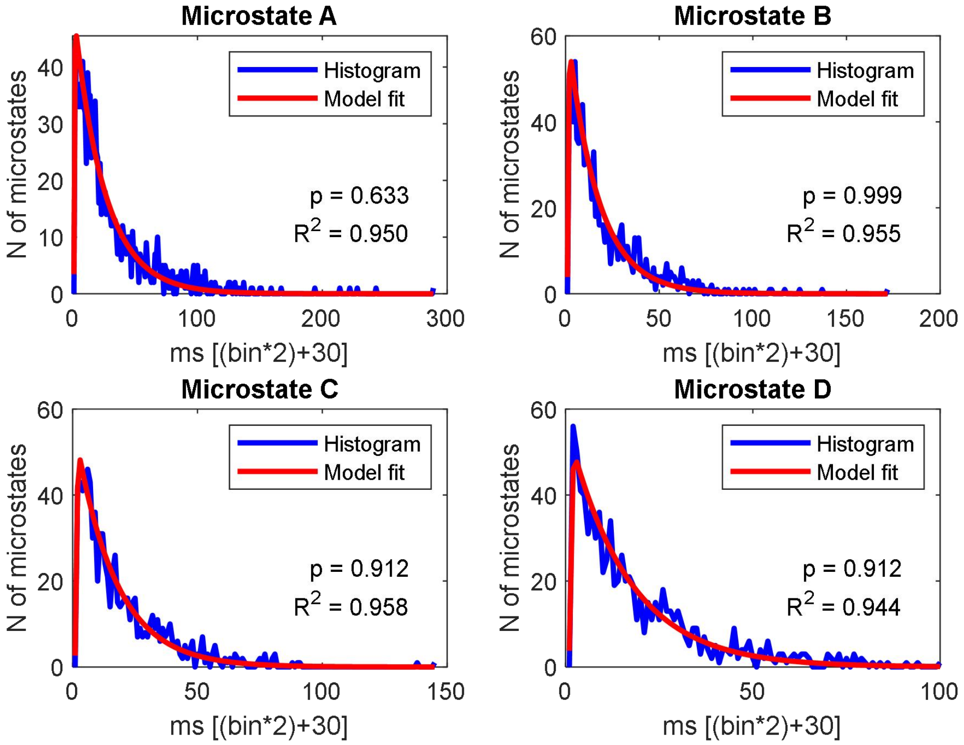

#### Session 55

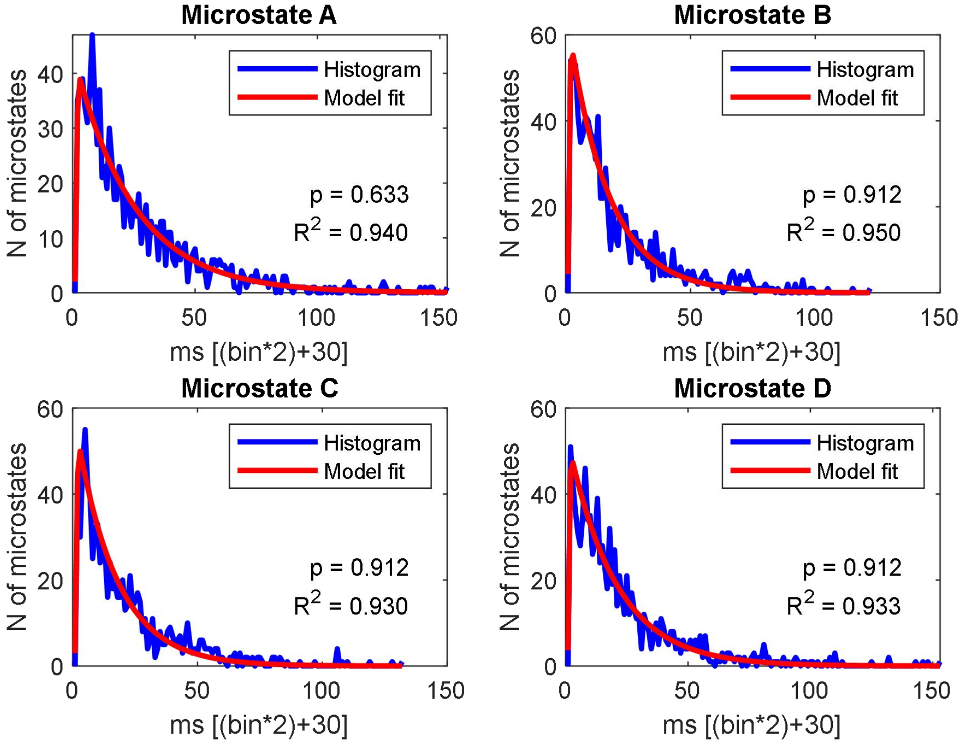


#### Session 56

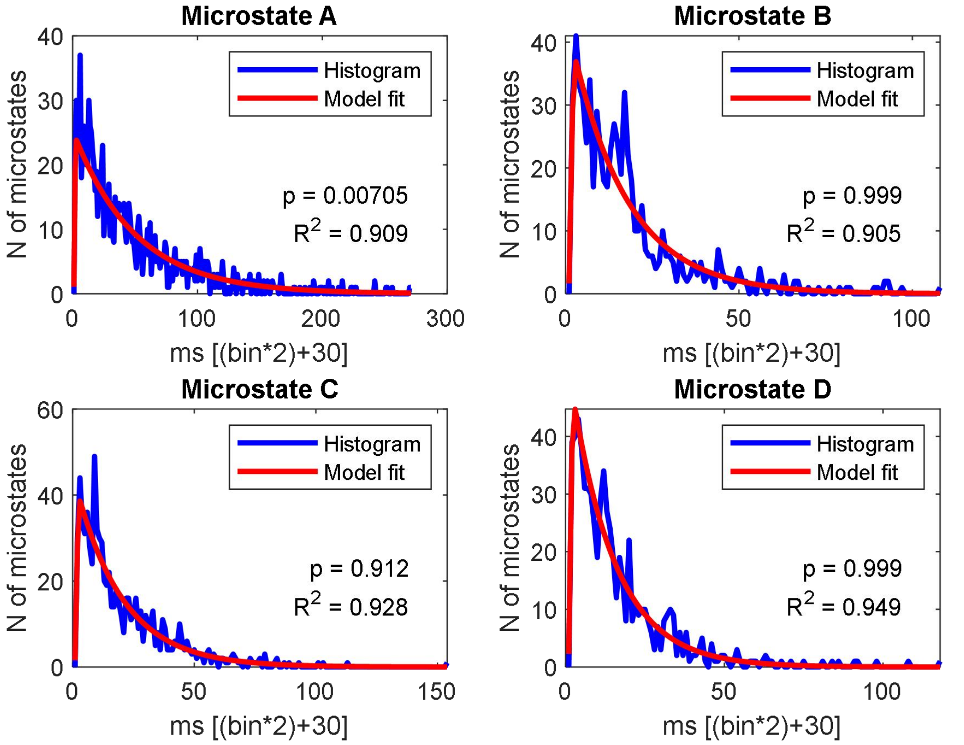

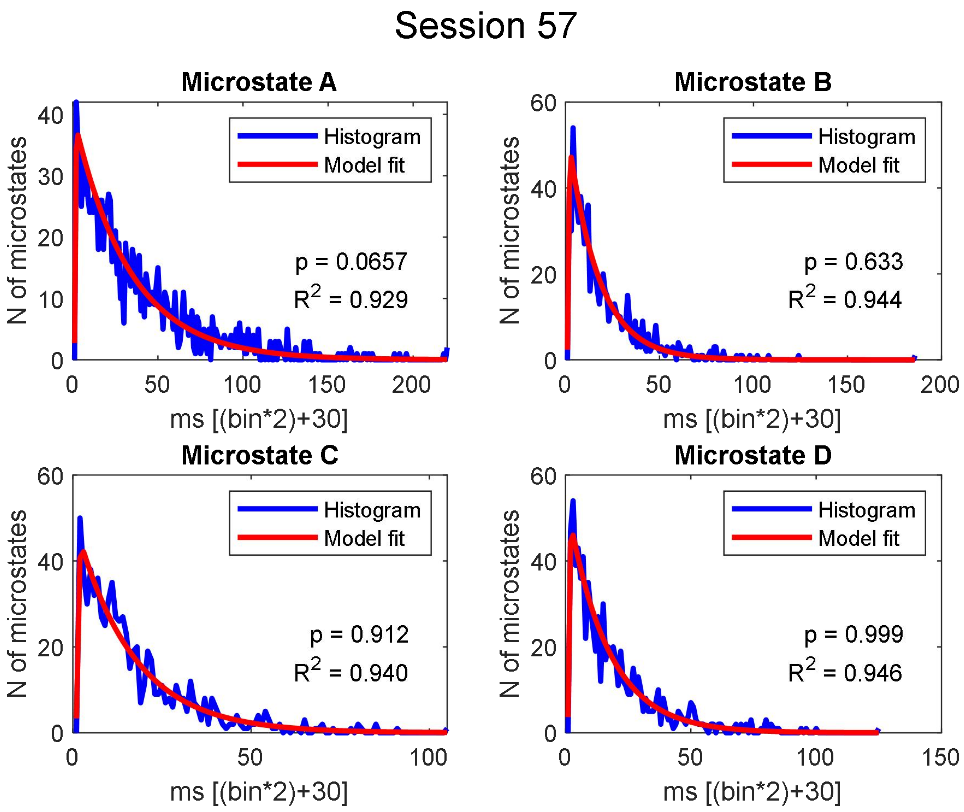
Session 57
Microstate A
Histogram
Model fit
p = 0.0657
R2 = 0.929
N of microstates
ms [(bin*2)+30]
Microstate B
Histogram
Model fit
p = 0.633
R2 = 0.944
N of microstates
ms [(bin*2)+30]
Microstate C
Histogram
Model fit
p = 0.912
R2 = 0.940
N of microstates
ms [(bin*2)+30]
Microstate D
Histogram
Model fit
p = 0.999
R2 = 0.946
N of microstates
ms [(bin*2)+30]

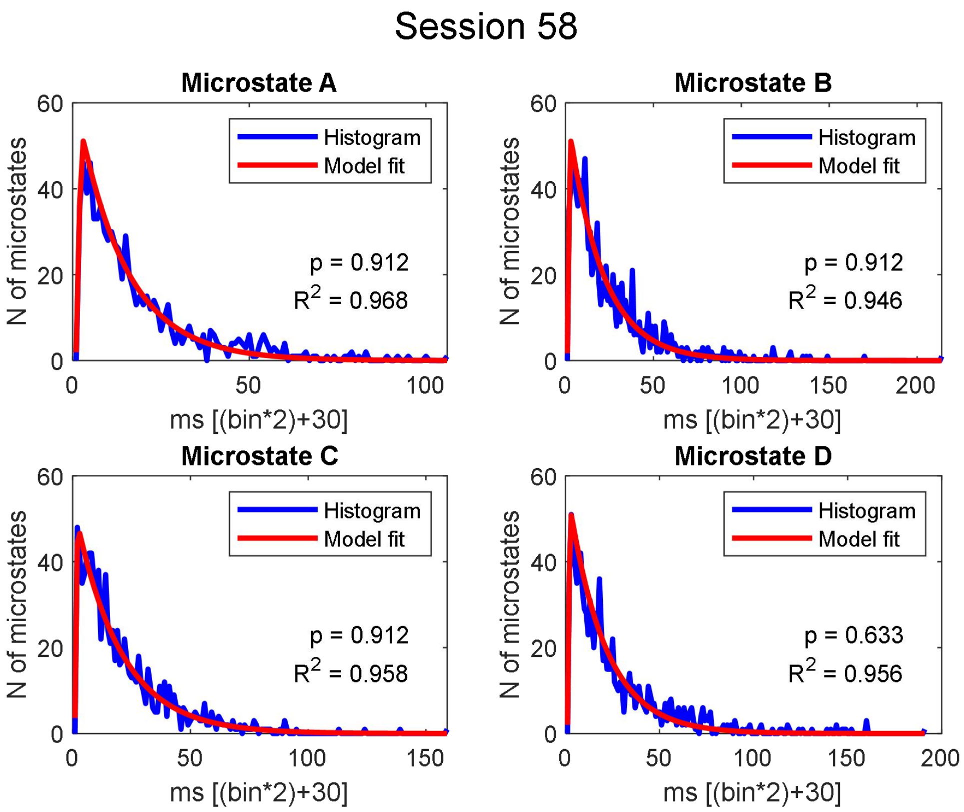
Session 58
Microstate A
Histogram
Model fit
p = 0.912
R2 = 0.968
N of microstates
ms [(bin*2)+30]
Microstate B
Histogram
Model fit
p = 0.912
R2 = 0.946
N of microstates
ms [(bin*2)+30]
Microstate C
Histogram
Model fit
p = 0.912
R2 = 0.958
N of microstates
ms [(bin*2)+30]
Microstate D
Histogram
Model fit
p = 0.633
R2 = 0.956
N of microstates
ms [(bin*2)+30]

### Session 59

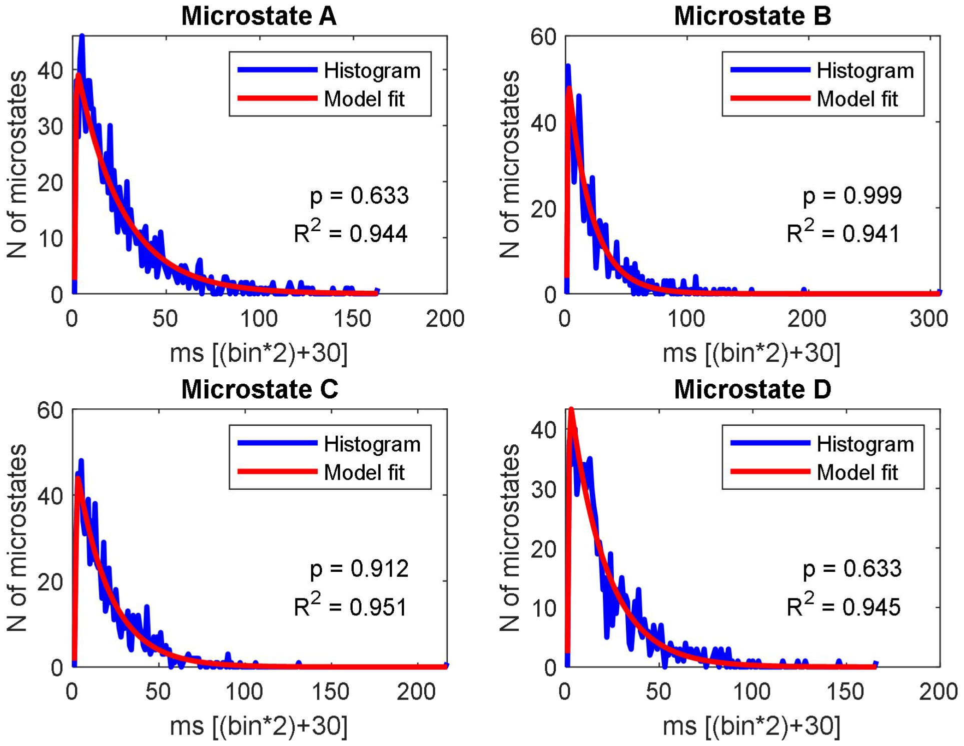


### Session 60

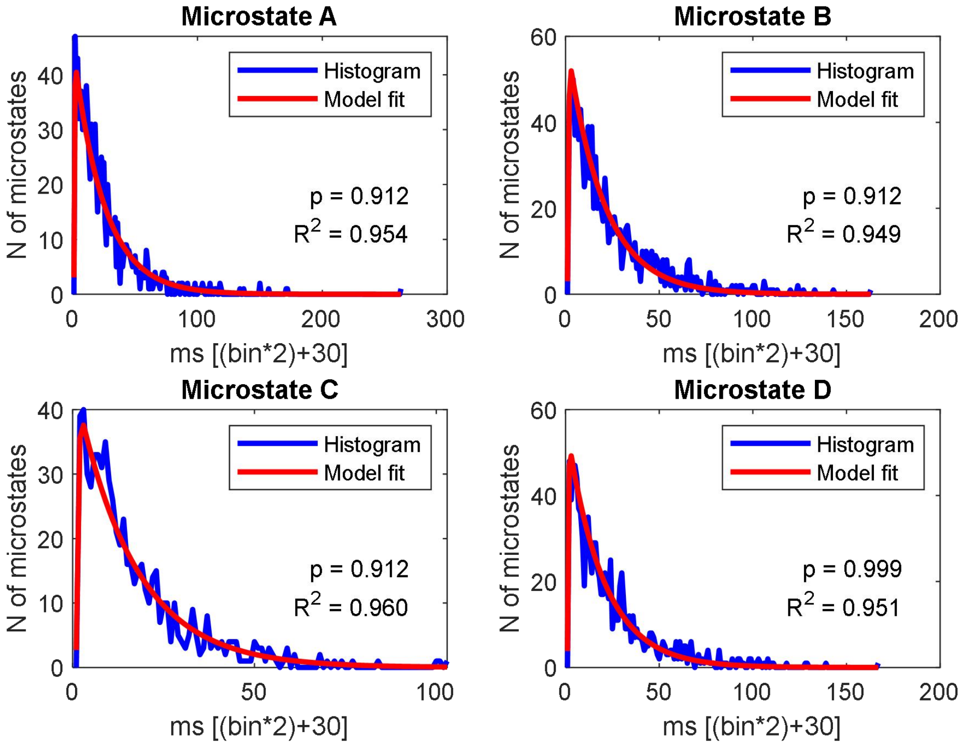

**Supplementary Figure 2. Autocorrelation Function of the of the z-scores of the series of  microstate durations**. The red lines correspond to the FDR p critical values for significant lag dependencies. The red cases represents those bins with a serial duration dependency ftomprevious bins.

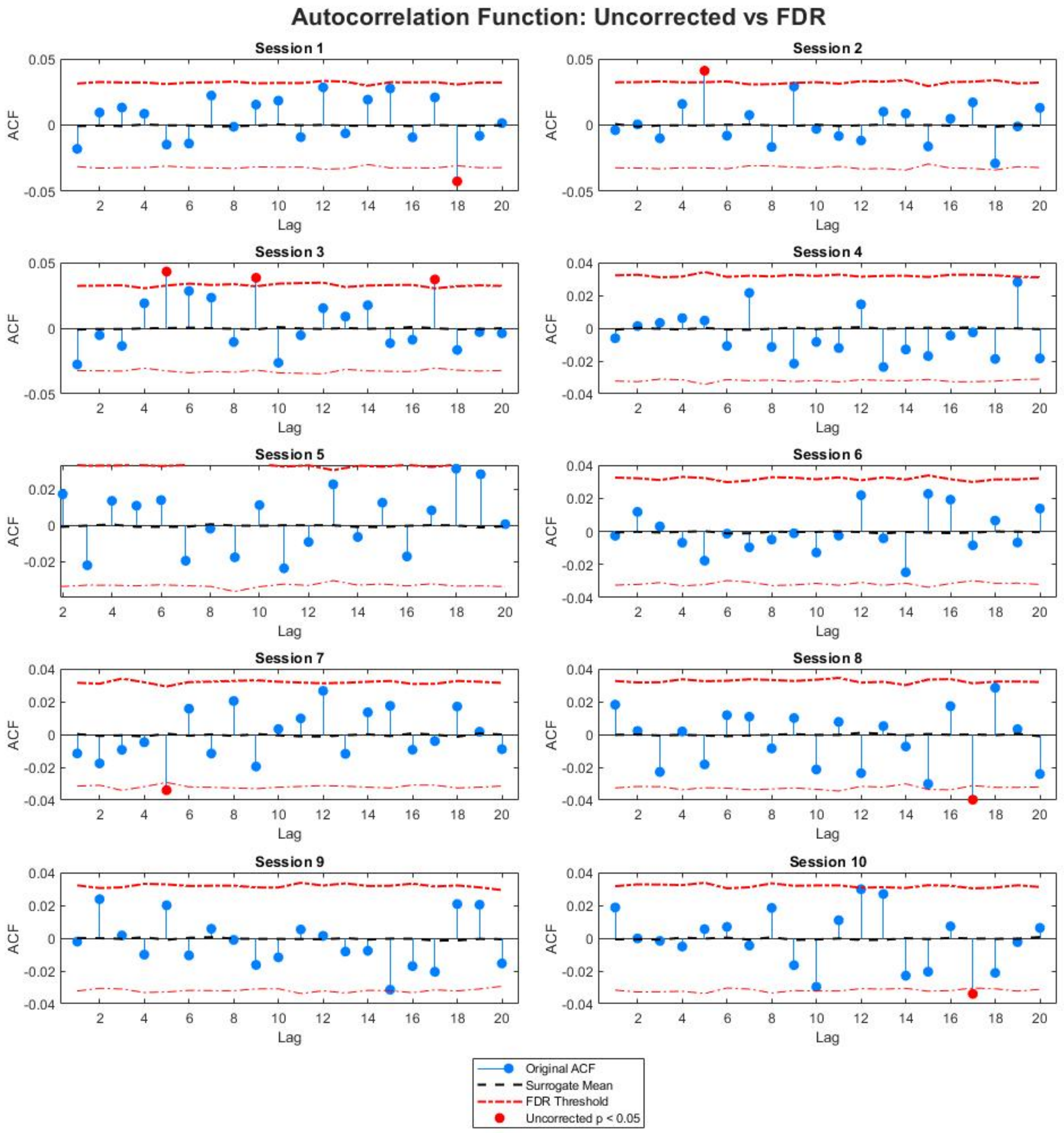

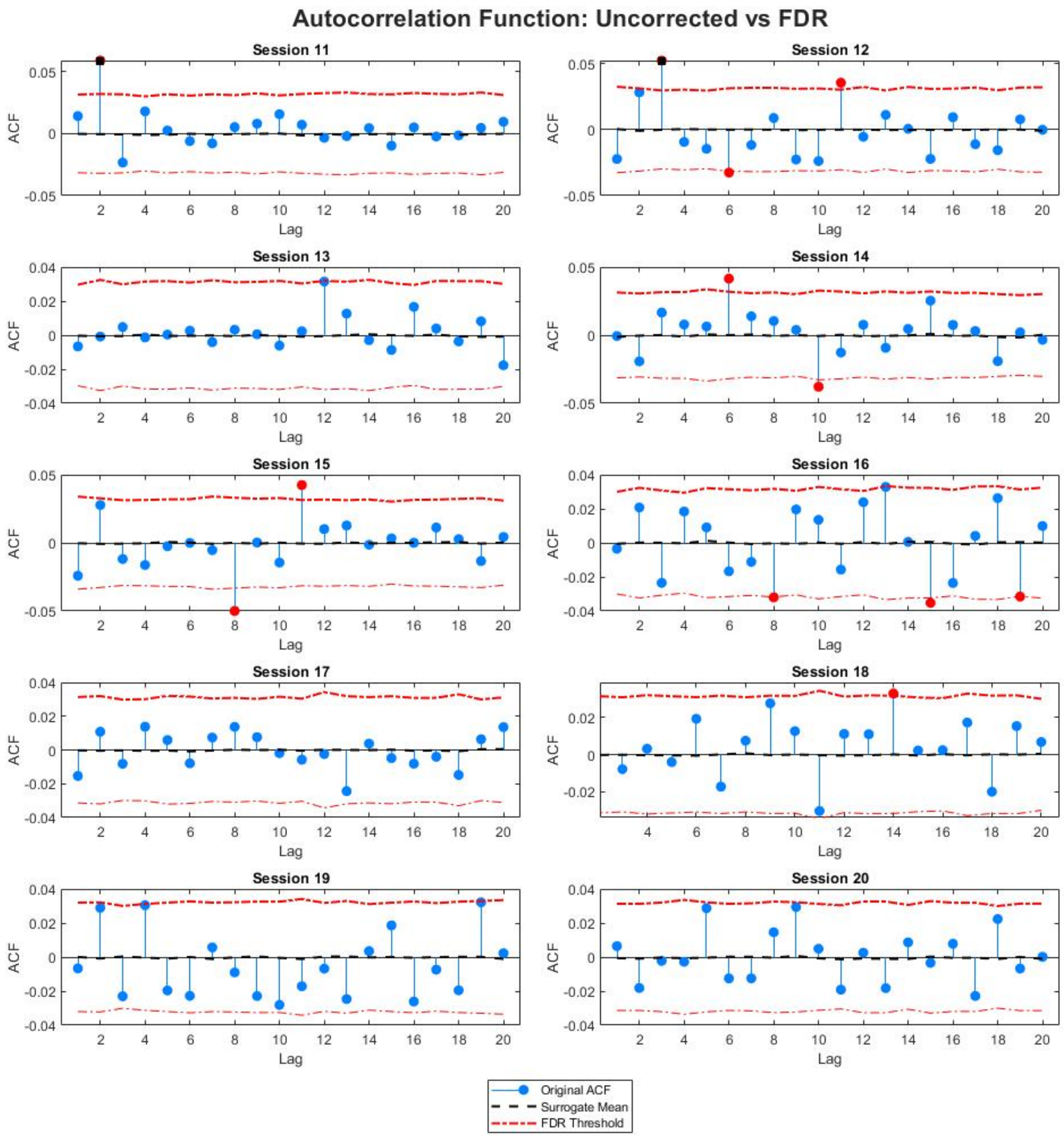

Autocorrelation Function: Uncorrected vs FDR
Session 11
Session 12
Session 13
Session 14
Session 15
Session 16
Session 17
Session 18
Session 19
Session 20
ACF
Lag
Original ACF
Surrogate Mean
FDR Threshold

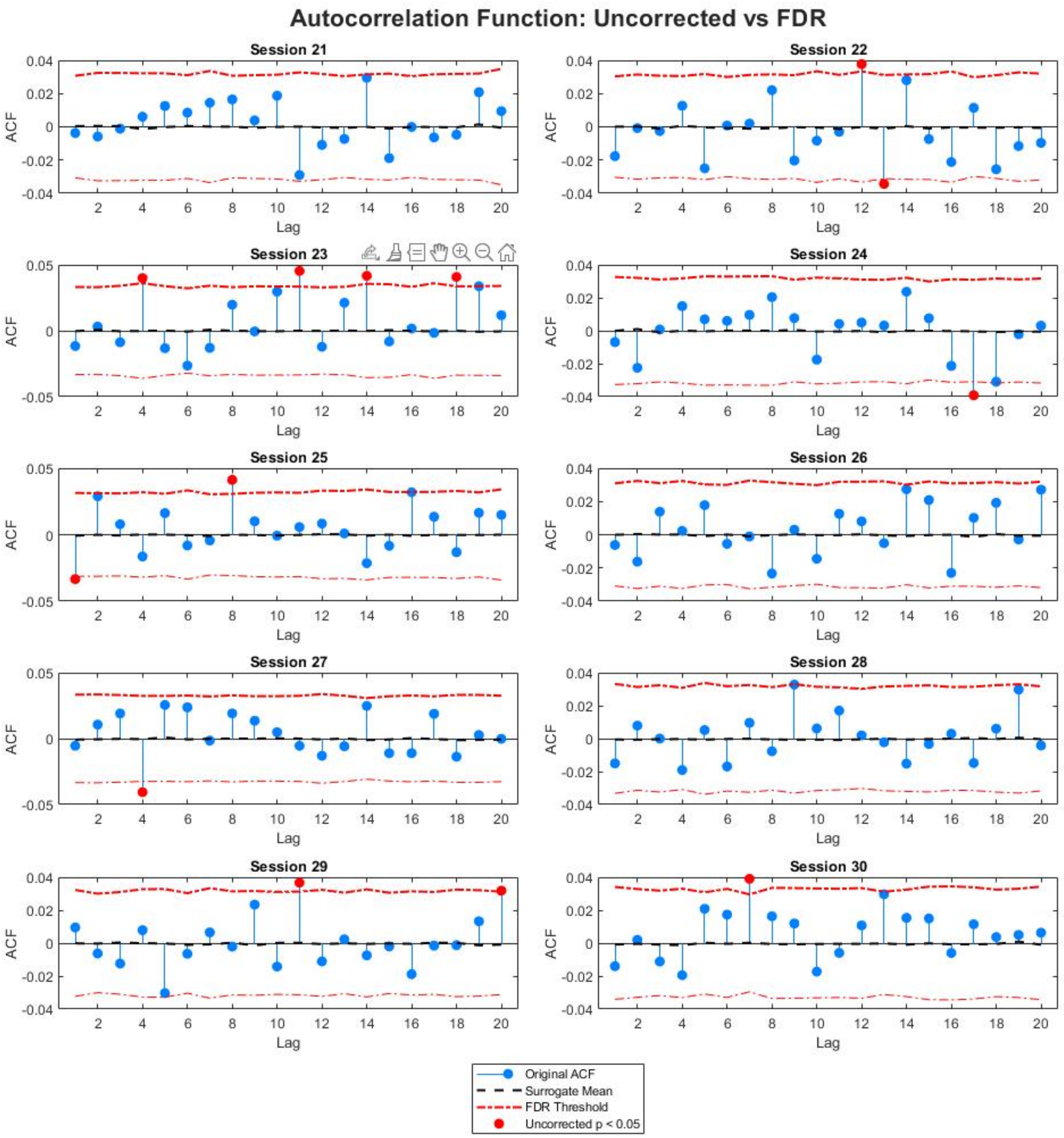
Autocorrelation Function: Uncorrected vs FDR
Session 21
Session 22
Session 23
Session 24
Session 25
Session 26
Session 27
Session 28
Session 29
Session 30
ACF
Lag
Original ACF
Surrogate Mean
FDR Threshold
Uncorrected p < 0.05

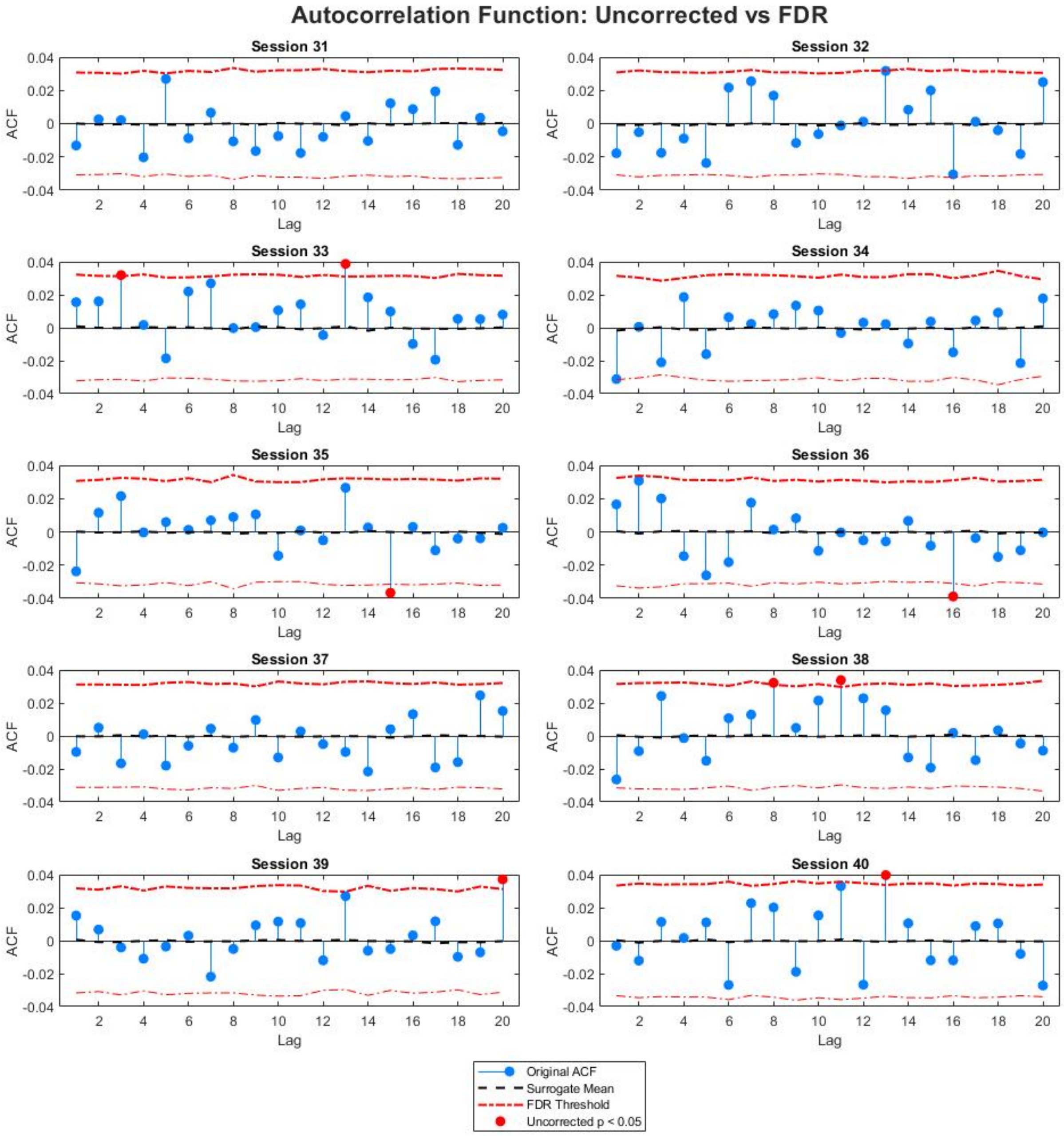

Autocorrelation Function: Uncorrected vs FDR
Session 31
Session 32
Session 33
Session 34
Session 35
Session 36
Session 37
Session 38
Session 39
Session 40
ACF
Lag
Original ACF
Surrogate Mean
FDR Threshold
Uncorrected p < 0.05

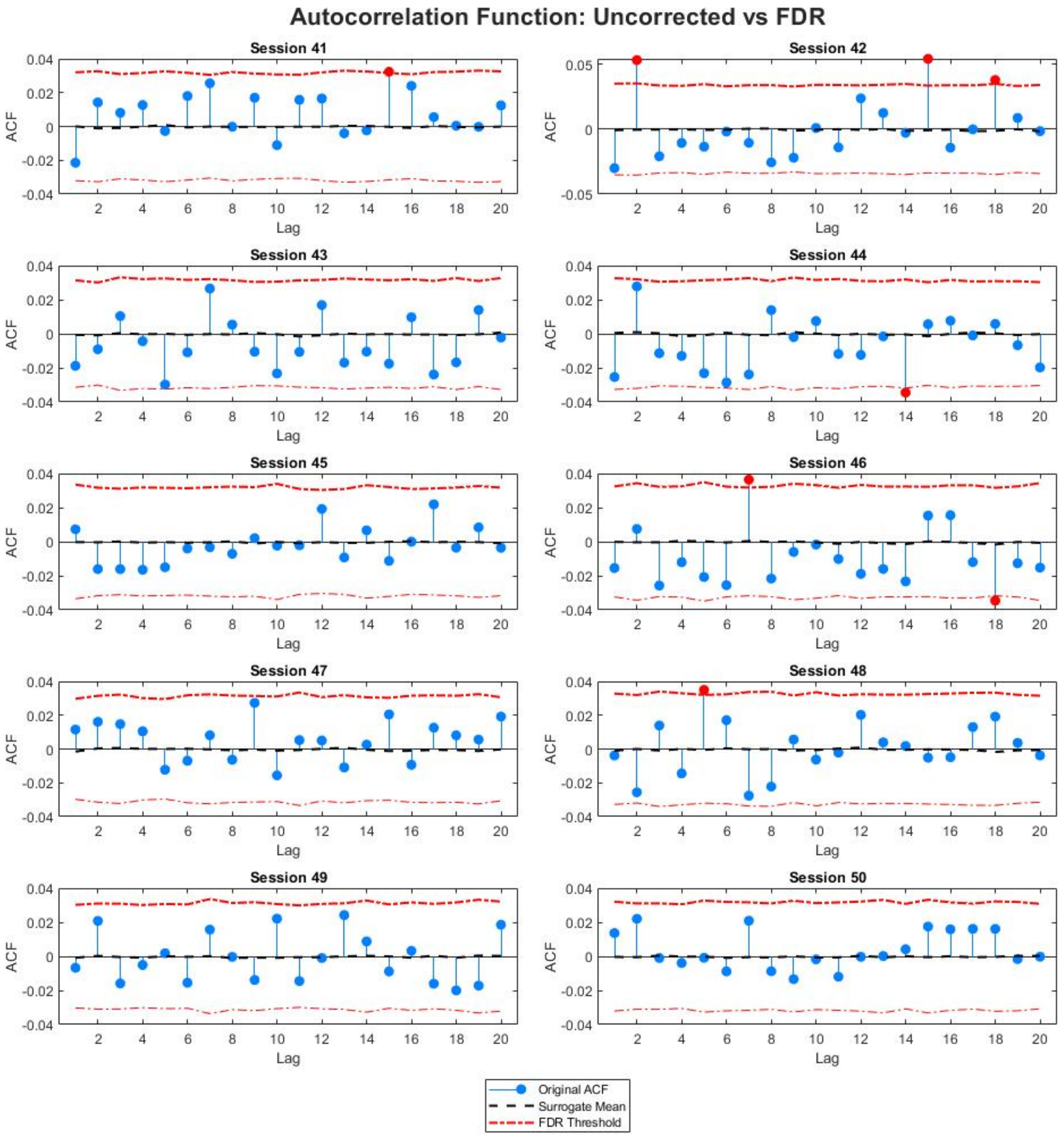
Autocorrelation Function: Uncorrected vs FDR
Session 41
Session 42
Session 43
Session 44
Session 45
Session 46
Session 47
Session 48
Session 49
Session 50
ACF
Lag
Original ACF
Surrogate Mean
FDR Threshold

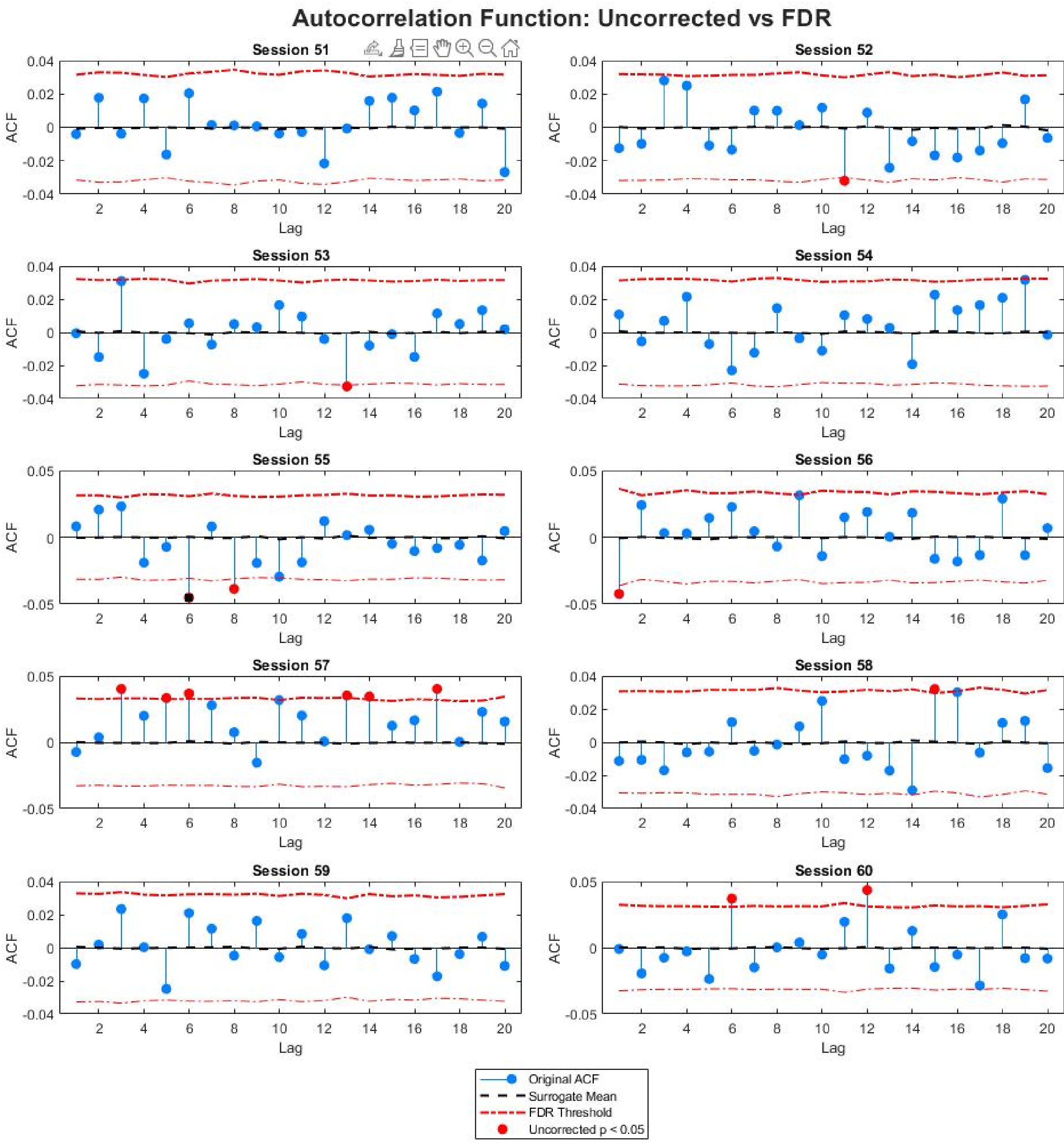

Autocorrelation Function: Uncorrected vs FDR
Session 51
Session 52
Session 53
Session 54
Session 55
Session 56
Session 57
Session 58
Session 59
Session 60
ACF
Lag
Original ACF
Surrogate Mean
FDR Threshold
Uncorrected p < 0.05

**Supplementary Table 1. Random and non-random microstate sequential dependency.** Chi-squared p-values for transitions of each microstate with respect to the other three (columns 2–61). Column 1 indicates the number of sessions in which transitions between microstates are non-random (p > 0.05).

| A | B | C | D |
|---|---|---|---|
| 16.0000 | 31.0000 | 27.0000 | 26.0000 |
| 0.6583 | 0.0012 | 0.0000 | 0.0058 |
| 0.3876 | 0.0590 | 0.0002 | 0.0402 |
| 0.0357 | 0.0000 | 0.0439 | 0.1050 |
| 0.0747 | 0.0515 | 0.0006 | 0.0524 |
| 0.1617 | 0.0000 | 0.3365 | 0.2870 |
| 0.5032 | 0.0196 | 0.2666 | 0.2570 |
| 0.4543 | 0.2165 | 0.6504 | 0.0722 |
| 0.0000 | 0.0798 | 0.0493 | 0.0004 |
| 0.7483 | 0.8595 | 0.8366 | 0.0881 |
| 0.0091 | 0.0000 | 0.0738 | 0.6193 |
| 0.2766 | 0.5053 | 0.1554 | 0.3175 |
| 0.2508 | 0.0665 | 0.4622 | 0.6056 |
| 0.3247 | 0.0213 | 0.0625 | 0.0507 |
| 0.4659 | 0.3275 | 0.0189 | 0.0662 |
| 0.2916 | 0.0780 | 0.0000 | 0.0010 |
| 0.0443 | 0.0085 | 0.8190 | 0.0055 |
| 0.5172 | 0.2872 | 0.1495 | 0.3841 |
| 0.0119 | 0.7440 | 0.0006 | 0.0230 |
| 0.7972 | 0.0370 | 0.0000 | 0.5826 |
| 0.0055 | 0.0440 | 0.2244 | 0.0134 |
| 0.0062 | 0.0480 | 0.0179 | 0.0022 |
| 0.0547 | 0.1985 | 0.0526 | 0.0855 |
| 0.2902 | 0.0000 | 0.0002 | 0.0046 |
| 0.2484 | 0.0771 | 0.4483 | 0.0753 |
| 0.1540 | 0.0101 | 0.7475 | 0.3143 |
| 0.0023 | 0.0351 | 0.1360 | 0.0820 |
| 0.0476 | 0.0001 | 0.0000 | 0.0007 |
| 0.0157 | 0.0015 | 0.1003 | 0.0115 |
| 0.0148 | 0.6254 | 0.0132 | 0.5720 |
| 0.0016 | 0.0000 | 0.0000 | 0.0000 |
| 0.9618 | 0.0014 | 0.0001 | 0.0263 |
| 0.2096 | 0.5244 | 0.1729 | 0.7202 |

| | | | |
|---|---|---|---|
| 0.4361 | 0.0001 | 0.4050 | 0.2511 |
| 0.7940 | 0.0418 | 0.0060 | 0.0967 |
| 0.4748 | 0.5901 | 0.4555 | 0.0978 |
| 0.7140 | 0.0778 | 0.2132 | 0.7139 |
| 0.0691 | 0.1958 | 0.0646 | 0.1640 |
| 0.4590 | 0.0148 | 0.1219 | 0.0029 |
| 0.5577 | 0.0001 | 0.0117 | 0.4095 |
| 0.7714 | 0.0000 | 0.0000 | 0.0000 |
| 0.1760 | 0.0126 | 0.0687 | 0.6041 |
| 0.0252 | 0.0000 | 0.0000 | 0.0000 |
| 0.0032 | 0.1228 | 0.0337 | 0.4046 |
| 0.6174 | 0.3909 | 0.8068 | 0.3583 |
| 0.5406 | 0.0288 | 0.0007 | 0.0014 |
| 0.8405 | 0.0000 | 0.0000 | 0.0000 |
| 0.2393 | 0.1683 | 0.4385 | 0.0470 |
| 0.0093 | 0.0001 | 0.0000 | 0.0009 |
| 0.3993 | 0.2543 | 0.2974 | 0.2269 |
| 0.3485 | 0.3420 | 0.8530 | 0.1631 |
| 0.2095 | 0.1972 | 0.0000 | 0.0000 |
| 0.2061 | 0.5267 | 0.0782 | 0.0250 |
| 0.1806 | 0.5580 | 0.5130 | 0.7657 |
| 0.7401 | 0.0249 | 0.0173 | 0.0046 |
| 0.4559 | 0.0490 | 0.2671 | 0.6598 |
| 0.0412 | 0.0000 | 0.0000 | 0.0000 |
| 0.9115 | 0.0000 | 0.0000 | 0.0000 |
| 0.9340 | 0.0606 | 0.0691 | 0.4805 |
| 0.6602 | 0.8708 | 0.3101 | 0.2407 |
| 0.2049 | 0.1023 | 0.7772 | 0.0120 |

***Access to the data***

The data for the analysis here described are obtained from the article of **Liu et al., 2024**, accessible at https://doi.org/10.6084/m9.figshare.24877770.v3 . The scripts used data in the ALLEEG structure, which are in the file modkmeans_result.mat accessible in the already indicated URL. The data analysis scripts below enclosed , access directly the field ALLEEG(session).microstate.fit.labels once in the workspace.

**Scripts**

**1.Script for figure 1. Definition of the competition model**

```
% geometric computation

close all
clear all
figure('Color', 'w');



for p=0.25:0.1:0.6

nf=100;
j=1
for t=1:0.1:10
f(j)=((1-p)^(t-1))*p;
j=j+1
end
tt=[0:5:450];
a=sum(f);
% normalization of the histogram to make to integral of f1=1
f1=f/a;
b=sum(f1)
f1=f1*nf
% Plot the first subplot
subplot(2, 2, 1);
plot(tt,f1);
hold on
title('A. Expected histograms of microstates following \newline a geometric distibution (eq. 1), changing p');
xlabel('Microstate durations (ms)');
ylabel('N of microstates');
end
hold off
```

```
%c=sum(f1);

%nf=100;

% Dependency of the microstate network probability to win competition from the time elapsed from previous microstate, to make the sigmoid asyntotically approach to the value (p)
p=0.6;
A=5;
j=1;
for t=1:0.1:10
t1=(-1)*t;
f2(j)=(1/(1+(exp((A*t1)+exp(2)))));
j=j+1
end

f2=f2*(p);
f3=1-f2;

subplot(2, 2, 2);
plot(tt,f3);
hold on
plot(tt,f2);

title('B. Probability 1-p`(t) of one microstate to have higher activity \newline than any other microstate p`(t) (eq.2)');
xlabel('time from previous microstate transition (ms)');
ylabel('p`(t) and 1-p`(t)' );
legend('microstate network 1-p`(t)','alternative microstate networks p`(t)', 'location','best')
```

```
% Complete competition model changing A parameter keeping p
fila=1;

p=0.1
for A=2:1:5
j=1
for t=1:0.2:10
t1=(-1)*t;
f2(fila,j)=(1/(1+(exp((A*t1)+exp(2)))));
j=j+1
end
fila=fila+1;
end
f2=f2*(p);

for fila=1:1:4
j=1
for t=1:1:46
f4(fila,t)=((1-f2(fila,t))^(t-1))*f2(fila,t);
end
end

a=sum(f4,2);
b=repmat(a,1,46);

% normalization of the histogram to make to integral of f1=1
f5=f4./b;

nf=100;
f6=f5*nf;
% computing the histogram of duration times
```

```
ttt=[0:10:450];

subplot(2, 2, 3);

plot(ttt,f6);

title('C. Expected histograms of microstate durations \newline changing
parameter A (eq. 2))');

xlabel('Microstate duration (ms)');

ylabel('N of microstates');







% Complete competition model changing p parameter keeping A

fila=1;

A=3;


for p=0.05:0.04:0.23

j=1

for t=1:0.2:10

t1=(-1)*t;

f7(fila,j)=(1/(1+(exp((A*t1)+exp(2)))))*p;

ttt(j)=j;

j=j+1

end

fila=fila+1;

end


for fila=1:1:4

j=1

for t=1:1:46

f8(fila,t)=((1-f7(fila,t))^(t-1))*f7(fila,t);

ttt(j)=j;
```

```
end
end

a=sum(f8,2);
b=repmat(a,1,46);

% normalization of the histogram to make to integral of f1=1
f9=f8./b;

nf=100;
f10=f9*nf;
% computing the histogram of duuration times
ttt=[0:10:450];
subplot(2, 2, 4);
plot(ttt,f10);
title(['D. Expected histograms of microstates \newline changing parameter p (eq. 2)']);
xlabel('Microstate durations (ms)');
ylabel('N of microstates');
```

#### 2.Script for adjusting the c-model to the frequency histograms of the microstated duration

```
%%%%%%%%%%%%%%%%%%%%%%%%%%%%%%%%%%%%%%%%%%%%%%%%%%%%%%%%%%%%%%%%%%%%%%
% fitting_competition_model
% Computes fitting of microstates A,B,C,D across sessions
% Requires ALLEEG in workspace
%%%%%%%%%%%%%%%%%%%%%%%%%%%%%%%%%%%%%%%%%%%%%%%%%%%%%%%%%%%%%%%%%%%%%%
close all
clearvars -except ALLEEG
clc

nSessions = 60;
```

```
nStates = 4;

%% ------------------------------------------------
% STORAGE MATRICES
%% ------------------------------------------------

correlacion_mat = zeros(nSessions,nStates);
A_out_mat = zeros(nSessions,nStates);
p_out_mat = zeros(nSessions,nStates);
ks2stat_mat = zeros(nSessions,nStates);
p_kolm_mat = zeros(nSessions,nStates);
h_mat = zeros(nSessions,nStates);
R_squared = zeros(nSessions,nStates);

f4_store = cell(nSessions,nStates);
hist_store = cell(nSessions,nStates);
nTrimmedZeros = zeros(nSessions,nStates);

%% ------------------------------------------------
% LOOP THROUGH SESSIONS
%% ------------------------------------------------

for session = 1:nSessions

x = ALLEEG(session).microstate.fit.labels;
x = x(:);

changeIdx = [1; find(diff(x) ~= 0) + 1; numel(x) + 1];
runLengths = diff(changeIdx);
runValues = x(changeIdx(1:end-1));

for s = 1:4
```

```
lengths = runLengths(runValues == s);

edges = 0:max(lengths);
counts = histcounts(lengths, edges);

% Trim leading zeros but keep one
firstNonZero = find(counts > 0, 1, 'first');

if ~isempty(firstNonZero) && firstNonZero > 1
nTrimmedZeros(session,s) = firstNonZero - 1;
counts = [0 counts(firstNonZero:end)];
end

%% RUN MODEL
[f4, correlacion, tj, A_out, p_out, h, p_kolm, ...
ks2stat, f2_max, hist_values_mod, f4_mod] = comp_model(counts);

%% STORE
correlacion_mat(session,s) = correlacion;
A_out_mat(session,s) = A_out;
p_out_mat(session,s) = p_out;
ks2stat_mat(session,s) = ks2stat;
p_kolm_mat(session,s) = p_kolm;
h_mat(session,s) = h;

f4_store{session,s} = f4;
hist_store{session,s} = counts;

end
end
```

```
R_squared = correlacion_mat.^2;

%% ------------------------------------------------
% PLOT RESULTS FOR ALL SESSIONS (WITH STATS)
%% ------------------------------------------------

for session_plot = 1:nSessions

figure('Name', ['Session ' num2str(session_plot)], 'NumberTitle',
'off','Color', 'w');

for s = 1:4

subplot(2,2,s)

hist_vals = hist_store{session_plot,s};
f4_vals = f4_store{session_plot,s};

% Plot data
plot(hist_vals,'b','LineWidth',2)
hold on
plot(f4_vals,'r','LineWidth',2)

title(['Microstate ' char('A'+s-1)])
xlabel('ms [(bin*2)+30]')
ylabel('N of microstates')
grid off

% Legend
lgd = legend('Histogram','Model fit','Location','northeast');

% ---- STATS ----
p_val = p_kolm_mat(session_plot,s);
```

```
R2 = R_squared(session_plot,s);

stats_txt = sprintf('p = %.3g\nR^2 = %.3f', p_val, R2);

% Place text BELOW legend (no overlap)

lgd = legend('Histogram','Model fit','Location','northeast');

text(0.90, 0.45, stats_txt, ...
'Units','normalized', ...
'HorizontalAlignment','right', ...
'VerticalAlignment','top', ...
'FontSize',9, ...
'BackgroundColor','w');

end
sgtitle(['Session ' num2str(session_plot)])

end

%% -------------------------------------------------
% SECOND FIGURE (UNCHANGED)
%% -------------------------------------------------

figure ('Color', 'w');

for session_plot = 1:nSessions

for s = 1:4

subplot(2,2,s)
```

```
f4_vals = f4_store{session_plot,s};

plot(f4_vals,'r','LineWidth',0.5)
hold on

title(['Microstate ' char('A'+s-1)])
xlabel ('ms [(bin*2)+30]')
ylabel('N of microstates')
legend('Model fit')
grid off

end
end
```

##### 3. *Function needed for running script 2*

```
function [f4, correlacion, tj, A_out, p_out, h, p_kolm, ...
ks2stat, f2_max, hist_values_mod, f4_mod] = comp_model(counts_C)
hist_values=counts_C;
% This function returns the sum and product of two numbers
%pruebas
%data=[1:1:30];
%p=0.2;
%A=0.3;
%%%%%%%%%%%
clear A_out p_out A p f f1 f2 f2_max f3 f4 correlacion_1 sum_data ...
size_dat size_data prueba hist_values_mod f4_mod
sum_data=sum(hist_values);
size_dat=size(hist_values);
size_data=size_dat(1,2);
hist_values=hist_values';
contador=1;
%compute the best p and A parameters
```

```
%close all

correlacion=0;

tj=1:1:size_data;

for p=0.01:0.01:0.99

for A=0.1:0.1:4.9

for t=1:1:size_data;

f(t,1)=((1-p)^(t-1))*p;

t1=(-1)*t;

f1(t,1)=1/(1+(exp((A*t1)+exp(2))));

f2=f1*(p);

%f3=1-f3;

f3(t,1)=((1-f2(t))^(t-1))*f2(t);

end

prob_sum=sum(f3);

norm1=1/prob_sum;

f3=f3*norm1;

correlacion_1=corr (hist_values,f3);

%correlacion_1

if correlacion_1 > correlacion

correlacion_out=correlacion_1;

correlacion=correlacion_out;

A_out=A;

p_out=p;

maximo=contador;

end

contador=contador+1;

prueba(contador,:)=f3;

prueba_1(contador,:)=f2;

end

end

f4=prueba(maximo,:)*sum_data;

f2_max=prueba_1(maximo,:);
```

```
%[h,p_kolm,ks2stat] = kstest2(hist_values,f4);
% same but collapsing values higher than 750 ms
hist_values=hist_values';
hist_values_mod(1,16)=0;
hist_values_mod(1,1:15)=hist_values(1,1:15);
hist_values_mod(1,16)=sum(hist_values(1,16:29));
f4_mod(1,1:16)=zeros;
f4_mod(1,1:15)=f4(1,1:15);
f4_mod(1,16)=sum(f4(1,16:29));
[h,p_kolm,ks2stat] = kstest2(hist_values_mod,f4_mod);
end
```

*4.* **Script to compute chi-squared t-test to determine if transitions occurs randomly or non-randomly (Table 2; Table 1 exceeds the object of resent report)**

```
clearvars -except ALLEEG
clc

%% ================= PARAMETERS =================
nSessions = 60;
nStates = 4;
nTrans = nStates * (nStates - 1);

%% ================= PREALLOCATION =================
ObsMat = zeros(nSessions,nTrans);
ExpMat = zeros(nSessions,nTrans);
ZMat = zeros(nSessions,nTrans);
PtransMat = zeros(nSessions,nTrans);

Chi2Mat = zeros(nSessions,nStates);
Chi2PMat = zeros(nSessions,nStates);
```

```matlab
%% ================= MAIN LOOP =================
for session = 1:nSessions
x = ALLEEG(session).microstate.fit.labels(:);
changeIdx = [1; find(diff(x) ~= 0) + 1];
data = x(changeIdx);
from = data(1:end-1);
to = data(2:end);
T = accumarray([from,to],1,[nStates,nStates]);
T(1:nStates+1:end) = 0;
stateCounts = histcounts(data,1:nStates+1)';
p_state = stateCounts / sum(stateCounts);
E = zeros(nStates,nStates);
for i = 1:nStates
N_i = sum(T(i,:));
if N_i == 0
continue
end
p_excl = p_state;
p_excl(i) = 0;
p_excl = p_excl / sum(p_excl);
E(i,:) = N_i * p_excl;
end
mask = ~eye(nStates);
ObsVec = T(mask)';
ExpVec = E(mask)';
ZVec = (ObsVec - ExpVec) ./ sqrt(ExpVec);
ZVec(ExpVec==0) = NaN;
PtransVec = 2*(1 - normcdf(abs(ZVec)));
ObsMat(session,:) = ObsVec;
ExpMat(session,:) = ExpVec;
ZMat(session,:) = ZVec;
PtransMat(session,:) = PtransVec;
```

```
for i = 1:nStates
Orow = T(i,:);
Erow = E(i,:);
Orow(i) = [];
Erow(i) = [];
valid = Erow > 0;
if sum(valid) < 2
Chi2Mat(session,i) = NaN;
Chi2PMat(session,i) = NaN;
continue
end
chi2 = sum((Orow(valid) - Erow(valid)).^2 ./ Erow(valid));
df = sum(valid) - 1;
p_chi2 = 1 - chi2cdf(chi2,df);
Chi2Mat(session,i) = chi2;
Chi2PMat(session,i) = p_chi2;
end
end

%% ================= COUNTS =================
for i = 1:nSessions
NumSigTransitions(i,1) = sum(PtransMat(i,:) < 0.05);
NumSigChi2PMat(i,1) = sum(Chi2PMat(i,:) < 0.05);
end

for j = 1:12
NumSigTransitions_columns(1,j) = sum(PtransMat(:,j) < 0.05);
end

for j = 1:4
NumSigChi2PMat_columns(1,j) = sum(Chi2PMat(:,j) < 0.05);
end
```

```
%% ================= TABLES ONLY =================

Table1 = array2table([NumSigTransitions_columns; PtransMat], ...

'VariableNames', {'A→B','A→C','A→D','B→A','B→C','B→D', ...

'C→A','C→B','C→D','D→A','D→B','D→C'});


Table2 = array2table([NumSigChi2PMat_columns; Chi2PMat], ...

'VariableNames', {'A','B','C','D'});


%% ================= OUTPUT =================

Table1

Table2
```

**5. Script to compute the Autocorrelation function. In** `LOOP OVER DATASETS` change step 1:10; 11:20....51:60

```
% ============================================================

% COMBINED AUTOCORRELATION FIGURE (5x2)

% WITH SURROGATES + UNCORRECTED + FDR VISUALIZATION

% ============================================================


close all

clearvars -except ALLEEG

clc


% ----------------------------

% Parameters

% ----------------------------

maxLag = 20;

nSurrogates = 1000;

alpha = 0.05;


% Preallocate storage

total_acfVals = zeros(maxLag,10);
```

```matlab
total_pVals = zeros(maxLag,10);
total_sigFDR = false(maxLag,10);
total_sigUNC = false(maxLag,10);

% ----------------------------
% Figure
% ----------------------------
fig = figure('Color','w','Position',[100 50 1000 1200]);
t = tiledlayout(5,2,'Padding','compact','TileSpacing','compact');

% ============================================================
% LOOP OVER DATASETS
% ============================================================
for n = 1:10

session = n;

% ----------------------------
% 1. Load data
% ----------------------------
x = ALLEEG(session).microstate.fit.labels;
x = x(:);

% ----------------------------
% 2. Run-length encoding
% ----------------------------
changeIdx = [1; find(diff(x) ~= 0) + 1; numel(x) + 1];
runLengths = diff(changeIdx);
runValues = x(changeIdx(1:end-1));

runLengths = runLengths(:);
runValues = runValues(:);
```

```
% ----------------------------
% 3. Z-score per subtype
% ----------------------------
z_runLengths = zeros(size(runLengths));
subtypes = unique(runValues);

for s = subtypes'
idx = (runValues == s);
data = runLengths(idx);
data = data(:);

if numel(data) > 1 && std(data) > 0
z = (data - mean(data)) ./ std(data);
else
z = zeros(size(data));
end

z_runLengths(idx) = z;
end

data = z_runLengths;
data = data(:);

% ----------------------------
% 4. Autocorrelation
% ----------------------------
mu = mean(data);
denom = sum((data - mu).^2);

acfVals = zeros(maxLag,1);
```

```
for lag = 1:maxLag

xlag = data(1:end-lag);

ylag = data(1+lag:end);

acfVals(lag) = sum((xlag - mu).*(ylag - mu)) / denom;

end


% ----------------------------

% 5. Surrogates

% ----------------------------

acfSurrogates = zeros(maxLag, nSurrogates);


for s = 1:nSurrogates

data_shuff = data(randperm(length(data)));


mu_shuff = mean(data_shuff);

denom_shuff = sum((data_shuff - mu_shuff).^2);


for lag = 1:maxLag

xlag = data_shuff(1:end-lag);

ylag = data_shuff(1+lag:end);


acfSurrogates(lag,s) = sum((xlag - mu_shuff).*(ylag - mu_shuff)) /
denom_shuff;

end

end


% ----------------------------

% 6. p-values (surrogate test)

% ----------------------------

pVals = zeros(maxLag,1);


for lag = 1:maxLag

pVals(lag) = (1 + sum(abs(acfSurrogates(lag,:)) >= abs(acfVals(lag)))) ...
```

```
/ (nSurrogates + 1);
end

% ----------------------------
% 7. Uncorrected significance
% ----------------------------
sigUNC = pVals < alpha;

% ----------------------------
% 8. FDR (Benjamini-Hochberg)
% ----------------------------
[pSorted, sortIdx] = sort(pVals);
m = maxLag;
threshBH = (1:m)'/m * alpha;

sigSorted = pSorted <= threshBH;
sigFDR = false(maxLag,1);

if any(sigSorted)
maxSig = find(sigSorted,1,'last');
sigFDR(sortIdx(1:maxSig)) = true;
end

% ----------------------------
% 9. FDR threshold envelope
% ----------------------------
pCrit = threshBH(end);

acfThresh = zeros(maxLag,1);
for lag = 1:maxLag
acfThresh(lag) = prctile(abs(acfSurrogates(lag,:)), 100*(1 - pCrit));
end
```

```matlab
% -----------------------------
% 10. Plot
% -----------------------------
ax = nexttile; hold on;

acfMean = mean(acfSurrogates,2);

% Original ACF
stem(1:maxLag, acfVals, 'filled', ...
'Color',[0 0.5 1], ...
'DisplayName','Original ACF');

% Surrogate mean
plot(1:maxLag, acfMean, 'k--', ...
'LineWidth',1.5, ...
'DisplayName','Surrogate Mean');

% FDR threshold
plot(1:maxLag, acfThresh, 'r-.', ...
'LineWidth',1.5, ...
'DisplayName','FDR Threshold');
plot(1:maxLag, -acfThresh, 'r-.', ...
'HandleVisibility','off');

% -----------------------------
% Uncorrected significant (RED)
% -----------------------------
plot(find(sigUNC), acfVals(sigUNC), 'ro', ...
'MarkerFaceColor','r', ...
'DisplayName','Uncorrected p < 0.05');
```

```
% -----------------------------
% FDR significant (BLACK)
% -----------------------------
plot(find(sigFDR), acfVals(sigFDR), 'ks', ...
'MarkerFaceColor','k', ...
'DisplayName','FDR significant');

xlabel('Lag')
ylabel('ACF')
title(sprintf('Session %d', n))
box on

% -----------------------------
% Store results
% -----------------------------
total_acfVals(:,n) = acfVals;
total_pVals(:,n) = pVals;
total_sigFDR(:,n) = sigFDR;
total_sigUNC(:,n) = sigUNC;

end

% -----------------------------
% 11. Legend
% -----------------------------
lg = legend(ax, 'Location','southoutside');
lg.Layout.Tile = 'south';

% -----------------------------
% 12. Title
% -----------------------------
sgtitle('Autocorrelation Function: Uncorrected vs FDR', ...
```

```
'FontSize',16,'FontWeight','bold');
```
